# The genetic landscape of Ethiopia: diversity, intermixing and the association with culture

**DOI:** 10.1101/756536

**Authors:** Saioa López, Ayele Tarekegn, Gavin Band, Lucy van Dorp, Nancy Bird, Sam Morris, Tamiru Oljira, Ephrem Mekonnen, Endashaw Bekele, Roger Blench, Mark G. Thomas, Neil Bradman, Garrett Hellenthal

## Abstract

The rich linguistic, ethnic and cultural diversity of Ethiopia provides an unprecedented opportunity to understand the level to which cultural factors correlate with -- and shape -- genetic structure in human populations. Using primarily novel genetic variation data covering 1,214 Ethiopians representing 68 different ethnic groups, together with information on individuals’ birthplaces, linguistic/religious practices and 31 cultural practices, we disentangle the effects of geographic distance, elevation, and social factors upon shaping the genetic structure of Ethiopians today. We provide evidence of associations between social behaviours and increased genetic differences among present-day peoples. We show that genetic similarity is broadly associated with linguistic classifications, but indicate pronounced genetic similarity among groups from disparate language classifications that may in part be attributable to recent intermixing. We also illustrate how groups reporting the same culture traits are more genetically similar on average and show evidence of recent intermixing, suggesting how shared cultural traits may promote admixture. In addition to providing insights into the genetic structure and history of Ethiopia, these results identify the most important cultural and geographic proxies for genetic differentiation and provide a resource for designing sampling protocols for future genetic studies involving Ethiopians.

## Introduction

Ethiopia is one of the world’s most ethnically and culturally diverse countries, with over 70 different languages spoken across more than 80 distinct ethnicities (www.ethnologue.com). Its geographic position and history (briefly summarised in SI section 1) motivated geneticists to use blood groups and other classical markers to study human genetic variation (Mourant et al., 1974, Harrison, 1976). More recently, the analysis of genomic variation in the peoples of Ethiopia has been used, together with information from other sources, to test hypotheses on possible migration routes at both ‘Out of Africa’ and more recent ‘Migration into Africa’ timescales (Pagani et al., 2015, Gallego-Llorente et al., 2015). The high genetic diversity in Ethiopians facilitates the identification of novel variants, and this has led to the inclusion of Ethiopian data in studies of the genetics of elite athletes (Rankinen et al., 2016, Ash et al., 2011, Scott et al., 2005), adaptation to living at high elevation (Huerta-Sanchez et al., 2013, Stobdan et al., 2015, Simonson, 2015, Ronen et al., 2014, Scheinfeldt et al 2012), milk drinking (Liebert et al., 2017, Jones et al., 2013, Ingram et al., 2009) and drug metabolising enzymes (Creemer et al., 2016, Browning et al., 2010, Sim et al., 2006).

While the relationships of Beta Israel with other Jewish communities have been the subject of focused research following their migration to Israel (Non et al., 2011, Behar et al., 2010, Thomas et al., 2002), studies involving genomic analyses of the history of wider sets of Ethiopian groups have been more limited (Tadesse et al., 2014, Poloni et al., 2009). Although as early as 1988 Cavalli-Sforza et al. (1988) drew attention to the importance of bringing together genetic, archaeological and linguistic data, there have been few attempts to do so in studies of Ethiopia (Boattini et al., 2013, Pagani et al., 2012, Gallego-Llorente et al., 2015, Baker et al 2017, Scheinfeldt et al 2019, Gopalan et al 2019). Generally, studies have been limited to analysing data from single autosomal loci, non-recombining portion of the Y chromosome and mitochondrial DNA (Semino et al., 2002, Kivisild et al., 2004, Poloni et al., 2009, Boattini et al., 2013, Messina et al., 2017) and/or relatively few ethnic groups (Scheinfeldt et al., 2012, Pagani et al., 2012, Pagani et al., 2015, Scheinfeldt et al., 2019, Gopalan et al., 2019, Prendergast et al., 2019), which has severely limited the inferences that can be drawn. Furthermore, hitherto there has been little exploration of how genetic similarity is associated with shared cultural practices (see however van Dorp et al., 2015) despite the considerable variation known to exist in cultural practices, particularly in the southern part of the country (The Council of Nationalities, Southern Nations and Peoples Region, 2017). For example, Ethiopian ethnic groups have a diverse range of religions, social structures and marriage customs, which may impact which groups intermix, and hence provide an on-going case study of socio-cultural selection (see for example Levine (2000), Freeman & Pankhurst (2003), The Council of Nationalities, Southern Nations and Peoples Region, (2017)) that can be explored using DNA.

Here we analyse autosomal genetic variation information at 534,915 single nucleotide polymorphisms (SNPs) in 1,214 Ethiopian individuals that include 1,082 previously unpublished samples and 132 samples from Lazaridis et al., 2014, Gurdasani et al., 2015 and Mallick et al 2016. Our study includes people from 68 distinct self-reported ethnicities (8-73 individuals per ethnic group) that comprise representatives of many of the major language groups spoken in Ethiopia, including Nilo-Saharan (NS) speakers and three branches (Cushitic, Omotic, Semitic) of Afroasiatic (AA) speakers, as well as languages of currently uncertain classification (Chabu, and the speculated, possibly extinct language of the Negede-Woyto) (www.ethnologue.com) (Fig 1a, Fig S1, Extended Table 1, SI Section 2). Each of the 1,082 newly genotyped individuals were selected from a larger collection on the basis that their self-reported ethnicity, and typically birthplace, matched that of their parents, maternal grandmother, paternal grandfather, and any other grandparents recorded, analogous to recent studies of population structure in Europe (Leslie et al., 2015, Byrne et al., 2018). For these individuals we also recorded their reported religious affiliation (four categories), first language (66 total classifications) and/or second language (40 total classifications) (Table S1). Furthermore, some of the authors of this study (A. Tarekegn, N. Bradman) translated into English and edited a compendium (originally published in Amharic) that documented the oral traditions and cultural practices of 56 ethnic groups of the Southern Nations, Nationalities and Peoples’ Region (SNNPR) of Ethiopia through interviews with members of different ethnic groups (The Council of Nationalities, Southern Nations and Peoples Region, 2017). From this new resource, we compiled a list of 31 practices that were reported as cultural descriptors by members of 47 different ethnic groups out of the 68 in this study (see Methods). These practices include self-declared cultural practices such as male and female circumcision, and 29 different pre-marital and marriage customs, including arranged marriages, polygamy, gifts of beads or belts, and covering the bride in butter.

**Figure 1.**
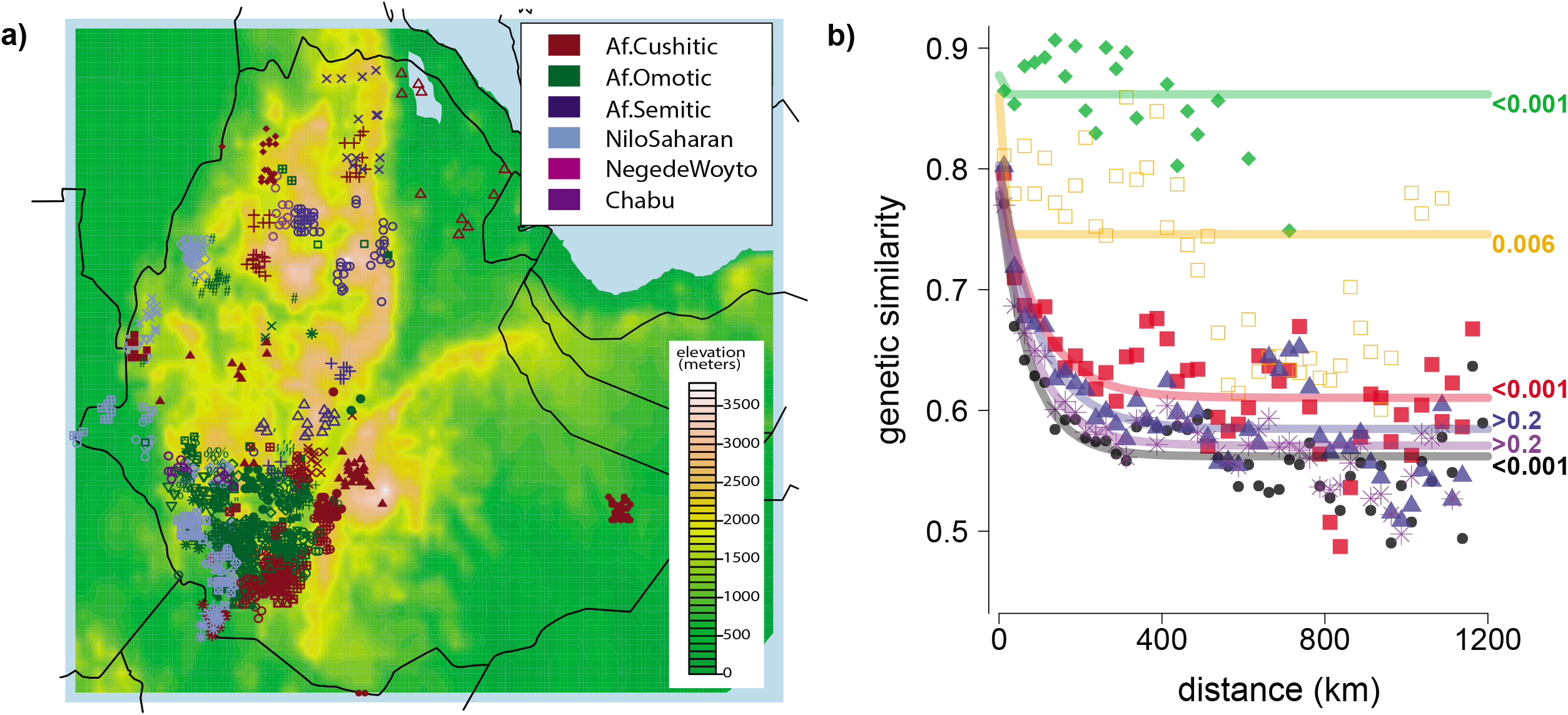
Genetic similarity decays with spatial distance among Ethiopians and correlates with shared reported ethnicity and language. **(a)** Locations of sampled Ethiopians based on birthplace (in some cases slightly moved due to overlap), with landscape colors showing elevation and colored symbols depicting the language category (plus unclassified languages Negede Woyto, Chabu) of each individual’s ethnic identity. The legend for the symbols is provided in Fig S1. **(b)** Fitted model for genetic similarity (1-TVD; under the “Ethiopia-internal” analysis) between pairs of individuals versus geographic distance, with points depicting the average genetic similarity within 25km bins, for all individuals (black; dots) or restricting to individuals who report the same group label (green; diamonds), same first language (orange; open squares), same second language (blue; triangles), same religious affiliation (purple; asterisks), or whose reported ethnicities are from the same language group (red; closed squares). Labels at right give permutation-based p-values when testing the null hypothesis of no increase in genetic similarity among individuals sharing the given trait (see Methods).

We compared SNP patterns in each present-day Ethiopian to those in all other present-day Ethiopians and to the 4,500 year-old Ethiopian sample “Mota”, a forager from southern Ethiopia that represents the only presently available ancient DNA sample from the country (Gallego-Llorente et al., 2015). We also compared them to a further 16 labelled groups comprised of 39 ancient individuals (Hofmanova et al 2016, Mathieson et al 2015, Schlebusch et al 2017, Keller et al 2012, Allentoft et al 2015, Olalde et al 2014, Lipson et al 2020, Prendergast et al 2019, Broushaki et al 2016, Haak et al 2015) and 264 present-day non-Ethiopian groups (Lazaridis et al 2014, Gurdasani et al 2015, Skoglund et al 2017, Mallick et al 2016, Lopez et al 2017, Lipson et al 2020) comprised of 2,678 individuals (average group sample size = 10, range: 1-100), including 106 unpublished samples from nine groups (Table S2). We focus on inferring patterns of haplotype sharing among individuals, which has increased resolution over commonly-used allele-frequency based techniques (Alexander et al., 2009, Price et al., 2006) when identifying latent population structure and inferring the ancestral history of peoples sampled from relatively small geographic regions, such as within a country (Hellenthal et al., 2014, Leslie et al., 2015, van Dorp et al., 2019).

Our results provide a comprehensive understanding of the relative strength to which different socio-cultural factors are associated with genetic distance in present-day Ethiopians. We provide evidence that recent intermixing is increased among groups, sometimes from distantly-related linguistic affiliations, that live nearby and/or share cultural practices. We also provide the inferred recent admixture history for members of 68 ethnic groups.

## Results

### Genetic distance is broadly associated with geography, ethnicity, linguistics and shared culture in Ethiopia

Principal components analysis (PCA) (Price et al 2006, Patterson et al 2006) applied to sampled African individuals revealed Ethiopians to be more genetically similar to each other and sampled groups from other east African countries (Kenya, Somalia, Sudan, Tanzania) than to other African populations (Fig S2b). Runs-of-homozygosity (Chang et al., 2015) and proportions of genome shared identical-by-descent (IBD) (Browning and Browning, 2011) among individuals vary substantially across Ethiopian ethnic groups (Fig S3ab). Ethiopia’s two largest ethnicities, Amhara and Oromo, have the lowest levels of IBD-sharing (Fig S3a), and we observe a significant (p-val<0.001) decrease of homozygosity versus increasing population census size across ethnic groups in the SNNPR (Fig S3c; census from 2007: The Council of Nationalities, Southern Nations and Peoples Region, 2017).

To measure genetic similarity between two Ethiopians, we calculated the total variation distance (TVD) (Leslie et al., 2015) between their haplotype-sharing patterns inferred by CHROMOPAINTER (Lawson et al., 2012) (see Materials and Methods). Mimicking van Dorp et al. (2015), we performed two CHROMOPAINTER analyses in order to infer the broad time periods over which individuals became isolated from one another (see schematic of approach in Fig S4). The first, which we call “Ethiopia-internal,” compares haplotype patterns in each Ethiopian to those in all other sampled individuals. TVD based on this analysis can be thought of as a haplotype-based analogue for commonly-used Fst (Weir & Cockerham, 1984), for which it is correlated in our analyses here (Pearson’s r = 0.63; Mantel-test p-value < 0.00001). However, TVD estimates have been shown to be more powerful at distinguishing subtle genetic differences among e.g. African groups (Busby et al 2016). The second, which we call “Ethiopia-external,” instead compares patterns in each Ethiopian only to those among individuals in non-Ethiopian groups. As the “Ethiopia-internal” analysis compares haplotype patterns in each Ethiopian to those in other Ethiopians, including other members of the same ethnic group, it is more sensitive to detecting endogamy effects and admixture among Ethiopian groups (van Dorp 2015, van Dorp 2019). In contrast, the “Ethiopia-external” analysis mitigates signals related to both of these factors, while remaining sensitive to inferring whether Ethiopians having varying proportions of ancestry related to non-Ethiopian sources due to e.g. different admixture histories (Leslie et al., 2015). We illustrate this in simulations mimicking our real data (Fig S5, SI Section 3).

We first considered how pairwise genetic similarity among Ethiopians is related to several factors. Under both the “Ethiopia-internal” and “Ethiopia-external” analyses, we found significant associations (p-val < 0.05) between genetic distance and each of geographic distance, elevation difference, ethnicity and first language, after controlling each factor for the others where possible (Fig 1b, Fig S6-S7, Tables S3-S7). In contrast, we found no significant association (p-val > 0.2) between genetic distance and each of religion and second language (Fig 1b, Fig S6, Tables S3-S7). However, within six of 16 groups for which we sampled at least five individuals from different religions, we found some nominal evidence (permutation-based p-val < 0.05) of genetic isolation between people reporting as Christians versus those reporting as Muslims or those reporting as practicing traditional religions (Table S8).

We next averaged pairwise genetic similarity values among individuals from the same versus different group labels (Fig S8). Consistent with the relationships depicted by PCA (Fig S2), on average Ethiopian groups are more genetically similar to other Ethiopian groups than they are to the non-Ethiopian groups included in this study (Fig S8, Extended Tables 3-4). We found a significant association between genetic similarity and reporting shared cultural traits among SNNPR groups under the “Ethiopia-internal” analysis (Mantel-test p-value < 0.03; Table S9), which remained after accounting for geographic or elevation distance (partial Mantel-test p-value < 0.05; Table S9) or language group (partial Mantel-test p-value < 0.03; Table S9).

To facilitate comparisons of genetic patterns among groups, we generated an interactive map that graphically displays the genetic similarity among groups under each of the “Ethiopia-internal” and “Ethiopia-external” analyses (https://www.well.ox.ac.uk/~gav/work_in_progress/ethiopia/v5/index.html), with averages summarised in Fig S8 and Extended Tables 3-4. We provide three observations based on these findings. The first observation is that, under the “Ethiopia-internal” analysis, Ari and Wolayta people who work as cultivators or weavers are more genetically similar to members of other ethnicities on average than they are to people from their own ethnicities who work as potters, blacksmiths and tanners (top left squares in Fig 2a). This suggests that occupation is a better proxy for genetic homogeneity than ethnicity in these groups, consistent with caste-like occupational classes observed in these groups (Freeman & Pankhurst, 2003; Todd 1978). Despite this, under the “Ethiopia-external” analysis, Ari and Wolayta are more genetically similar to members of their own ethnicities on average, regardless of occupation (bottom right of squares in Fig 2a). Therefore, contradicting evidence from the “Ethiopia-internal” analysis (Extended Table 3) and Fst (Extended Table 9), the “Ethiopia-external” results suggest that individuals from different occupations within the same ethnic group are more recently related to each other than they are to any other ethnic group.

**Figure 2.**
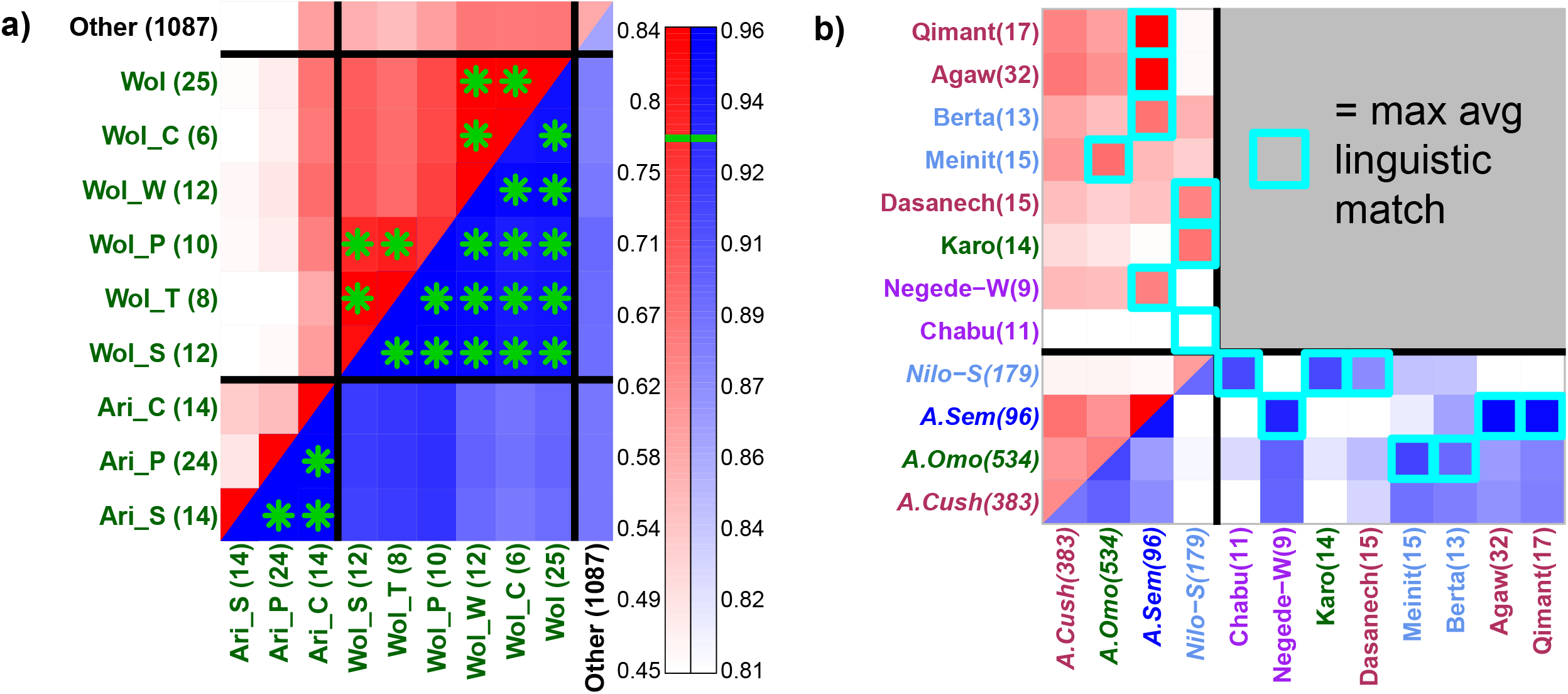
Genetic similarity suggests recent endogamy within caste-like occupational groups and shows shared ancestry among some linguistically divergent groups. Average pairwise genetic similarity (1-TVD) between individuals from different Ethiopian labelled groups (colored on axis by language category -- see Fig 1a), under the “Ethiopia-internal” analysis (top left, red color scale) versus the “Ethiopia-external” analysis (bottom right, blue color scale). **(a)** Genetic similarity between Ari and Wolayta (Wol) caste-like occupational groupings (C = cultivator, P = potter, S = blacksmith, T = tanner, W = weaver), with green asterisks denoting relatively high similarity above the green lines in legend at right. This illustrates how the “Ethiopia-external” analysis shows increased similarity between groups of the same ethnicity relative to that seen under the “Ethiopia-internal” analysis. **(b)** Average pairwise genetic similarity among individuals from four language classifications (italics), and the average genetic similarity between individuals from these four language groups and those from eight ethnic groups (non-italics). For each of the eight ethnic groups, the cyan squares denote the language group with the highest average genetic similarity to that ethnic group under each analysis. This illustrates how the linguistically-unclassified Chabu and Negede-Woyto are most genetically similar on average to Nilo-Saharan and Afroasiatic Semitic speakers, respectively, and highlights six other groups that are more genetically similar to members from a different language group than they are to members of their own language group.

The second example concerns the two sampled groups in our study for which Ethnologue ascribes no linguistic classification, the Chabu and Negede-Woyto. Each are significantly differentiable (p-val < 0.001) from all other ethnic groups under the “Ethiopia-internal” analysis (Fig 2b, Fig S8a, Extended Table 3). The Chabu, a hunter-gatherer group and linguistic isolate, exhibit the strongest overall degree of genetic differentiation from all other ethnic groups, consistent with previous analyses highlighting their genetic isolation (Scheinfeldt et al 2019, Gopalan et al 2019). However, under the “Ethiopia-external” analysis, the Chabu show similar genetic patterns to other NS speaking groups, while the Negede-Woyto are not significantly distinguishable from multiple ethnic groups representing all three AA branches (Fig 2b, Fig S8b, Fig S9). The Chabu’s similarity to NS speakers reflects previous findings from genetics (Scheinfeldt et al., 2019, where the Chabu were referred to as Sabue, Gopalan et al 2019) and linguistics (Blench, 2006; Ehret, 1992). However, we clarify this further by inferring the Chabu to be significantly most genetically similar to the Mezhenger (Fig S8a), with whom they have been suggested to share origins (Dira and Hewlett 2017).

Third, we find unexpectedly close genetic similarities among groups classified into distantly related linguistic categories (Fig 2b, Fig S8). For example, the AA-speaking Karo and Dasanech are on average more genetically similar to NS speakers than to other AA speakers. In contrast, the NS speaking Meinit and Berta are more similar to AA speakers. At a finer linguistic level, the AA Cushistic-speaking Agaw and Qimant are most genetically similar to sampled AA Semitic-speakers, with the Qimant and AA Semitic-speaking Beta Israel having been reported previously to be related linguistically to the Agaw (Appleyard, 1996). These observations demonstrate that shared linguistic affiliation, even using broad categories, is not always a reliable proxy for relatively higher genetic similarity. However, on average individuals from the AA Cushitic, AA Omitic, AA Semitic, and NS classifications, as well as individuals from separate sub-branches within each of these categories, are genetically distinguishable from each other under both the “Ethiopia-internal” and “Ethiopia-external” analyses (p-val < 0.001; SI Section 4; Fig S9; Extended Tables 7-8), consistent with Pagani et al 2012. This suggests that the first three tiers of Ethiopian language classifications at www.ethnologue.com are genetically -- in addition to linguistically -- separable on average, and that these genetic differences are not solely attributable to endogamy effects but also to differential ancestry related to non-Ethiopians. We also find that several groups spanning the three AA classifications of Cushitic, Omotic, Semitic show high genetic similarity to each other on average and less genetic similarity to NS speakers (Fig 2b, Figs S8-S9). We find no clear evidence Omotic are an outgroup to other AA language groups as has been claimed (Baker et al 2017), at least among Ethiopians.

### The recent admixture history of Ethiopia

We attempted to shed light on the ancestry of different Ethiopian groupings by comparing their haplotype sharing patterns under the “Ethiopia-external” analysis to those in a set of reference populations meant to reflect ancestral source populations. To do so, we first used fineSTRUCTURE (Lawson et al., 2012) to assign Ethiopians into 78 clusters of relative genetic homogeneity (Fig S10, Extended Table 2). Unsuprisingly, given genetic similarity observations (Fig 1b, Fig S6, Fig S8), these clusters were notably associated with ethnic label (Fig S10), with clusters inferred using an alternative approach ADMIXTURE (Alexander et al 2009) also often corresponding to a single ethnic group (Fig S11). However, using clusters rather than self-reported label can increase power by merging ethnic groups with similar genetics. This also can clarify ancestry inference, as it does not assume that all individuals reporting the same ethnicity share recent ancestry. We applied SOURCEFIND (Chacón-Duque et al., 2018) and GLOBETROTTER (Hellenthal et al., 2014) to infer and describe admixture events in each of these 78 clusters (see Methods, SI section 5). Simulations mimicking these data showed our approach provides highly accurate inferred sources and dates of admixture (Fig S5d, SI section 3).

GLOBETROTTER infers clear admixture events in 68 of the 77 Ethiopian clusters containing more than one individual, with dates ranging from ∼100 to 4200 years ago (SI section 5, Fig 3, Fig S12, Extended Table 5). Out of 275 reference populations, SOURCEFIND infers only 13 contributed >5% towards describing ancestry patterns within any of these 68 clusters: the 4,500-year-old Ethiopian Mota and 12 present-day groups from Chad, Egypt, Kenya, Saudi Arabia, Somalia, Sudan, Tanzania, Uganda and Yemen (Fig 3, Fig S12, Extended Table 5). Strikingly, matching to Mota shows a significant decrease with increasing spatial (geographic and elevation) distance between the average location of individuals in each cluster and where Mota was discovered (linear regression p-value < 0.0005, Fig S13, Table S10).

**Figure 3.**
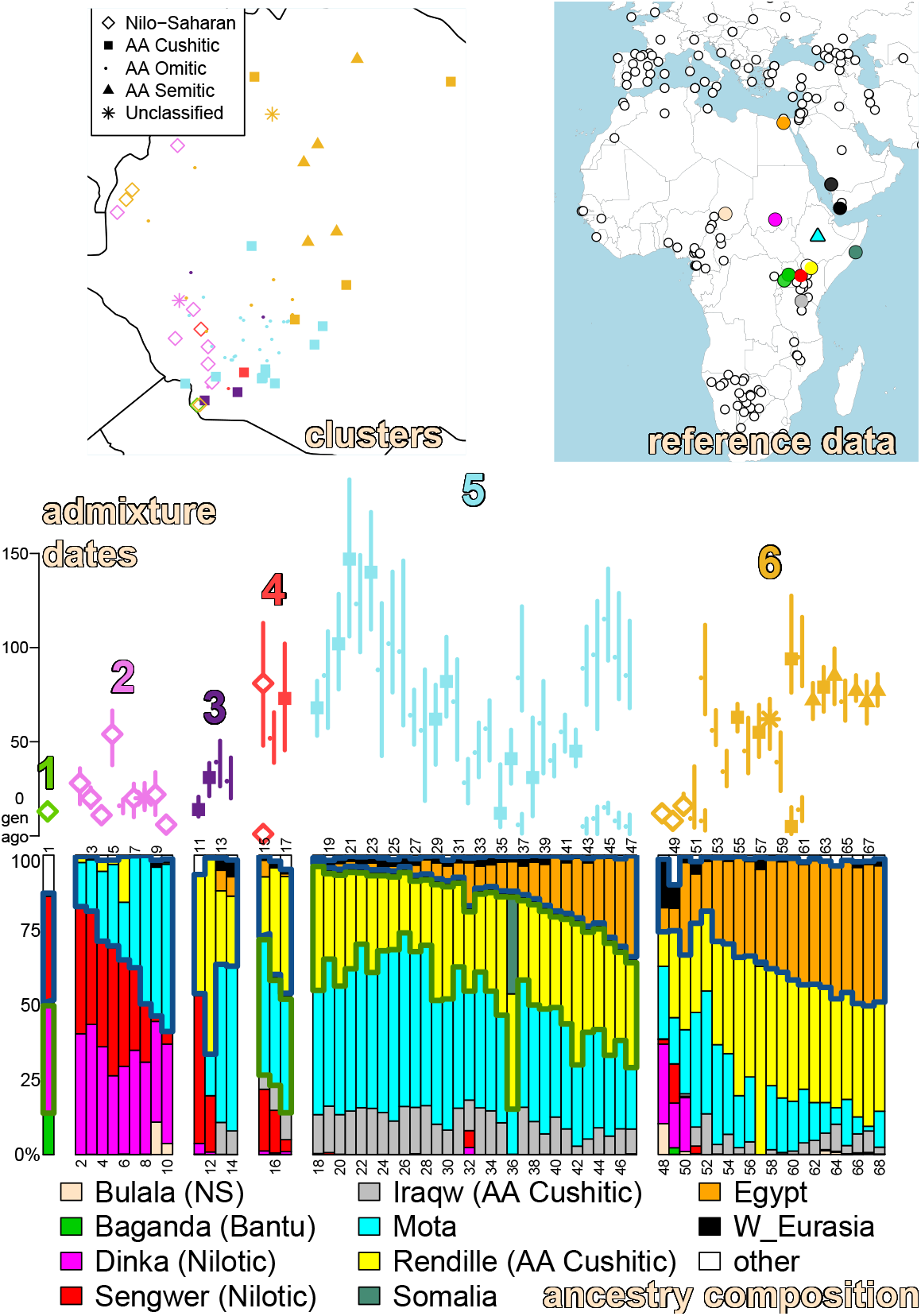
Inferred ancestral composition and recent admixture events in each Ethiopian cluster. **(top-left)** FINESTRUCTURE-inferred genetically homogeneous clusters of Ethiopians, with location placed on the map by averaging the latitude/longitude of each cluster’s individuals. Colors denote which of six types of admixture event (1-6 below) each cluster falls into and symbols provide the most-represented language group among individuals’ ethnicities in each cluster. **(top-right)** A subset of the 264 non-Ethiopian present-day reference populations, plus the 4.5kya Ethiopian Mota (Gallego-Llorente et al., 2015; cyan diamond), that DNA patterns in each Ethiopian cluster were compared to under the “Ethiopian-external” analysis. Filled circles (legend at bottom) indicate reference populations that contributed >5% of ancestry to at least one Ethiopian using SOURCEFIND. **(Middle)** Inferred admixture dates in generations from present (symbols give means and correspond to legend in the top-left panel, line=95% CI), colored by the six types of admixture event. **(Bottom)** SOURCEFIND-inferred ancestry proportions for each Ethiopian cluster (key for numbers in Extended Table 5). Blue and green borders in the ancestry composition highlight different admixing sources. In particular we enclose the reference populations representing one of the inferred admixing sources with a thick blue line. In Ethiopian groups with >2 inferred sources, we also enclose the reference populations representing the second source with a thick green line. Using this information, we highlight six types of inferred admixture events among: (1) three sources related to the Baganda, Dinka and Sengwer, (2) two sources related to Mota and Dinka/Sengwer, (3) two sources related to Rendille and Mota or Sengwer, (4) three sources related to Rendille, Mota and Sengwer, (5) three sources related to Egypt/W.Eurasia, Rendille and Mota/Iraqw, and (6) two sources related to Egypt/W.Eurasia and Rendille/Mota.

We observe six broad categories of admixture, correlated with both geography and linguistics (Fig 3, Fig S12). For example, 12 clusters primarily containing individuals from NS-speaking groups (clusters 1-5, 7, 9, 10, 15, 48-50 on Fig 3, Fig S12) show evidence of admixture involving a source related to Bantu (Baganda) and/or Nilotic (Sengwer, Dinka) speakers, with date estimates <30 generations ago in all but two of these clusters. Similar admixture is inferred in the AA Omotic speaking Karo (cluster 6), AA Cushitic speaking Dasanech (cluster 11) and linguistically-unclassified Chabu (cluster 8) that each show relatively high genetic similarity to NS-speakers (Fig 2A). In contrast, clusters primarily containing AA speakers, including all Ari and Woylata clusters (clusters 22, 24, 25, 39, 41, 43, 45, 54, 56) and a cluster containing the linguistically-unclassified Negede-Woyto (cluster 58), typically show evidence of admixture between two or more sources related to the 4.5kya Ethiopian Mota, Cushitic-speaking Rendille from Kenya and Egypt/W.Eurasian groups, over a broader range of dates (5-147 generations ago). Among these, five northern clusters containing AA Semitic-speakers and the AA Cushitic-speaking Agaw (clusters 62, 64, 66-68), plus two geographically nearby clusters containing the AA Cushitic-speaking Qimant (cluster 63) and AA Omotic-speaking Shinasha (cluster 65), show the highest amounts of Egypt-like ancestry in our dataset and similar admixture dates (point estimates 71-85 generations ago).

### Pervasive recent intermixing among groups is associated with geographic proximity and shared cultural practices

The surprising genetic similarity among dissimilar language groups may be attributable in part to ethnic groups adopting their current languages relatively recently and/or to recent intermixing among distinct Ethiopian groups. To test for the latter, we also applied GLOBETROTTER to each of the 77 Ethiopian clusters under the “Ethiopia-internal” analysis, which includes Ethiopians as surrogates for admixing sources and hence can characterize intermixing that has occurred among Ethiopian groups. GLOBETROTTER found evidence of admixture in 61 clusters in this analysis, 46 (75.4%) of which had estimated dates <30 generations ago (<900 years ago) (Extended Table 6). Across clusters, inferred dates under the “Ethiopia-internal” analysis typically are much more recent than those inferred under the “Ethiopia-external” analysis (Fig 4a). This indicates that the “Ethiopia-internal” analysis is identifying recent intermixing among Ethiopian groups rather than relatively older admixture involving sources related to non-Ethiopians; otherwise dates under the two analyses would be similar. Furthermore, we found that recent intermixing occurs more frequently than expected among clusters whose individuals reside geographically near to each other (p-value < 0.00002, Fig 4b).

**Figure 4.**
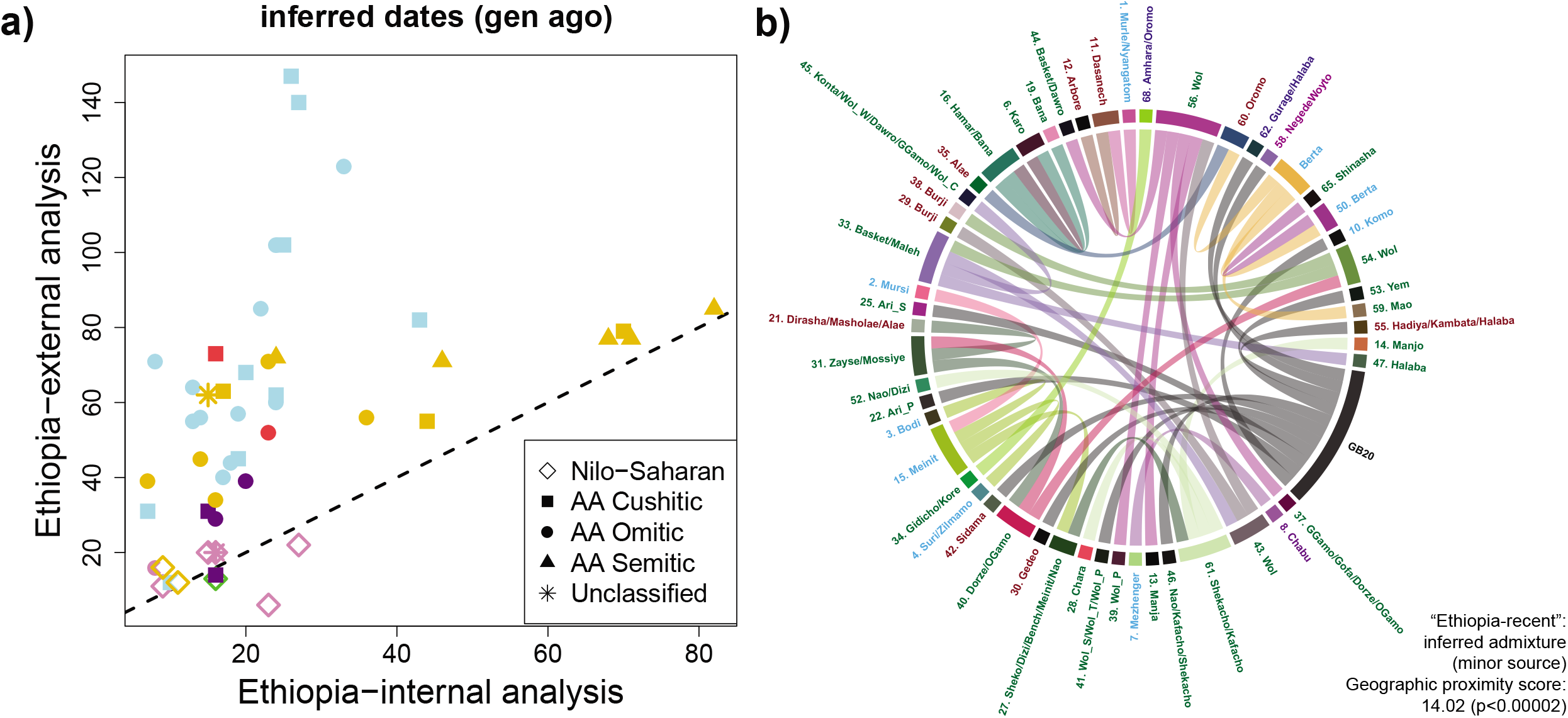
Evidence of recent intermixing among nearby Ethiopian groups. **(a)** GLOBETROTTER-inferred dates (generations from present) for each Ethiopian cluster inferred to have a single date of admixture under each of the “Ethiopia-external” and “Ethiopia-internal” analyses. Inferred dates typically are more recent under the latter, indicating this analysis is picking up relatively more recent intermixing among sources represented by present-day Ethiopian clusters. Colors match those in Fig 3 for these clusters. **(b)** GLOBETROTTER-inferred ancestry sources under the “Ethiopia-internal” analysis. Each Ethiopian cluster *X*, also including Mota, has a corresponding color (outer circle). Lines of this color emerging from *X* indicate that *X* was inferred as the best surrogate for the admixing source contributing the minority of ancestry to each other cluster it connects with. The thickness of lines is proportional to the contributing proportion. Ethiopian clusters, with labels colored by language category according to Fig 1a, are ordered by the first component of a principal components analysis applied to the geographic distance matrix between groups, i.e. so that geographically close groups are next to each other. The “geographic proximity score” gives the average ordinal distance between an admixture target and the surrogate that best represents the source contributing a minority of the admixture, with the p-value testing the null hypothesis that admixture occurs randomly between groups (i.e. independent of the geographic distance between them) based on permuting cluster labels around the circle.

We next explored whether groups that share cultural practices also show evidence of recent intermixing. Supporting this, we found a significant association (p-value < 0.05) between genetic similarity and shared cultural practices only under the “Ethiopia-internal” analysis that is sensitive to intermixing among Ethiopian groups (Table S9). Six traits out of the 20 reported by more than one ethnic group exhibited nominally higher (per-test p-value < 0.05) genetic similarity among ethnic groups participating in the practice relative to those who did not participate or whose participation in the practice was unknown (Fig 5). These practices include male and female circumcision and four different marriage practices (see SI section 6 for details). Groups sharing one of these six traits in common showed higher average genetic similarity than that expected based on linguistic affiliation and spatial distance (Fig 5), and we see increased evidence of recent intermixing among groups reporting male/female circumcision and sororate/cousin marriages relative to other SNNPR groups (Fig 5, Table S11, see Methods, SI section 5). As an example, GLOBETROTTER infers admixture occurring 16 generations ago (95% CI: 12-21) in a cluster of the AA Cushitic-speaking Dasanech (cluster 11 in Fig 4b), from a source most genetically related to a cluster containing the NS-speaking Murle and Nyangatom that share practices of arranged and abduction marriages (Fig 4b, Extended Table 6).

**Figure 5.**
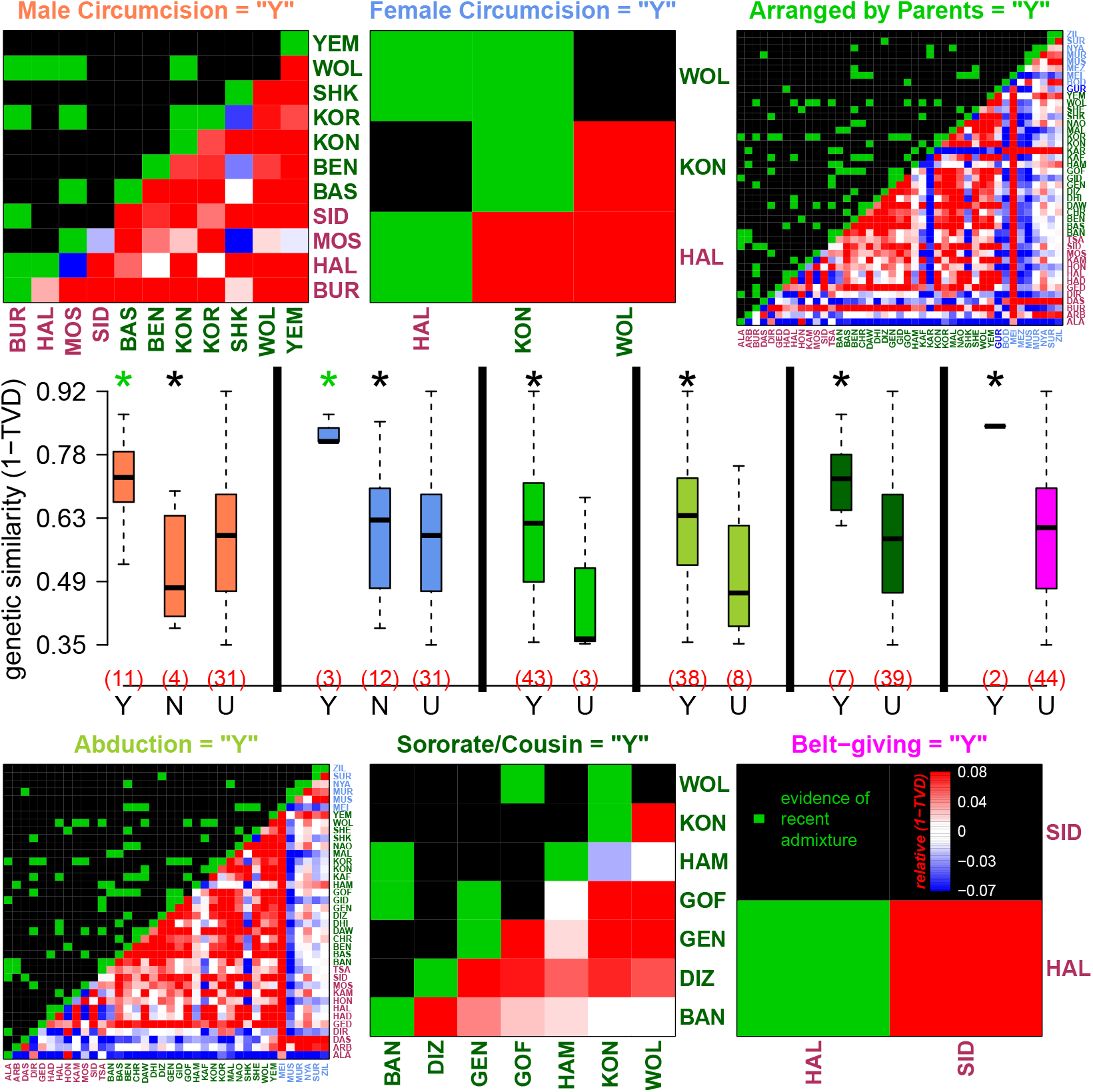
Sharing of self-declared cultural practices is associated with genetic similarity. Boxplots depict the pairwise genetic similarity (under the “Ethiopia-internal” analysis) among ethnic groups that reported practicing (“Y”), not practicing (“N”) or gave no information about practicing (“U”) each of six different cultural traits (labeled above heatmaps, with text colors matching the corresponding boxplots’ colors). Numbers of groups in each category are in parentheses in red. Stars above the boxplots denote whether there is a significant increase (p-val < 0.05) in genetic similarity among groups in (black) “Y” versus “U” or (green) “Y” versus “N”. The bottom right of each heatmap shows the increase (red) or decrease (blue) in average genetic similarity relative to that expected based on the ethnicities’ language classifications (key in bottom right heatmap), after accounting for the effects of spatial distance, between every pairing of ethnicities who reported practicing (“Y”) the given trait (axis labels colored by language group as in Fig 1a; group labels given in Table S1). Green squares in the top left portion of the heatmaps indicate whether >=1 pairings of individuals from different ethnic groups share atypically long DNA segments relative to all other comparisons of people from the two groups, which is indicative of recent intermixing between the two groups (see SI section 5).

## Discussion

Here we apply statistical analyses to a large-scale Ethiopian cohort densely sampled across ethnicities, geography and annotated for cultural practices (SI section 2). These resources enabled us to disentangle the myriad of factors contributing to the substantial genetic structure of Ethiopian populations. Wherever possible we only included individuals whose ethnicity matched that they reported for their parents and grandparents, which – if accurate – should exclude instances of ethnic re-identification and between-group intermixing occurring within the last two generations. This inclusion criterion is similar to a recent study of the UK (Leslie et al., 2015), and suggests that the patterns we have inferred reflect genetic patterns in Ethiopia approximately two generations prior to the present-day. This plausibly underrepresents genetic similarity and intermixing among ethnic groups that would be observable in a random sample, though our results support widespread recent intermixing among ethnic groups nonetheless (Fig 4).

We demonstrate how two different types of analyses, which we term “Ethiopia-internal” and “Ethiopia-external”, can disentangle relatively recent from ancient shared ancestry to better understand the origins of different ethnic groups (Fig S4-S5). As examples of this, minority discriminated groups included in this study, such as the Manjo from Kefa Sheka (Freeman & Pankhurst, 2003), the Manja from Dawro (Dea, 2007), the Ari/Wolayta Blacksmiths/Potters/Tanners (Biasutti, 1905; Pankhurst, 1999), the Chabu and the Negede-Woyto (Teclehaimanot, 1984; Legesse, 2013; Dira & Hewlett, 2017), each show relatively low genetic similarity to other Ethiopians using Fst (Extended Table 9) and under the “Ethiopia-internal” analysis (Fig 2a, Fig S8a, Extended Table 3). However, these genetic differences disappear under the “Ethiopia-external” analysis (Fig 2a, Fig S8b, Extended Table 4), suggesting that the high levels of genetic differentiation between discriminated peoples and other Ethiopians (e.g. measured by Fst) have arisen through their relatively recent isolation from other groups. Consistent with this isolation, these groups also exhibit signatures of recent endogamy as reflected by higher degrees of genetic homogeneity (Fig S3ab), with each of these groups separating into distinct clusters in ADMIXTURE analysis (Fig S11, Lawson et al 2018).

For example, we infer very similar sources and dates of admixture in independent analyses of distinct clusters that correspond to the occupational groups within the Ari (clusters 22, 24 and 25 in Fig 3, Fig S12) under the “Ethiopia-external” analysis, with overlapping 95% confidence intervals spanning 42-146 generations (Extended Table 5). A parsimonious explanation of these findings, matching our simulations (Fig S5), is that the ancestors of the Ari were a single population when these admixture events occurred. This in turn suggests the ancestors of different Ari occupational workers became isolated from one another only within the past ∼146 generations (<4200 years, assuming 28 years per generation, Fenner et al 2005). This correponds to the time period during which iron working is thought to have first appeared in Ethiopia (Phillipson, 2005) and supports the marginalisation theory of their origins (Lewis, 1962), which is consistent with findings from previous genetic studies (van Dorp 2015, Gopalan 2019).

Analogous to this, the Chabu, who are not linguistically classified by Ethnologue, share admixture events (dated to 300-900 years ago) and ancestry proportions similar to those inferred in the Mezhenger (Fig 3, Fig S12, Extended Table 5). These inferences are consistent with a high degree of intermarrying among the Chabu and Mezhenger, as has been proposed (Gopalan et al 2019; Anbessa & Unseth, 1989), and/or that these two groups split within the last ∼900 years. Nonetheless, we see the strongest overall average degree of genetic distance (TVD) between the Chabu and all other groups under the “Ethiopia-internal” analysis, consistent with previous reports of a decline in genetic diversity over the past 1000 years in the Chabu (Gopalan et al 2019). For the Negede-Woyto, the other group in this study for which there is no established linguistic classification in Ethnologue, we infer a relatively high amount of Egyptian-like ancestry (Fig 3, Fig S12), which is consistent with the group’s own origin narrative of a migration from Egypt by way of the Abay river (Teclehaimanot 1984). The ancestry proportions and admixture dates inferred in the Negede-Woyto are similar to those in the Beta Israel and Agaw, whom some scholars have proposed possible genealogical relationships with (Legesse, 2013), and show the highest average similarity to AA Semitic speakers (p-val > 0.05; Fig S9).

A caveat to the interpretation that groups with similar inference under the “Ethiopian-external” analysis share similar recent ancestry is that this analysis will have decreased (or no) power to discriminate between Ethiopian groups that indeed have separate ancestral sources if we have not included relevant non-Ethiopian groups to represent these separate sources. The large number of non-Ethiopian groups included in this sample, particularly those geographically proximal to Ethiopia, diminishes this possibility, but more samples from other sources, in particular DNA from ancient individuals in Ethiopia, may increase precision in identifying older ancestral differences between Ethiopians using these techniques.

Both analyses show a strong concordance between genetic differences and geographic distance among individuals (Fig 1b, Fig S6), analogous to that shown previously among peoples sampled from European (Novembre et al., 2008, Leslie et al., 2015), African (Tishkoff et al 2009, Scheinfeldt et al 2019) and worldwide countries (Li et al., 2008). We also show a correlation between genetic similarity and elevation difference, even after correcting for genetic similarity over geographic distances. Strikingly, we also see a correlation between spatial distance and the degree of genetic ancestry related to an ancient individual (Gallego-Llorente et al., 2015) whose remains were found in the Gamo Highlands of present-day Ethiopia 4,500 years ago (Fig S13). This suggests a notable preservation of some population structure in parts of Ethiopia (Gallego-Llorente et al., 2015, Gopalan et al., 2019).

While our “Ethiopia-internal” GLOBETROTTER results (Fig 4) suggest associations between genetics and geography are in part due to recent intermixing among nearby people, the “Ethiopia-external” SOURCEFIND and GLOBETROTTER results indicate varying patterns of non-Ethiopian-related ancestry that are also associated with geography (Fig 3, Fig S12). In particular Ethiopians in the southwest, typically NS speakers plus a few non-NS speaking groups (Chabu, Dasanech, Karo), are more related to Bantu and Nilotic speakers relative to AA speakers in the northeast that instead show more ancestry related to Egyptians and West Eurasians (Fig 3, Fig S12). The inferred timing and sources of admixture related to Egypt/W.Eurasian-like sources, starting around 100-125 generations ago (∼2800-3500 years ago; Fig 3, Fig S12) as in previous findings (Pickrell et al., 2014; Pagani et al., 2012), is consistent with significant contact and gene flow between the peoples of present day Ethiopia and northern Africa even before the rise of the kingdom of D’mt and interactions with the Saba kingdom of southern Yemen which traded extensively along the Red Sea (Currey, 2014; Phillipson, 2012). This timing is also consistent with trading ties between the greater Horn and Egypt dated back only to 1500 BCE, when a well-preserved wall relief from Queen Hateshepsut’s Deir el-Bahari temple shows ancient Egyptian seafarers heading back home from an expedition to what was known as the Land of Punt (SI Section 1A). On the other hand, inferred admixture dates in groups with varying amounts of ancestry related to Bantu and Nilotic speakers are dated to <1100 years ago, with the exceptions of the NS-speaking Kwegu (∼1500 years ago) and a second inferred older date (>1400 years ago) in the NS-speaking Meinit, which may reflect recent intermixing of NS-speakers with other Ethiopians. Such recent intermixing is consistent with mixed ancestry signals we see in some NS groups (e.g. see clusters containing Berta, Meinit and Nyangatom in clusters 15, 48-50 in Fig 3, Fig S12).

To facilitate comparison, our SOURCEFIND analysis included reference groups related to the four proxies used for ancestry sources in ancient and present-day East African groups reported in Prendergast et al 2019 (see SI section 5). We excluded their aDNA samples as reference groups, because they reported them to be admixed by these four sources. While using different reference groups and techniques complicates direct comparisons, our inferred sources of ancestry broadly agree with that study. For example, the Agaw (clusters 66, 67) have relatively more Levant-like ancestry (which we match most closely to Egypt), the Ari (clusters 22, 24, 25; called Aari in Prendergast et al 2019), have relatively more Mota-like ancestry and the Ethiopian Mursi (cluster 2) have relatively more Dinka-like ancestry (Fig 3, Fig S12). Simulations mimicking the admixture inferred here show high accuracy in inferred dates and sources, though illustrate an inherent limitation that older dates of admixture (e.g. those reported in Prendergast 2019) may be masked by more recent admixture (Fig S5). Thus complex intermixing events, such as those exhibited here, can be difficult to dissect fully with these approaches and sample sizes, e.g. distinguishing between multiple pulses or continuous admixture. A potential example are the NS-speaking Berta (clusters 48, 50), in which we infer only a single recent date of admixture but whom have complicated sources of ancestry that suggest multiple events (Fig 3, Fig S12).

Interestingly, the association between culture and genetic similarity is only apparent under the “Ethiopia-internal” analysis that is more sensitive to recent shared ancestry (Table S9). Another example consistent with this trend is that the NS-speaking Suri, Mursi, and Zilmamo, the only three Ethiopian ethnic groups that share the practice of wearing decorative lip plates, show atypically high genetic similarity under the “Ethiopia-internal” analysis but similarity levels comparable to other NS speakers under the “Ethiopia-external” analysis (Fig S14, Table S12). This suggests a recent separation of these groups, i.e. more recent than they separated from all other sampled Ethiopian groups, and/or recent intermixing among them.

Overall these examples illustrate how genetic data provide a rich additional source of information that can either corroborate or conflict with records from other disciplines (linguistics, geography, archaeology, anthropology, sociology and history) while adding further details and/or novel insights and directions for future investigation. Our interactive map is designed to facilitate evaluation of the genetic evidence for such records, providing results from both the “Ethiopia-internal” and “Ethiopia-external” analysis to enable comparisons analogous to the examples above. Future work can compare these and other published genetic results (e.g. Scheinfeldt et al 2019, Gopalan et al 2019, Prendergast et al 2019) to oral histories recorded for various ethnic groups. For example, some Mezhenger report that their ancestors originally migrated from Sudan to the present-day Gambella Regional State where Anuak lived, after which they migrated with the AA Omotic-speaking Sheko for a period before settling in their present-day homeland (The Council of Nationalities, Southern Nations and Peoples Region, 2017). Consistent with this, in the “Ethiopia-external” analysis the Mezhenger have high inferred ancestry matching to the Sudanese Dinka (Fig 3, Fig S12, Extended Table 5), and in the “Ethiopia-internal” analysis they have an inferred admixture event ∼300-600 years ago among three sources that are best represented by clusters containing the Anuak, Sheko and other NS-speaking groups near the Mezhenger (Extended Table 6).

Our study also highlights the importance of considering topographical and cultural factors, in particular language, ethnicity and in some cases occupation, when designing sampling strategies for future Ethiopian genetic studies, e.g. genome-wide association studies (GWAS), which our interactive map can also assist with. Similar sampling strategies may be necessary to capture the genetic structure of peoples in some other African countries that also exhibit relatively high levels of genetic diversity (Tishkoff et al., 2009, Busby et al., 2016). Finally, our analyses illustrate how cultural practices, e.g. participation in certain cultural and marriage customs, can operate as both a barrier and a facilitator of gene flow among groups and consequently act as an important factor in human diversity and evolution.

## Materials and Methods

### Samples

DNA samples from the 1,082 Ethiopians whose autosomal genetic variation data are newly reported in this study (following quality control, see below) were collected in several field trips from 2000-2010, through a long-standing collaboration including researchers at University College London and Addis Ababa University, and with formal approval granted by the Ethiopian Science and Technology Commission and National Ethics Review Committee and by the UK ethics committee London Bentham REC (formally the Joint UCL/UCLH Committees on the Ethics of Human Research: Committee A and Alpha, REC reference number 99/0196, Chief Investigator MGT). Analyses reported here were approved by the UCL Research Ethics Committee (reference number 5188/001). All study participants, including non-Ethiopians whose genetic variation data are newly reported in this study, gave their informed consent. Local permissions were obtained in all cases where applicable local ethical approval and regulations existed, e.g. Cameroon, Ministry of Higher Education and Scientific Research, Permits 0188/MINREST/B00/D00/D10/ D12 and 317/MINREST/B00/D00/D10 and University of Yaounde I; Ethiopia, Ethiopian Science and Technology Commission. Sample collection/usage for all unpublished data included in this study were approved by the UK ethics committee London Bentham REC (formally the Joint UCL/UCLH Committees on the Ethics of Human Research: Committee A and Alpha, REC reference number 99/0196, Chief Investigator MGT). The analyses reported here were approved by UCL REC (Project ID: 5188/001).

Buccal swab samples were collected from anonymous donors over 18 years of age, unrelated at the paternal level. For all individuals we recorded their, their parents’, paternal grandfather’s and maternal grandmother’s village of birth, language, cultural ethnicity and religion. In order to mitigate the effects of admixture from recent migrations that may be causing any genetic distinctions between ethnic groups to blur, analogous to Leslie et al. (2015), where possible we genotyped those individuals whose grandparents’ birthplaces and ethnicity were coincident. However, for a few ethnic groups (Bana, Meinit, Negede Woyto, Qimant, Shinasha, Suri), we did not find any individuals fulfilling this birthplace condition; in such cases we randomly selected individuals whose grandparents had the same ethnicity. In these cases, the geographical location was calculated as the average of the grandparents’ birthplaces (see SI section 2). Information about elevation was obtained using the geographic coordinates of each individual in the dataset with the “Googleway” package. All the Ethiopian individuals included in the dataset are classified into 75 groups based on self-reported ethnicity (68 ethnic groups) plus occupations (Blacksmith, Cultivator, Potter, Tanner, Weaver) within the Ari and Wolayta ethnicities. Table S1 shows the number of samples from each Ethiopian population and ethnic group that passed genotyping QC and were used in subsequent analyses. Fig 1a shows the geographic locations (i.e. birthplaces) of the Ethiopian individuals, though jittered to avoid overlap.

We also incorporated 2,678 non-Ethiopians (after quality control below) from 264 labeled present-day populations for comparison. Among these, non-Ethiopian samples newly released in this study include 23 Arabs from Israel, 13 Arabs from Palestine, 8 Bedouins from Saudi Arabia, 18 Berbers from Morocco, 7 Kotoko from Cameroon, 6 Muganda/Baganda from Uganda, 6 Mussese from Uganda, 13 Senegalese and 12 Syrians. All newly reported DNA samples in this study were genotyped using the Affymetrix Human Origins SNP array, which targets 627,421 SNPs (prior to our quality control), and merged with the Human Origin datasets published by Lazaridis et al. (2014) and Lazaridis et al. (2016), excluding their haploid samples (some ancient humans and primates). To these data we also added present-day Indians and Iranians published by Broushaki et al., 2016 and Lopez et al., 2017, and genomes from present-day Africans published by Skoglund et al. (2017), Gurdasani et al. (2015) and Mallick et al (2019) (Table S2a). We also included 21 high coverage published ancient samples (>1X average coverage) from Africa (Schlebusch et al, 2017; Prendergast et al, 2019; Lipson et al, 2020), including GB20 ‘Mota’ from Ethiopia (Gallego-Llorente et al., 2015), and 19 high coverage (>5X) published ancient non-African samples (Broushaki et al, 2016; Haak et al, 2015; Hofmanova et al, 2016; Keller et al, 2012; Mathieson et al, 2015) (Table S2b).

BAM files for ancient samples were downloaded from the ENA website (https://www.ebi.ac.uk/ena), with each file checked for correct format and metadata using PicardTools. We estimated post-mortem damage using ATLAS (Link et al 2017) with “pmd”, recalibrating each BAM file using ultra-conserved positions from UCNE (https://ccg.epfl.ch/UCNEbase/) and running ATLAS with “recal”, and then generated maximum likelihood genotype calls and phred-scaled genotype likelihood (PL) scores for each position using ATLAS with “call”. We used Conform-GT (https://faculty.washington.edu/browning/conform-gt.html) to ensure that strand was consistent with 1000 Genomes (1000 Genomes Project Consortium 2015) across present-day and ancient datasets, merging the data and running Beagle 4.1 (Browning and Browning 2016) with “modelscale=2” and the genetic maps at http://bochet.gcc.biostat.washington.edu/beagle/genetic_maps/plink.GRCh37.map.zip to re-estimate genotypes and impute missingness. We used vcf2gprobs, gprobsmetrics and filterlines (https://faculty.washington.edu/browning/beagle_utilities/utilities.html) to filter SNPs with an imputation accuracy of less than 0.98, and then we phased all samples using shapeit4 (Delaneau et al 2019) with “--pbwt-depth 16” and using their provided genetic maps.

To identify putatively related individuals, we used PLINK v1.9 (Chang et al., 2015) with “--genome” to infer pairwise PI_HAT values, after first pruning for linkage disequilibrium using “--indep-pairwise 50 10 0.1”. Instead of using the same fixed PI_HAT threshold value for all populations, we identified individuals with outlying PI_HAT values relative to other members of the same group label, in order to avoid removing too many individuals from populations with relatively low diversity. Specifically, we found all pairings of individuals from populations (i,k) that had PI_HAT > 0.15 and PI_HAT > min(X_i + 3*max{0.02,S_i} , Y_i + 3*max{0.02,D_i}, X_k + 3*max{0.02,S_k} , Y_k + 3*max{0.02,D_k}), where {X_i, Y_i, S_i, D_i} are the {mean, median, standard deviation, median-absolute-deviation}, respectively, of pairwise PI_HAT values among individuals from population i. For populations with <=2 sampled individuals, the standard deviation and median-absolute-deviations are undefined or 0; therefore in such cases we added to the list any pairings with PI_HAT > 0.15 that contained >=1 person from that population. Using a stepwise greedy approach, we then selected individuals from this list that were in the most pairs to be excluded from further analysis, continuing until at least one individual had been removed from every pair. This resulted in a total of 234 individuals removed, including 62 Ethiopians. All remaining Ethiopian pairs after this procedure had PI_HAT < 0.2.

Following the quality control described above, the total number of samples in the merge was 3,892, analyzed at 534,915 autosomal SNPs. We performed a principal-components-analysis (PCA) on the SNP data using smartpca (Patterson et al 2006, Price et al 2006) from EIGENSOFT version 7.2.0, with standard parameters and the lsqproject option. For the PCA of all individuals (Fig S2a), we used five outlier removal iterations (default), while for the PCA of only African individuals (Fig S2b), we did not perform any outlier removal iterations to prevent more isolated populations being removed.

### Genetic diversity and homogeneity

We used three different approaches to assess within-group genetic homogeneity in the Ethiopian ethnic groups. First, we computed the observed autosomal homozygous genotype counts for each sample using the --het command in PLINK v1.9 (Chang et al., 2015), taking the median value within each group. Second, we pruned SNP data based on linkage disequilibrium (--indep-pairwise 50 5 0.5), which left us with 359,281 SNPs, and used PLINK v1.9 to detect runs of homozygosity (ROH). This ROH procedure find runs of consecutive homozygous SNPs within groups that are identical-by-descent; here we report the total length of these runs per individual (Fig S3B). Third, we used FastIBD (Browning and Browning, 2011), implemented in the software BEAGLE v3.3.2, to find tracts (in basepairs) of identity by descent (IBD) between pairs of individuals. For each population and chromosome, fastIBD was run for ten independent runs using an IBD threshold of 10^−10^, as recommended by Browning and Browning (2011), for every pairwise comparison of individuals. For each population, we report the fraction of the genome that each pair of individuals shares IBD (Fig S3A).

We assessed whether the degree of genetic diversity in Ethiopian ethnic groups was associated with census population size, by comparing different measures of genetic diversity described above (homozygosity, IBD and ROH) with the census population size using standard linear regression (Fig S3C). As population census are not always available and can be inaccurate, we limited this analysis to ethnic groups in the SNNPR, for whom census information was recently reported (The Council of Nationalities, Southern Nations and Peoples Region, 2017).

### Using chromosome painting to evaluate whether genetic differences among ethnic groups are attributable to recent or ancient isolation

To quantify relatedness among individuals, we employed a “chromosome painting” technique, implemented in CHROMOPAINTER (Lawson et al., 2012), that identifies strings of matching SNP patterns (i.e. shared haplotypes) between a phased target haploid and a set of phased reference haploids. By modelling correlations among neighboring SNPs (i.e. “haplotype information”), CHROMOPAINTER has been shown to increase power to identify genetic relatedness over other commonly-used techniques such as ADMIXTURE and PCA (Lawson et al., 2012, Hellenthal et al., 2014, Leslie et al., 2015). In brief, at each position of a target individual’s genome, CHROMOPAINTER infers the probability that a particular reference haploid is the one which the target shares a most recent common ancestor (MRCA) relative to all other reference haploids. These probabilities are then tabulated across all positions to infer the total proportion of DNA for which each target haploid shares an MRCA with each reference haploid. We can then sum these total proportions across the reference haploids assigned to each of *K* pre-defined groups.

Following van Dorp et al. (2015), we used two separate CHROMOPAINTER analyses that differed in the *K* pre-defined groups used:

1. “Ethiopian-external”, which matches (i.e. paints) DNA patterns of each sampled individual to that of non-Ethiopians from *K*=264 groups only (Table S2a).
2. “Ethiopia-internal”, which matches DNA patterns of each sampled individual to that of all sampled groups, comprising 264 non-Ethiopian groups plus the 78 Ethiopian clusters defined in Fig S10 and the 4 Ethiopian groups from Mallick et al 2016, leading to *K*=346 groups total.

Relative to our genetic similarity score (1-TVD, described in the next section) under the “Ethiopia-internal” analysis, our score under the “Ethiopia-external” analysis mitigates the effects of any recent genetic isolation (e.g. endogamy) that may differentiate a pair of Ethiopians. This is because individuals from groups subjected to such isolation typically will match relatively long segments of DNA to only a subset of Ethiopians (i.e. ones from their same group) under analysis (1). However, this isolation will not affect how the same individuals match to each non-Ethiopian under analysis (2), for which they typically share more temporally distant ancestors. Consistent with this, in our sample the average size of DNA segments that an Ethiopian individual matches to another Ethiopian is 0.68cM in the “Ethiopia-internal” analysis, while the average size that an Ethiopian matches to a non-Ethiopian is only 0.23cM in the “Ethiopia-external” analysis, despite the latter analysis matching to substantially fewer individuals overall and hence having a higher *a priori* expected average matching length per individual.

Following López et al., 2017, van Dorp et al., 2019, and Broushaki et al., 2016, for each analysis (1) and (2) we estimated the CHROMOPAINTER algorithm’s mutation/emission (Mut, “-M”) and switch rate (Ne, “-n”) parameters using ten steps of the Expectation-Maximisation (E-M) algorithm in CHROMOPAINTER applied to chromosomes 1, 8, 15 and 22 separately, analysing only every ten of 4,081 individuals as targets for computational efficiency. This gave values of {321.844, 0.0008304} and {178.8922, 0.0006667} for {Ne, Mut} in CHROMOPAINTER analyses (1) and (2), respectively, after which these values were fixed in a subsequent CHROMOPAINTER run applied to all chromosomes and target individuals. The final output of CHROMOPAINTER includes two matrices giving the inferred genome-wide total expected counts (the CHROMOPAINTER “.chunkcounts.out” output file) and expected lengths (the “.chunklengths.out” output file) of haplotype segments for which each target individual shares an MRCA with every other individual.

### Inferring genetic similarity among Ethiopians under two different CHROMOPAINTER analyses

Separately for each of the “Ethiopia-internal” and “Ethiopia-external” CHROMOPAINTER analyses, for every pairing of Ethiopians *i,j* we used total variation distance (TVD) (Leslie et al., 2015) to measure the genetic differentiation (on a 0-1 scale) between their *K*-element vectors of CHROMOPAINTER-inferred proportions (with *K* defined above for both analyses), i.e:

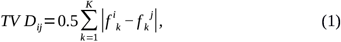

where *f* ^*i*^_*k*_ is the total proportion of genome-wide DNA that individual *i* is inferred to match to individuals from group *k* (see schematic in Fig S4). Throughout we report 1−*TV D*_*ij*_ , which is a measure of genetic similarity. When calculating the genetic similarity between two groups, we average (1−*TV D*_*ij*_) across all pairings of individuals *(i,j)* where the two individuals are from different groups (e.g. for Fig 2, Fig 5, Fig S8-S9, Extended Tables 3-4). We note an alternative approach to measure between-group genetic similarity is to first average each *f*^*i*^_*k*_ across individuals from the same group, and then use (1) to calculate TVD between the groups by replacing each *f* ^*i*^_*k*_ with its respective average value. Potentially this could give more power by reducing noise in the inferred copy vector for each group through averaging. However, here we instead use our approach of averaging (1−*TV D*_*ij*_) across individuals because of the considerable reduction in computation time when performing large numbers of permutations when assessing significance.

To test whether individuals from group *A* are more genetically similar on average to each other than an individual from group *A* is to an individual from group *B*, we repeated the following procedure 100K times. Let *n*^❑^_*B*_ and *n*^❑^_*A*_ be the number of sampled individuals from *A* and *B*, respectively, with *n*^❑^_*X*_=*min* (*n*_*A*_ ,*n*^❑^_*B*_) . First we randomly sampled *floor* (*n*^❑^_*X*_ / 2) individuals without replacement from each of *A* and *B* and put them into a new group *C*. If *n*^❑^_*X*_ /2 is a fraction, we added an additional unsampled individual to *C* that was randomly chosen from *A* with probability 0.5 or otherwise randomly chosen from *B*, so that *C* had *n*^❑^_*X*_ total individuals. We then tested whether the average genetic similarity, 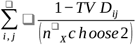, among all (*n*^❑^_*X*_ *choose* 2) pairings of individuals (*i,j*) from *C* is greater than or equal to that among all (*n*^❑^_*X*_ *choose* 2) pairings of *n*^❑^_*X*_ randomly selected (without replacement) individuals from group *Y*, where Y ∈ {A,B} (tested separately). We report the proportion of 100K such permutations where this is true as our one-sided p-value testing the null hypothesis that an individual from group *Y* has the same average genetic similarity with someone from their own group versus someone from the other group (Fig S8, Extended Tables 3-4). Overall this permutation procedure tests whether the ancestry profiles of individuals from *A* and *B* are exchangeable, while accounting for sample size and avoiding how some permutations may by chance put an unusually large proportion of individuals from the same group into the same permuted group.

For each Ethiopian group *A*, in Figure S8 and Extended Tables 3-4 we also report the other sampled group *A*_*max*_ with highest average pairwise genetic similarity to *A*. To test whether *A*_*max*_ is significantly more similar to group *A* than sampled group *B* is, we permuted the group labels of individuals in *A*_*max*_ and *B* to make new groups 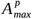 and *B*^*p*^ that preserve the respective sample sizes. We then found the average genetic similarity between all pairings of individuals where one in the pair is from *B*^*p*^ and the other from *A*, and subtracted this from the average genetic similarity among all pairings of individuals where one is from 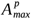 and the other is from *A*. Finally, we found the proportion of 100K such permutations where this difference is greater than that observed in the real data (i.e. when replacing *B*^*p*^ with *B* and 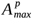 with *A*_*max*_), reporting this proportion as a p-value testing the null hypothesis that individuals from group *A*_*max*_ and group *B* have the same average genetic similarity to individuals from group *A*. For each *A*, any group *B* where we cannot reject the null hypothesis at the 0.001 type I error level (not adjusting for multiple testing) is enclosed with a white rectangle in Figure S8 and reported in Extended Tables 3-4.

As individuals are not allowed to match to themselves under the CHROMOPAINTER model, one potential issue with our paintings of Ethiopians under the “Ethiopian-internal” analysis is that each Ethiopian is allowed to match to one less individual in the cluster to which it is assigned relative to Ethiopians outside that cluster. For example, if cluster *A* contains ten Ethiopians, each of those Ethiopians are allowed to match to nine people from cluster *A* under the “Ethiopia-internal” analysis, while Ethiopians outside of cluster *A* are matched to all ten. This may create a slight discrepancy in the *f* ^*i*^_*k*_ values among Ethiopians for the 78 elements of *k* representing the Ethiopian clusters, which in turn may affect differences in TVD among Ethiopian group labels. To test this, we repeated the above using an alternative “Ethiopia-internal” painting where each Ethiopian is matched to all other Ethiopians from their cluster and *n*_*k*_ −1 Ethiopians from each other Ethiopian cluster *k* after randomly removing one individual, while matching to all individuals from every non-Ethiopian group as before. This gives a *K=346* length vector of *f* ^*i*^_*k*_ values for each Ethiopian *i* as before, but where each Ethiopian now has been painted against the same numbers of individuals from the *K* groups. We found that results change very little, e.g. with the TVD values among all pairwise combinations of Ethiopian groups (Fig S8A, Extended Table 3) having correlation *r>0*.*999*. This likely reflects how, for the given sample sizes in the *k* clusters, removing one individual from a cluster *k* results in people matching slightly more to the remaining *n*_*k*_ −1 individuals in that cluster, so that the total matching to *k* remains relatively unchanged. For comparison, in Extended Table 3 we provide columns at the far right end showing which groups were the closest match under this alternative “Ethiopia-internal” analysis; we note there are few changes relative to the original “Ethiopia-internal” analysis.

### Testing for associations between genetic similarity and spatial distance, shared group label, language and religious affiliation

To test for a significant association between genetic similarity and spatial distance, we used novel statistical tests that are analogous to the commonly-used Mantel test (Mantel, 1967) but that account for the non-linear relationships between some variables and/or adjust for correlations among more than three variables. We calculated genetic similarity (*G*_*ij*_) between individuals *i* and *j* as *G*_*ij*_ *= 1-TVD*_*ij*_, geographic distance (*d*_*ij*_) using the haversine formula applied to the individuals’ location information, and elevation distance (*h*_*ij*_) as the absolute difference in elevation between the individuals’ locations. We assessed the significance of associations between *G*_*ij*_ and *d*_*ij*_ and between *G*_*ij*_ and *h*_*ij*_ using 1000 permutations of individuals’ locations.

When using distance bins of 25km, we noted that the mean genetic similarity across pairs of individuals showed an exponential decay versus geographic distance in the “Ethiopia-internal” analysis (Fig 1b). Therefore, we assumed

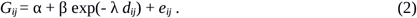

To infer maximum likelihood estimates (MLEs) for (α,β,λ), we first used the “Nelder-Mead” algorithm in optim() in R to infer the value of λ that minimizes the sum of *e*_*ij*_^*2*^ across all pairings of individuals *i,j* when α=0 and β=1, and then found the MLE for α and β under simple linear regression using this fixed value of λ. As the main observed signal of association between genetic and spatial distance is the increased *G*_*ij*_ at small values of *d*_*ij*_, (e.g. *d*_*ij*_=0, which is not always accurately fit via the Nelder-Mead algorithm), our reported p-values are the proportion of permutations for which the mean *G*_*ij*_ among all (*i,j*) with permuted *d*_*ij*_ < 25km is greater than or equal to that of the (unpermuted) real data (Table S6a).

In contrast, we noted a linear relationship between mean *G*_*ij*_ and *d*_*ij*_ in the “Ethiopia-external” analysis (Fig S6b) and between mean *G*_*ij*_ and *h*_*ij*_ when using 100km elevation bins under both analyses (Fig S6ac). Therefore, for these analysis we assumed:

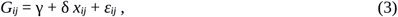

where *x*_*ij*_ = *d*_*ij*_ or *h*_*ij*_. Separately for each analysis, we found the MLEs for (γ,δ) using lm() in R. When testing for an association with elevation, we only included individual pairs (*i,j*) whose elevation distance was less than 2500km, which occurred in 730,880 (99.6%) of 733,866 total comparisons, to avoid undue influence from outliers. As we expect (and observe) the change in genetic similarity δ to be negative as spatial distance increases, our reported p-values provide the proportion of permutations for which the MLE of δ in the 1000 permutations is less than or equal to that of the real data (Table S6b-d).

As *d*_*ij*_ and *h*_*ij*_ are correlated (r = 0.22, Fig S7cd), we also assessed whether each was still significantly associated with *G*_*ij*_ after accounting for the other under the “Ethiopia-internal” analysis. To test whether geographic distance was still associated with genetic similarity after accounting for elevation difference, we assumed:

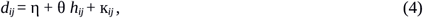

and used lm() in R to infer maximum likelihood estimates for (η,θ). Then to test for an association between genetic similarity and geographic distance after accounting for elevation, we used equation (2) but replacing *G*_*ij*_ with the fitted residuals *ε*_*ij*_ = *G*_*ij*_ - γ - δ*h*_*ij*_ from equation (2) and replacing *d*_*ij*_ with the fitted residuals К_*ij*_ = *d*_*ij*_ - η - θ *h*_*ij*_ from equation (4). We then repeated the procedure described above to calculate permutation-based p-values, first shifting К_*ij*_ to have a minimum of 0 (Table S6a,c). Similarly, to test for an association between genetic similarity and elevation difference after accounting for geographic distance, we replaced *x*_*ij*_ in equation (3) with the fitted residuals from an analogous model to (4) that instead regresses elevation on geographic distance, and replaced *G*_*ij*_ in equation (3) with the fitted residuals *e*_*ij*_ = *G*_*ij*_ - α - β exp(- λ *d*_*ij*_) from equation (2). We used the same permutation procedure described above to generate p-values (Table S6b,d).

We then tested whether sharing the same (A) self-reported group label, (B) language category of reported ethnicity, (C) self-reported first language, (D) self-reported second language, or (E) self-reported religious affiliation were significantly associated with increased genetic similarity after accounting for geographic distance or elevation difference. For (A), we used 75 group labels for (A) (Table S1), 66 first languages for (C), and 40 second languages for (D). For (B), we used the four labels in the second tier of linguistic classifications at www.ethnologue.com for which we have data (i.e. Afroasiatic Nilotic, Afroasiatic Semitic, Afroasiatic Cushitic, Nilo-Saharan Core-Satellite), excluding the Negede-Woyto and Chabu as they have not been classified into any language family. For (E), we compared genetic similarity across three religious affiliations (Christian, Jewish, Muslim), excluding religous affiliations recorded as “Traditional” as practices within these affiliations may vary substantially across groups.

To test whether each of these factors are associated with genetic similarity, we repeated the above analyses that use equations (2)-(4) while restricting to individuals (including permuted individuals) that share the same variable Y, separately for Y={A,B,C,D,E}. Our reported p-values give the proportion of permutations for which genetic similarity among permuted individuals sharing the same Y is more extreme than or equal to that of the real (un-permuted) data. For the “Ethiopia-internal” analysis when testing genetic similarity against geographic distance, this is the same p-value procedure as above, i.e. the proportion of permutations for which the mean *G*_*ij*_ among all (*i,j*) with permuted *d*_*ij*_ < 25km is greater than or equal to that of the (unpermuted) real data (Table S6a). When testing genetic similarity against geographic distance under the “Ethiopia-external” analysis, or testing genetic similarity against elevation difference under either analysis, this was instead defined as having any fitted value of *G*_*ij*_, at 48 equally-spaced bins of *d*_*ij*_ ∈{12.5,1187.5km} or 25 equally-spaced bins of *h*_*ij*_ ∈{50,2450m}, greater than or equal to that of the observed data.

As group label, language and religion can also be correlated with spatial distance and with each other (e.g. see Fig S7ab), we performed additional permutation tests where we fixed each of (A)-(E) when carrying out the permutations described above. For example, when fixing (A), we only permuted birthplaces and each of (B)-(E) across individuals within each group label, hence preserving the effect of group label on *G*_*ij*_. Applying this permutation procedure for each of (A)-(E), we repeated all tests described above, reporting p-values in Table S6.

For each of geographic distance, elevation difference, and (A)-(E), our final p-values reported in the main text and Fig 1b and Fig S6 that test for an association with genetic similarity are the maximum p-values across the six permutation tests that permute all individuals freely or fix each of (A)-(E) while permuting (i.e. the maximum values across rows of Table S6), with the following two exceptions. First, relative to the distances between birthplaces among all individuals, Ethiopians who share the same group label or who share the same first language live near each other (Table S7), so that permuting birthplaces while fixing group label or first language do not permute across large spatial distances. Therefore, we ignore those permutations when reporting our final p-values for geographic distance and elevation difference (i.e. in the main text and Fig 1b, Fig S6). Secondly, the high correlation between group label and first language (Fig S7ab) makes accounting for one challenging (in terms of loss of power) when testing the other. Furthermore, few permutations are possible when testing language group while accounting for group label (0 permutations available) or first language. Therefore, we excluded permutations fixing group and fixing first language when testing each of group, first language and language group when reporting our final p-values in the main text and Fig S6. Note we do observe a significant association with genetic similarity and ethnicity after accounting for spatial distance (geographic or elevation) and major language group, suggesting ethnicity explains genetic similarity beyond that of classifications according to the second language tier of at Ethnologue. We caution that these analyses assume that the relationships among genetic, geographic and elevation distance can be modelled with simple linear or exponential functions, which is sometimes debatable (Fig S7cd), indicating larger sample sizes may reveal deviations from these assumptions.

### Classifying Ethiopians into genetically homogeneous clusters

We used fineSTRUCTURE (Lawson et al 2012) to classify 1,268 Ethiopians (which includes all sampled Ethiopians except the eight Ethiopians from Mallick et al 2016 that were added later) into clusters of relative genetic homogeneity. To do so, we first used SHAPEIT (Deleneau et al., 2012) to jointly phase individuals using default parameters and the linkage disequilibrium-based genetic map build 37 (available at https://github.com/johnbowes/CRAFT-GP/find/master). We then employed CHROMOPAINTER to paint each individual against all others, i.e. in a manner analogous to the “Ethiopian-internal” analysis, though using a slightly different set of reference populations (e.g. samples from Mallick et al 2016 were not included due to unavailability at the time) and hence slightly different {Ne, Mut} values of {192.966, 0.000801}. We used default parameters, with the fineSTRUCTURE normalisation parameter “c” estimated as 0.20245. To focus on the fine-scale clustering of Ethiopians, we fixed all non-Ethiopian samples in the dataset as seven super-individual populations (Africa, America, Central Asia Siberia, East Asia, Oceania, South Asia and West Eurasia) that were not merged with the rest of the tree. We performed 2,000,000 sample iterations of Markov-Chain-Monte-Carlo (MCMC), sampling an inferred clustering every 10,000 iterations. Following Lawson et al. (2012), we next used fineSTRUCTURE to find the single MCMC sampled clustering with highest overall posterior probability. Starting from this clustering, we then performed 100,000 additional hill-climbing steps to find a nearby state with even higher posterior probability. This gave a final inferred number of 180 clusters containing Ethiopians. Results were then merged into a tree using fineSTRUCTURE’s greedy algorithm. We used a visual inspection of this tree to merge clusters, starting at the bottom level of 180 clusters, that had small numbers of individuals of the same ethnicity, as shown in Fig S10. After merging, we ended up with a total of 78 Ethiopian clusters.

We followed Leslie et al 2015 to generate a measure of cluster certainty using the last 100 fineSTRUCTURE MCMC samples. In particular for each of these 100 MCMC samples, we assigned a certainty score for each individual i being assigned to each final cluster j (out of 78) as the percentage of individuals assigned to the same cluster as individual i in that MCMC sample that are found in final cluster j. (For each individual i, note these percentages sum to 100% across the 78 final clusters.) For each combination of individual and final cluster, we averaged these certainty scores across all 100 MCMC samples. For each of our 78 final clusters, in Extended Table 2 we report the average certainty score of being assigned to that cluster across all individuals assigned to that cluster. This average certainty score had a mean of 44.7% across all clusters (range: 5.6-88.8%). For comparison, the average certainty score of being assigned to a cluster other than the final classification we used had a mean of 0.7% across all clusters (range: 0.1-1.2%). We note that clusters do not necessarily correspond to distinct groups that split from one another in the past, but instead provide a convenient means to increase power and clarity of ancestry inference by (i) merging people with similar genetic variation patterns, and (ii) separating individuals of the same self-identified label that have different genetic variation patterns.

#### Clustering Ethiopians using ADMIXTURE

We also used ADMIXTURE v.1.3.0 (Alexander et al 2009) to cluster Ethiopians. To do so, we first pruned the dataset for SNPs in linkage disequilibrium using PLINK v.2 (Chang et al 2015), removing SNPS with an r^2> 0.1 within a 50-SNP window, which left 139,032 SNPs. We then applied ADMIXTURE to the Ethiopians using these SNPs and a varying number of clusters K=2-15 and default parameters.

### Describing the genetic make-up of Ethiopians as a mixture of recent ancestry sharing with other groups

We applied SOURCEFIND (Chacón-Duque et al., 2018) to each of the 78 clusters to infer the proportion of ancestry that each clusters’ individuals share most recently with 275 ancestry surrogate populations, consisting of 264 present-day non-Ethiopian populations and aDNA samples from 11 populations including Mota (SI section 5). Briefly, SOURCEFIND identifies the reference groups for which each Ethiopian cluster shares most recent ancestry, and at what relative proportions, while accounting for potential biases in the CHROMOPAINTER analysis e.g. attributable to sample size differences among the surrogate groups. To do so, first each surrogate group and Ethiopian cluster *k* is described as a vector of length 264, where each element *i* in the vector for group *k* contains the total amount of genome-wide DNA that individuals from *k* are, on average, inferred to match to all individuals in group *i* under the “Ethiopia-external” CHROMOPAINTER analysis. These elements are proportional to the *f* ^*i*^_*k*_ described in the section “Inferring genetic similarity among Ethiopians under two different CHROMOPAINTER analyses” above. SOURCEFIND then uses a Bayesian approach to fit the vector for each Ethiopian cluster as a mixture of those from the 275 surrogate populations, inferring the mixture coefficients via MCMC (Chacón-Duque et al., 2018). In particular SOURCEFIND puts a truncated Poisson prior on the number of non-Ethiopian groups contributing ancestry to that Ethiopian cluster. We fixed the mean of this truncated Poisson to 4 while allowing 8 total groups to contribute at each MCMC iteration, otherwise using default parameters. For each Ethiopian cluster, we discarded the first 50K MCMC iterations as “burn-in”, then sampled mixture coefficients every 5000 iterations, averaging these mixture coefficients values across 31 posterior samples. In Extended Table 5 and Fig 3, Fig S12, we report the average mixture coefficients as our inferred proportions of ancestry by which each Ethiopian cluster relates to the 275 reference groups, though noting only 13 of these 275 contribute >5% to any cluster in these results.

### Identifying and dating admixture events in Ethiopia

Under each of the “Ethiopia-internal” and “Ethiopia-external” analyses, we applied GLOBETROTTER (Hellenthal et al., 2014) to each Ethiopian cluster to assess whether its ancestry could be described as a mixture of genetically differentiated sources who intermixed (i.e. admixed) over one or more narrow time periods (SI section 5). GLOBETROTTER assumes a “pulse” model whereby admixture occurs instantaneously for each admixture event, followed by the random mating of individuals within the admixed population from the time of admixture until present-day.

When testing for admixture in each Ethiopian cluster under the “Ethiopia-external” analysis, we used 130 groups (119 present-day groups and 11 ancient groups) as potential surrogates to describe the genetic make-up of the admixing sources, excluding groups that contributed little in the SOURCEFIND analysis for computational efficiency. When testing for admixture under the “Ethiopia-internal” analysis, we added as surrogates 64 of the 78 inferred Ethiopian clusters, removing 14 clusters (marked by asterisks in the first column of Extended Table 2) that contained small numbers of individuals from several ethnic groups and hence would confuse interpretation of results.

GLOBETROTTER requires two paintings of individuals in the target population being tested for admixture: (1) one that is primarily used to identify the genetic make-up of the admixing source groups (used as “input.file.copyvectors” in GLOBETROTTER), and (2) one that is primarily used to date the admixture event (used as the “painting_samples_filelist_infile” in GLOBETROTTER). For both the “Ethiopia-external” and “Ethiopia-internal” analyses, we used the respective paintings described in “Using chromosome painting to evaluate whether genetic differences among ethnic groups are attributable to recent or ancient isolation” above to define the genetic make-up of each group for painting (1). For (2), following Hellenthal et al. (2014), we painted each individual in the target cluster against all other individuals except those from the target cluster, using ten painting samples inferred by CHROMOPAINTER per haploid of each target individual. For the “Ethiopia-external” analysis, by design the painting in (2) is the same as the one used in (1). For the “Ethiopia-internal” analysis, we had to repaint each individual in the target group for step (2); to do so we used the previously estimated CHROMOPAINTER {Ne, Mut} parameters of {180.5629, 0.000610556}.

In all cases, we ran GLOBETROTTER for five mixing iterations (with each iteration alternating between inferring mixture proportions versus inferring dates) and performed 100 bootstrap re-samples of individuals to generate confidence intervals around inferred dates. We report results for null.ind = 1, which attempts to disregard any signals of linkage disequilibrium decay in the target population that is not attributable to genuine admixture when making inference (Hellenthal et al 2014). All GLOBETROTTER results, including the inferred sources, proportions and dates of admixture, are provided in Extended Tables 5-6 and summarized in Fig 3 and Fig S12 -- see SI Section S5 for more details. To convert inferred dates in generations to years in the main text, we used years ∼= 1975 - 28 x (generations + 1), which assumes a generation time of 28 years (Fenner 2005) and uses an average birthdate of 1975 for sampled individuals that matches our recorded information.

### Permutation test to assess significance of genetic similarity among individuals from different linguistic groups

To test whether individuals from language classification *A* are more genetically similar to each other than an individual from classification *A* is to an individual from classification *B*, we followed an analogous procedure to that detailed above to test for genetic differences between group labels *A* and *B*. Again let *n*^❑^_*A*_ and *n*^❑^_*B*_ be the number of sampled individuals from *A* and *B*, respectively, with *n*^❑^_*X*_=*min* (*n*_*A*_ ,*n*^❑^_*B*_) . For each of 100K permutations, we first randomly sampled *floor* (*n*^❑^_*X*_ / 2) individuals without replacement from each of *A* and *B* and put them into a new group *C*. If *n*^❑^_*X*_ /2 is a fraction, we added an additional unsampled individual to *C* that was randomly chosen from *A* with probability 0.5 or otherwise randomly chosen from *B*, so that *C* had *n*^❑^_*X*_ total individuals. We then tested whether the average genetic similarity, 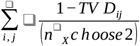, among all (*n*^❑^_*X*_ *choose* 2) pairings of individuals (*i,j*) from *C* is greater than or equal to that among all (*n*^❑^_*X*_ *choose* 2) pairings of *n*^❑^ _*X*_ randomly selected (without replacement) individuals from group *Y*, where *Y ∈ {A,B}* (tested separately). Individuals from the same ethnic/occupation label (i.e. those listed in Table S1) are often substantially genetically similar to one another (Fig S8, Extended Tables 3-4), which may in turn drive similarity among individuals within the same language classification. Therefore, whenever a language classification contained more than two different ethnic/occupation labels, we restricted our averages to only include pairings (*i,j*) that were from different ethnic/occupation labels (including in permuted group *C* individuals). We report the proportion of 100K such permutations where this is true as our one-sided p-value testing the null hypothesis that an individual from language classification *Y* has the same average genetic similarity with someone from their own language group versus someone from the other language group (Fig S9, Extended Tables 7-8). To test whether classifications *A* and *B* are genetically distinguishable, we take the minimum such p-value between the tests of *Y=A* and *Y=B* (Fig S9), which accounts for how some linguistic classifications include more sampled individuals and/or more sampled ethnic groups that therefore may decrease their observed average genetic similarity.

### Genetic similarity versus cultural distance

Between each pairing of 46 sampled SNNPR ethnic groups, we calculated a cultural similarity score as the number of practices, out of 31 reported in the SSNPR book (The Council of Nationalities, Southern Nations and Peoples Region, 2017) and described in SI Section 6, that the pair reported either both practicing or both not practicing (see Extended Table 10 for all groups’ recorded practices). Despite the SSNPR book also containing information about the Ari, we did not include them among these 46 because of the major genetic differences among caste-like occupational groups (Fig 2a). For the Wolayta, we included individuals that did not report belonging to any of the caste-like occupational groups.

We also calculated a second cultural similarity score whereby practices shared by many groups contributed less to a pair’s score than practices shared by few groups. To do so, if *H* ethnic groups in total reported participating in a practice, any pair of ethnicities that both reported participating in this practice added a contribution of *1*.*0/H* to that pair’s cultural similarity score, rather than a contribution of 1 as in the original cultural similarity score. Similarly, if *Z* ethnic groups in total reported not participating in a practice, any pair of ethnicities that both reported not participating in this practice added a contribution of *1*.*0/Z* to that pair’s cultural similarity score.

Genetic similarity, geographic distance and elevation difference between two ethnic groups *A, B* were each calculated as the average such measure between all pairings of individuals where *i* is from *A* and *j* from *B*. We then applied a mantel test using the *mantel* package in the *vegan* library in R with 100,000 permutations to assess the significance of association between genetic and cultural similarity across all pairings of ethnic groups (Table S9). We also used separate partial mantel tests, using the mantel.partial function in R with 100,000 permutations, to test for an association between genetic and cultural similarity while accounting for one of (i) geographic distance, (ii) elevation difference, or (iii) shared language classification (Table S9). To account for shared language classification, we used a binary indicator of whether *A,B* were from the same language branch: AA Cushitic, AA Omotic, AA Semitic, NS Satellite-Core.

For each of the 31 cultural practices, all 46 ethnic groups were classified as either (i) reporting participation in the practice, (ii) reporting not participating in the practice or (iii) not reporting whether they participated in the practice. For cultural practices where at least two of (i)-(iii) contained >=2 groups, we tested the null hypothesis that the average genetic similarity among groups assigned to category X was equal to that of groups assigned to Y, versus the alternative that groups in X had a higher average genetic similarity to each other. To do so, we calculated the difference in mean genetic similarity among all pairs of groups assigned to X versus that among all pairs assigned to Y. We then randomly permuted ethnic groups across the two categories 10,000 times, calculating p-values as the proportion of times where the corresponding difference between permuted groups assigned to X versus Y was higher than that observed in the real data. For 16 of 31 cultural practices, we tested X=(i) versus Y=(iii). For one cultural practice, we tested X=(ii) versus Y=(iii). For three cultural practices, we tested {X=(i) versus Y=(ii)}, {X=(i) versus Y=(iii)}, and {X=(ii) versus Y=(iii)}.

Six practices gave a p-value < 0.05 for one of the above permutation tests (Fig 5). These p-values remained after first adjusting for spatial distance as described in this paragraph. We calculated the average genetic similarity between all ethnic groups sharing these six practices after accounting for the effects of spatial distance and language classification. To account for spatial distance, we used equations (2)-(4) above, first adjusting geographic distance out of each of genetic similarity and elevation difference, and then regressing the residuals from the genetic similarity versus geographic distance regression against the residuals from the elevation difference versus geographic distance regression. We take the residuals for individuals *i,j* from this latter regression as the adjusted genetic similarity between individuals *i* and *j* (denoted *G\**_*ij*_). In each of the above regressions, we fit our models using all pairs of Ethiopians that were not from the same language classification at the branch level (i.e. AA Cushitic, AA Omotic, AA Semitic, NS Satellite-Core), in order to account for only spatial distance effects that are not confounded with any shared linguistic classification. We calculate the average spatial-distance-adjusted genetic similarity between each ethnic group *A,B* as the average *G\**_*ij*_ between all pairings of individuals where *i* is from *A* and *j* from *B*. Then to adjust for language classification, we calculated the expected spatial-distance-adjusted genetic similarity for each pairing of language branches *C,D* as the average adjusted genetic similarity across all pairings of ethnic groups *A, B* where *A* is from *C* and *B* is from *D*. For each pair of ethnic groups that share a reported cultural trait shown in Fig 5, we show the adjusted genetic similarity between that pair minus the expected spatial-distance-adjusted genetic similarity based on their language classification. This therefore illustrates the genetic similarity between the two groups after adjusting for that expected by their spatial distance from each other and their respective language classifications (lower right triangles of heatmaps in Fig 5).

For each of these six cultural practices shown in Fig 5, we also assessed whether there was evidence of recent intermixing among people from pairs of groups that both reported the given practice (see SI section 5). To do so, we indicate in the upper left triangles of the heatmaps in Fig 5 whether >=1 pairings of individuals, one from each group, have average CHROMOPAINTER-inferred MRCA segments >= 2.5cM longer than the median length of average inferred MRCA segments across all such pairings of individuals from the separate groups. We calculated the proportion out of 10,000 random samples of *n* groups (sampled from the 46 SNNPR groups analysed here) where a greater or equal number of group pairings showed this trend, also considering various different values of excess average MRCA segment size (Table S11).

## Supporting information

SOM

## Acknowledgements

This work is funded by BBSRC (Grant Number BB/L009382/1). GH is supported by a Sir Henry Dale Fellowship jointly funded by the Wellcome Trust and the Royal Society (Grant Number 098386/Z/12/Z) and supported by the National Institute for Health Research University College London Hospitals Biomedical Research Centre. We also acknowledge the UCL Biosciences Big Data equipment grant from BBSRC (BB/R01356X/1). We thank David Reich and the Children’s Hospital of Philadelphia for genotyping the samples on the Human Origins array. We thank Karl Skorecki for assistance with sampling. We also thank three anonymous reviewers for their helpful comments.

## Notes

### Competing Interest Statement

The authors have declared no competing interest.

## References

Alexander, D. H., Novembre, J. & Lange, K. Fast modelbased estimation of ancestry in unrelated individuals. Genome Res. 19, 1655-1664 (2009).

Allentoft, M.E. et al. Population genomics of Bronze Age Eurasia. Nature 522:167-172 (2015).

Anbessa, T. & Unseth, P. Toward the classification of Chabu (Mikeyir). In: Topics in Nilo-Saharan linguistics (Helmut Buske, Hamburg, 1989).

Appleyard, D. Preparing a Comparative Agaw Dictionary In: Cushitic & Omotic Languages (ed. Griefenow-Mewis & Voigt, 1996)

Ash, G. I. et al.. No association between ACE gene variation and endurance athlete status in Ethiopians. Med. Sci. Sports Exerc. 43, 590-597 (2011).

Baker, J.L., Rotimi, C.N., & Shriner, D. Human ancestry correlates with language and reveals that race is not an objective genomic classifier. Scientific Reports 7:1572 (2017).

Behar, D. M. et al.. The genomewide structure of the Jewish people. Nature 466, 238-242 (2010).

Biasutti, R. Pastori, agricoltori e cacciatori nell ’Africa Orientale’. Bolletino del Reale Societa geografica italiana 6, 155–179 (1905).

Blench, R. Archaeology, Language, and the African Past (Rowman Altamira, 2006)

Boattini, A. et al.. mtDNA variation in East Africa unravels the history of AfroAsiatic groups. Am. J. Phys. Anthropol. 150, 375-385 (2013).

Broushaki, F. et al.. Early Neolithic genomes from the eastern Fertile Crescent. Science 353, 499-503 (2016).

Browning, S.R. & and Browning, B.L. Genotype imputation with millions of reference samples. Am J Hum Genet 98, 116-126, (2016).

Browning, S. L., Tarekegn, A., Bekele, E., Bradman, N. & Thomas, M. G. CYP1A2 is more variable than previously thought: a genomic biography of the gene behind the human drug metabolizing enzyme. Pharmacogenet. Genomics 20, 647-664 (2010).

Browning, S. R. & Browning, B. L. Haplotype phasing: existing methods and new developments. Nat. Rev. Genet. 12, 703–714 (2011).

Busby, G. et al.. Admixture into and within sub-Sarahan Africa. eLife 5, e15266 (2016).

Byrne, R. P. et al.. Insular Celtic population structure and genomic footprints of migration. PLoS Genet. 14, e1007152 (2018).

CavalliSforza, L. L., Piazza, A., Menozzi, P. & Mountain, J. Reconstruction of human evolution: bringing together genetic, archaeological, and linguistic data. Proc. Natl. Acad. Sci. U. S. A. 85, 6002-6006 (1988).

ChacónDuque, J. C. et al.. Latin Americans show widespread Converso ancestry and imprint of local Native ancestry on physical appearance. Nat. Commun. 9, 5388 (2018).

Chang, C. C. et al.. Secondgeneration PLINK: rising to the challenge of larger and richer datasets. GigaScience 4 (2015).

Creemer, O. J. et al.. Contrasting exome constancy and regulatory region variation in the gene encoding CYP3A4: an examination of the extent and potential implications. Pharmacogenet. Genomics 26, 255-270 (2016).

Currey, J. Foundations of an African Civilisation: Aksum and the northern Horn (Oxford, 2014).

Dea, D. Rural Livelihoods and Social Stratification Among The Dawro, Southern Ethiopia. (Addis Ababa: Social Anthropology Dissertation Series, No 14, Series Editor Gebre Yntiso, Department of Sociology and Social Anthropology, Addis Ababa University, 2007).

Delaneau, O., Marchini, J. & Zagury, J. F. A linear complexity phasing method for thousands of genomes. Nat. Methods 9, 179–181 (2012).

Delaneau, O. et al.. Accurate, scalable and integrative haplotype estimation. Nature Communications 10, 5436 (2019).

Dira, S. J. & Hewlett, B. S. The Chabu huntergatherers of the highland forests of Southwestern Ethiopia. Hunt. Gatherer Res. 3, 323-352 (2017).

Ehret, C. Do Krongo and Chabu belong in Nilo-Saharan? In: Nilo-Saharan Linguistic Analysis and Documentation (Rüdiger Köppe Verlag, Cologne, 1992)

Fenner, J.N. Cross-cultural estimation of the human generation interval for use in genetics-based population divergence studies. Am J Phys Anthropol 128, 415–423 (2005).

Freeman, D. & Pankhurst, A. Peripheral People: The Excluded Minorities of Ethiopia (Hurst and Company, London, 2003).

GallegoLlorente, M. et al.. Ancient Ethiopian genome reveals extensive Eurasian admixture throughout the African continent. Science 350, 820-822 (2015).

Gopalan, S. et al.. Huntergatherer genomes reveal diverse demographic trajectories following the rise of farming in East Africa. BioRxiv. doi: https://doi.org/10.1101/517730 (2019).

Gurdasani, D. et al.. The African genome variation project shapes medical genetics in Africa. Nature 517, 327–332 (2015).

Haak, W. et al.. Massive migration from the steppe was a source for Indo-European languages in Europe. Nature 522: 207–211 (2015).

Harrison, G. A. Genetic and anthropological studies in the human adaptability section of the International Biological Programme. Philos Trans. R. Soc. Lond. B Biol. Sci. 274, 437-445 (1976).

Hellenthal, G. et al.. A genetic atlas of human admixture history. Science 343, 747–751 (2014).

Hofmanova, Z. et al.. Early farmers from across Europe directly descended from Neolithic Aegeans. PNAS 113:6686–6891 (2016).

HuertaSanchez, E. et al.. Genetic signatures reveal highaltitude adaptation in a set of Ethiopian populations. Mol. Biol. Evol. 30, 1877–1888 (2013).

Ingram, C. J. et al.. Multiple rare variants as a cause of a common phenotype: several different lactase persistence associated alleles in a single ethnic group. J. Mol. Evol. 69, 579-88 (2009).

Jones, B. L. et al.. Diversity of lactase persistence alleles in Ethiopia: signature of a soft selective sweep. Am. J. Hum. Genet. 93, 538-544 (2013).

Keller. A. et al. New insights into the Tyrolean Iceman’s origin and phenotype as inferred by whole-genome sequencing. Nature Communications 3:698 (2012).

Kivisild, T. et al.. Ethiopian mitochondrial DNA heritage: Tracking gene flow across and around the gate of tears. Am. J. Hum. Genet. 75, 752–770 (2004).

Lawson, D., Hellenthal, G., Myers, S. & Falush, D. Inference of population structure using dense haplotype data. PLoS Genet. 8: e1002453 (2012).

Lawson, D., van Dorp, L., & Falush, D. A tutorial on how not to overinterpret STRUCTURE and ADMIXTURE bar plots. Nature Communications 9: 3258 (2018).

Lazaridis, I. et al.. Ancient human genomes suggest three ancestral populations for presentday europeans. Nature 513, 409–413 (2014).

Lazaridis, I. et al.. Genomic insights into the origin of farming in the ancient Near East. Nature 536, 419-424 (2016).

Legesse, W. Religion and Everyday Life: NegedeWoyto Muslim Community of Bahir Dar Town Ethiopia (MA Thesis, Addis Ababa University, 2013).

Leslie, S. et al.. The finescale genetic structure of the British population. Nature 519, 309-314 (2015).

Levine, D. N. Greater Ethiopia: The Evolution of A Multiethnic Society, Second Edition (Chicago and London: The University of Chicago Press, 2000).

Lewis, H. Historical Problems in Ethiopia and the Horn of Africa. Ann. New York Acad. Sci. 96, 504–511 (1962).

Li, J. Z. et al.. Worldwide human relationships inferred from genomewide patterns of variation. Science 319, 1100-1104 (2008).

Liebert, A. et al.. Worldwide distributions of lactase persistence alleles and the complex effects of recombination and selection. Hum Genet. 136, 1445-1453 (2017).

Link, V. et al.. ATLAS: Analysis Tools for Low-depth and Ancient Samples. BioRxiv, doi.org/10.1101/105346 (2019).

Lipson, M. et al.. Ancient West African foragers in the context of African population history. Nature 577: 665–670 (2020).

López, S. et al.. The Genetic Legacy of Zoroastrianism in Iran and India: Insights into Population Structure, Gene Flow, and Selection. Am. J. Hum. Genet. 101, 353-368 (2017).

Mallick, S. et al.. The Simons Genome Diversity Project: 300 genomes from 142 diverse populations. Nature 538:201-206 (2016).

Mantel, N. The detection of disease clustering and a generalized regression approach. Cancer Res. 27, 209–220 (1967).

Mathieson, I. et al.. Genome-wide patterns of selection in 230 ancient Eurasians. Nature 528:499–503 (2015).

Messina, F. et al.. Linking between genetic structure and geographical distance: Study of the maternal gene pool in the Ethiopian population. Ann. Hum. Biol. 44, 53-69 (2017).

Mourant, A. E. et al.. The blood groups and haemoglobins of the Kunama and Baria of Eritrea, Ethiopia. Ann. Hum. Biol. 1, 383-392 (1974).

Non, A. L., AlMeeri, A., Raaum, R. L., Sanchez, L. F. & Mulligan, C. J. Mitochondrial DNA reveals distinct evolutionary histories for Jewish populations in Yemen and Ethiopia. Am. J. Phys. Anthropol. 144, 1-10 (2011).

Novembre, J. et al.. Genes mirror geography within Europe. Nature 456, 98-101 (2008).

Olalde et al. Derived immune and ancestral pigmentation alleles in a 7,000-year-old Mesolithic European. Nature 507, 225–228 (2014).

Pagani, L. et al.. Tracing the route of modern humans out of Africa by using 225 human genome sequences from Ethiopians and Egyptians. Am. J. Hum. Genet. 96, 986-991 (2015).

Pagani, L. et al.. Ethiopian Genetic Diversity Reveals Linguistic Stratification and Complex Influences on the Ethiopian Gene Pool. Am. J. Hum. Genet. 91, 83–96 (2012).

Pankhurst, A. ‘Caste’ in Africa: The Evidence from SouthWestern Ethiopia Reconsidered. Africa 69, 485–509 (1999).

Patterson, N., Price, A.L. & Reich, D. Population Structure and Eigenanalysis. PLoS Genetics 2, e190

Phillipson, D. W. African Archaeology (Cambridge University Press, 2005)

Phillipson, D. W. Foundations of and African Civilisation: Aksum & The Northern Horn 1000 BC-AD 1300 (Suffolk & Rochester: James Currey, 2012)

Pickrell, J. K. et al.. Ancient west Eurasian ancestry in southern and eastern Africa. Proc. Natl. Acad. Sci. U. S. A. 111, 2632–2637 (2014).

Poloni, E. S. et al.. Genetic evidence for complexity in ethnic differentiation and history in East Africa. Ann. Hum. Genet. 73, 582-600 (2009)

Prendergast, M.E. et al.. Ancient DNA reveals a multistep spread of the first herders into sub-Saharan Africa. Science 365, eaaw6275 (2019).

Price A. L. et al.. Principal components analysis corrects for stratification in genomewide association studies. Nat. Genet. 38, 904–909 (2006).

Rankinen, T. et al.. No Evidence of a Common DNA Variant Profile Specific to World Class Endurance Athletes. PLoS One 11, e0147330 (2016).

Ronen, R., Zhou, D., Bafna, V. & Haddad, G. G. The genetic basis of chronic mountain sickness. Physiology (Bethesda) 29, 403-412 (2014).

Scheinfeldt, L. B. et al.. Genetic adaptation to high altitude in the Ethiopian highlands. Genome Biol. 13: R1 (2012).

Scheinfeldt, L. B. et al.. Genomic evidence for shared common ancestry of East African hunting gathering populations and insights into local adaptation. Proc. Natl. Acad. Sci. U. S. A. pii: 201817678 (2019).

Schlebusch, C.M. et al.. Southern African ancient genomes estimate modern human divergence to 350,000 to 260,000 years ago. Science 358, 652-655 (2017).

Scott, R. A. et al.. Mitochondrial DNA lineages of elite Ethiopian athletes. Comp. Biochem. Physiol. B Biochem. Mol. Biol. 140, 497-503 (2005).

Semino, O., SantachiaraBenerecetti, A. S., Falaschi, F., CavalliSforza, L. L. & Underhill, P. A. Ethiopians and Khoisan share the deepest clades of the human Ychromosome phylogeny. Am. J. Hum. Genet. 70, 265–268 (2002).

Sim, S. C. et al.. A common novel CYP2C19 gene variant causes ultrarapid drug metabolism relevant for the drug response to proton pump inhibitors and antidepressants. Clin. Pharmacol. Ther. 79, 103-113 (2006).

Simonson, T.S. Altitude Adaptation: A Glimpse Through Various Lenses. High Alt. Med. Biol. 16, 125-137 (2015).

Skoglund, P. et al.. Reconstructing Prehistoric African Population Structure. Cell 171, 59-71 (2017).

Stobdan, T. et al.. Endothelin receptor B, a candidate gene from human studies at high altitude, improves cardiac tolerance to hypoxia in genetically engineered heterozygote mice. Proc. Natl. Acad. Sci. U. S. A. 112: 10425-10430 (2015).

Tadesse, L., Tafesse, F. & Hamamy, H. Communities and community genetics in Ethiopia. Pan. Afr. Med. J. 18: 115 (2014).

Teclehaimanot, G. S. The Wayto of Lake Tana: An EthnoHistory (MA Thesis in History, School of Graduate Studies, Addis Ababa University, 1984).

The Council of Nationalities (CoN), Southern Nations and Peoples Region (SNNPR), (in Amharic). A Profile of the Nations, Nationalities and Peoples of the Southern Region, Second Edition (December 2004EC). Translated from Amharic and subedited by Ayele Tarekegn and Neil Bradman, Published by the Council of Nationalities with financial support from the German Co operation (GIZ), Addis Ababa: Berhanena Selam Printing Enterprise, April 2017 CE.

Council of Nationalities (CoN), Southern Nations, Nationalities and Peoples’ Region (SNNPR), (in Amharic). Yedebubbiheroch, biheresebochna Hizboch Profile (title translated as A Profile of the Nations, Nationalities and Peoples of the Southern Region), Third Edition, Published by the CoN, SNNPR, Addis Ababa: BerhanenaSelam Printing Enterprise, Miazia (April) 2008EC.

Thomas, M. G. et al.. Founding mothers of Jewish communities: geographically separated Jewish groups were independently founded by very few female ancestors. Am. J. Hum. Genet. 70, 1411-1420 (2002).

Tishkoff, S. A. et al.. The genetic structure and history of Africans and African Americans. Science 324, 1035-1044 (2009).

Todd, D. The Origins of Outcastes in Ethiopia: Reflections on an Evolutionary Theory. Abbay 9, 145–158 (1978).

van Dorp L. et al.. Evidence for a common origin of blacksmiths and cultivators in the Ethiopian Ari within the last 4500 years: Lessons for clusteringbased inference. PLoS Genet. 11, e1005397 (2015).

van Dorp, L. et al.. Genetic legacy of state centralization in the Kuba Kingdom of the Democratic Republic of the Congo. Proc. Natl. Acad. Sci. U. S. A. 116, 593-598 (2019).

Weir, B. S. & Cockerham, C. C. Estimating Fstatistics for the analysis of population structure. Evolution 38, 1358–1370 (1984).

