## Supplementary material for "The genetic landscape of Ethiopia: diversity, intermixing and the association with culture": SOM

### SI Section 1. A brief history of Ethiopia with particular reference to foreign contacts

#### SI Section 1A. Summary

The modern state of Ethiopia, located in the Horn of Africa, encompasses approximately 1.1M km of varied climates and environments, including mountain ranges, high plateaus, forests, lakes and low, arid, extremely hot regions (Marcus, 2002). The Rift Valley, the largest geographic trench on earth which has yielded important hominid fossils, runs northeast-southwest across the country. The population of Ethiopia now exceeds 100 million with groups speaking over 80 languages classified as Semitic, Cushitic, Omotic and Nilo-Saharan. Christian, Muslim, Traditional and Jewish religions are all represented.

The country's archaeological record extends more than 3 million years with genetic and other evidence suggesting that East Africa is possibly the region (Quintana-Murci et al., 1999) via which the majority of anatomically modern humans left the African continent to people the rest of the world. *Ardipithecus ramidus*, from the Afar region of northeast Ethiopia, remains the world's oldest hominid fossil find at 4.4M years ago.

It is not thought that the expansion of the Bantu speaking peoples, approximately 5,000ybp, extended into the area covered by the modern Ethiopian state (Vansina, 1995; Ashley, 2010; Clist, 1987). Contacts between the north and, respectively, Egypt and Arabia, although varying in intensity, have continued for at least the past 6,000 years (Currey, 2014; Phillipson, 2012).

While Aksum, in the northern region, was in contact with the world of antiquity (Phillipson, 2012), the south was only incorporated into what was then commonly known in the western world as Abyssinia in the late nineteenth century CE.

Before the 19th Century interaction with European powers was limited, although in the 16th Century Portuguese emissaries had a six-year presence (Alvarez, 1961), during which plants from the Americas were introduced to local farmers. For five years from 1936 to 1941 Italy occupied the country. The independence of the modern state was recognised internationally in 1947.

**SI Section 1B. The Pre-History of Ethiopia**

Ethiopia can legitimately claim the longest archaeological record of any country in the world (Phillipson, 1998). The discovery in November 1974 of the first and most famous *Australopithecus* specimen named *afarensis* or Lucy (with 40% of her osteological remains) was followed by a number of hominid, and later fossil remains, including Selam, the oldest (dated 4.3 MYA) and most complete (60%) to date (Kalb, 2001; Johanson, 2009; Suwa et al., 2009; Lovejoy et al., 2009). There appears to be broad scholarly consensus on the antiquity of Ethiopian agriculture (Brandt, 1984; Phillipson, 1993). By about 7500 YBP, the Afroasiatic family of languages appeared in what is now eastern Sudan, northeast of Khartoum. Wild grass would already have been collected as food in the greater Horn of Africa region by this time. Not long afterwards, domestication of plants in the greater Horn may have begun. Nicolai Ivanovich Vavilov (1955) proposed that Ethiopia was probably one of eight centres of plant domestication in the world. At about the fourth millennium BCE, fauna and flora appear to have been introduced into the country from abroad. The fauna include cattle,[1] sheep[2] and goats and the flora wheat, barley and sorghum, all introduced from a likely West Asian source. In later times, Ethiopians maintained close ties with the Graeco-Roman and Eastern Mediterranean worlds. The earliest surviving evidence for trading ties between the greater Horn and Egypt date back to 1500 BCE, when a well-preserved wall relief from Queen Hateshepsut's Deir el-Bahari temple showed ancient Egyptian seafarers heading home from an expedition to what was known as the Land of Punt. (The Land of Punt is generally accepted by modern scholarship to be a reference to the greater Horn of Africa region (Phillips, 1997).) It is very likely that there was wider involvement in the Red Sea trade involving the Arabian Peninsula and the Eastern Mediterranean regions. Contacts with South Arabia have been better attested archaeologically than those with Nubia and Egypt.[3]

**SI Section 1C. Ethiopian Christianity and contacts with Egypt, the Levant and Europe**

Sometime during the second quarter of the fourth-century CE Aksum adopted Christianity (Sergew 1972). Unlike corresponding events in the Roman Empire, the first to be converted to the new creed in Ethiopia were the ruling class. Palaeographic and documentary sources show that the first Christian ruler of Aksum was King Ezana.[4] The conversion of the ruling class to the new faith seems to have resulted in important transformations that provided politico-ideological legitimacy to the monarchy and also in the sphere of the wider highland Ethiopian popular culture. The celebrated document known as the *Kibra Nagast*, historic Ethiopia's national epic, represents an epitome of this event of revolutionary proportions (Budge, 1922; Shahid, 1976). It took the new religion over a century to reach the broad masses through the evangelical activities of two groups of foreign missionaries, known as the *Tsadqan* and the "Nine Saints" (Sergew 1972). These missionaries were monks who came to Ethiopia from the Eastern Mediterranean region around the end of the fifth century CE. As sufficiently trained clergy, the missionaries appear to have completed the task of translating the Bible mainly from the Old Greek version (the Septuagint), which was translated from the older Hebrew/Aramaic versions (Polotsky 1964; Ullendorff 1968, 1980, 1987; Knibb 1988) into the local vernacular, Ge'ez. It is claimed that this literary activity led to the introduction of some foreign terms into the local language, for example, the Armenian word *adja* (adcha or adjar) for 'emmer wheat' as has long been used in Ethiopia (Harlan, 1969) and the Syriac term *haymanot* for 'religion' (Sergew, 1972). The missionaries appear to have established churches and monasteries, both built and rock-hewn, and generally helped propagate the new faith among the people.

The Alexandrine See came to have the status of spiritual suzerainty and guidance over the Ethiopian Orthodox Church (Taddesse, 1972). Invariably, all heads, known in Ge'ez as *Abun*, of the latter Church were Egyptian bishops, duly consecrated and appointed by the former down to 1959 when Ethiopian bishops begun to serve as the heads of their own Church. From the outset, a special relationship of mutual help and interdependence, appears to have been established between Church and State. The Church seems to have acted as the most prominent ideological arm of the State. In return, the latter endowed the former with massive material subsidies used to establish new churches and monasteries as well as proselytising among, first, believers of traditional religions and later Muslims. This arrangement was manifest, most markedly, during the fourteenth and fifteenth centuries (Taddesse, 1972). Throughout the medieval period, Ethiopian Orthodox Christian monks and devout lay pilgrims are known to have flocked, in comparatively large numbers, to Jerusalem where they made contact with people from other countries. An Ethiopian monk named Father Gregory provided the famous German scholar Hiob Ludolf with reliable material to write his *A New History of Ethiopia* (published in 1682). This book was the second work to provide generally correct information about the country to western Europe following the account of the Portuguese Embassy of 1520-1526 referred to below.

Notwithstanding the contacts described above, Ethiopia was relatively isolated from the early-seventh century to the late nineteenth century. However, limited, more or less regular, contacts were maintained with the outside world, chiefly Egypt, the Holy Land, and, to a lesser extent, the Vatican (Taddesse, 1972; Sergew, 1972). Ideas were exchanged and knowledge filtered through in what may have arguably been a two-way traffic. Since the mid-fourteenth century, Ethiopia also maintained some communication with western Europe through visiting travellers and explorers, diplomats, Christian missionaries and scholars (Crawford, 1958; Ullendorff, 1960). In 1520 a fourteen-man Portuguese Embassy, which included a chaplain-chronicler, Father Francesco Alvares, visited the country establishing a well-documented official link between Ethiopia and a European country (Alvarez, 1961; Castanhoso, 1902). The Embassy remained until 1526 when it returned to Lisbon (Alvarez, 1961). Detailed accounts of the venture and observations were published in 1540. It is thought that the Portuguese were instrumental in introducing New World flora including pepper and perhaps also corn, cotton, and beans. Later Jesuit missionaries, led by the Spaniard Father Pedro Paez, succeeded (after years of persuasion) in converting Emperor Susneyos (r. 1607-1632) and some of his most important courtiers to Catholicism in 1622.

Jesuit involvement in the country led to religious wars. Emperor Susneyos abdicated in favour of his son, Emperor Fasil(adas) (r. 1632-1667), who founded Gondar as Imperial Capital in 1634/5 and pursued a close-door policy banning all European visitors to the country (Merid, 1971; Berry, 1976). Despite this edict, some European travellers did succeed in entering the country.

**SI Section 1D. Later travellers to Ethiopia**

French physician Charles Poncet (1699-1700) and the Scottish traveller James Bruce (1790, 1813), both spent time in Gondar, as did an Armenian jeweller Yohannes T'ovmacean (1764-1766) who also left an important record (Nerssian & Pankhurst, 1982). Ethio-European relations accelerated in the early nineteenth century, when many more Europeans visited the country (Ullendorff, 1960; Rubenson, 1978; Malécot, 1972) with numbers increasing in the late nineteenth and early twentieth centuries.

The Ethiopian victory at the battle of Adwa on the 2nd of March 1896 resisting Italy's colonial ambitions led to the opening of permanent diplomatic missions representing many European countries, including Italy itself (1896), France and Britain (1897), the USA (1904) and subsequently many other nations. Today the capital of Ethiopia, Addis Ababa (founded in 1889 when King Menelik of Shawa became Emperor Menelik II of Ethiopia), hosts the headquarters of the Organisation of African Unity (renamed the African Union).

**Notes**

1 The earliest African attestation of cattle has been dated to ca. 9000 BCE.

2 Sheep are known in the Nile Valley from at least 5000 BCE (Muigai & Hanotte, 2013).

3 Phillipson (1998: 24) has the following to say on this subject: "*It is remarkable how few are the artefacts of demonstrably Egyptian origin that have been recovered from archaeological sites in Ethiopia and Eritrea … . Contacts with the Nile Valley, both in Nubia and further downstream, probably became stronger during Aksumite times, both through trade in raw materials (perhaps accompanied by military subjugation and through links between Christian Ethiopia and her co-religionists. This was often the route by which Ethiopian pilgrims travelled to Jerusalem, the place where most regular contact was established between Ethiopian and people from other countries*"

4 Legend has it that there were two Greek-speaking Christians from Egypt, named Frumentius and Aedesius, who had been taken captive from the Red Sea littoral and brought to the royal court in Aksum where a deceased king was survived by his minor son and the Queen Mother (Sergew 1972: pages 95-100). Frumentius and Aedesius reportedly succeeded in gradually converting the young monarch and his mother to embrace the new religion.

### SI Section 2. Sampled Ethiopian groups

Today Ethiopia is one of the most populated countries in the world ("Ethiopia". The World Factbook, CIA, 2017), home to over 90 ethnic groups and 86 living languages. The largest groups are the Oromo (34.4% of the total population – source: Ethiopia People 2017, theodora.com), the Amhara (27%), the Somali (6.2%) and the Tigraway (6.1%) that make up around three-quarters of the population. The rest of the groups represent low percentages of the total population, and some of them represent minorities with less than 10,000 members. Christianity is the most practiced religion in Ethiopia (62.8%), followed by Islam (33.9%), traditional faiths (2.6%) and other religions (0.6%) (Ethiopian Central Statistical Agency, 2009). Ethiopia is administratively divided into nine regions: Afar, Amhara, Benishangul-Gumuz, Gambela, Harari, Oromia, Somali, Southern Nations, Nationalities and Peoples' and Tigray. This is subdivided into 68 zones, that in turn are subdivided into districts or woredas and two chartered cities (Addis Ababa and Dire Dawa).

Languages spoken in Ethiopia can be classified into two language families: Afroasiatic and Nilo-Saharan. Branches of the Afroasiatic languages represented in Ethiopia are Cushitic, Omotic and Semitic (www.ethnologue.com), while Nilo-Saharan languages spoken in Ethiopia have been classified as members of two groups: Chari-Nile and Koman (Bender, 1976). Previous work has suggested that linguistic affiliation is the main factor driving genetic structure in Ethiopians (Pagani et al., 2012). Table S1 and Figure 1a show the languages associated with the samples included for this work.

For the Ethiopians, we genotyped those individuals whose grandparents' birthplaces were coincident, with the exception of a few ethnic groups (Meinit, Qimant, Suri, Negede-Woyto, Shinasha, Bana) where we did not find any sampled individual fulfilling this condition. In these cases, the geographical location was calculated as the average point between the birth locations of the paternal grandfather and the maternal grandmother. The latitude and longitude coordinates of birthplaces of donors were recorded as the roughly central place in the locality of their birth, which was obtained by one or other of the following means: on-site use of a GARMIN GPS unit during the data collection, information provided by a local service (Information Systems Services (ISS)) and manual searches using Google Maps, OpenStreetMap and, in a few cases, other programs. Information about elevation was obtained using the geographic coordinates of each individual in the dataset with the "Googleway" package.

Samples were collected by co-author AT in a programme that consisted of two collection expeditions a year taking place from 1998 to 2011 and which frequently involved communication through local translators in remote locations. Consistency is an extremely difficult thing to achieve in Ethiopian linguistic nomenclature. Neither the Ethnologue (www.ethnologue.com) nor many linguists consistently give primacy to the opinions of native speakers in ascribing a name to a language. Sometimes a group may declare that they speak a language that linguists may designate a dialect. Consequently the Ethnologue may use a common name for a cluster of closely related tongues. AT recorded the name of a language as declared by the speakers who were sampled. Similarly AT's practice was to use the self-adopted names of ethnic groups rather than names used to refer to them by their neighbours or other outsiders, all of which they are likely to consider derogatory. All information recorded with respect to donors was either provided by the individual donors themselves or by local informants. Information about linguistic classifications used in this study are provided in Table S1 and Extended Table 1. As examples of some of the complications, the labeled group "Manja" refers to a caste-like hunter-gatherer group claiming Kafa as their original language, but who presently live in the Dawro Zone and report Manja as their first language and Dawro as their second language. They claim to be descendants of a gift of slaves by the king of Kafa to the king of Dawro. The labeled group "Manjo" refers to a caste-like hunter-gatherer group, locally called Manjo, who live amongst the Kafa and the Shekacho in the Kafa and Sheka administrative Zones (previously the Kafa-Sheka Zone). Manjo study participants living in the Sheka Zones reported their first language as Shekacho and their second language as Amharic. More generally, it is believed that Manjo living in Sheka Zone speak Shekacho as their first language while Manjo living in Kafa Zone speak Kafa as their first language.

**SI Section 3. Simulations**

We performed simulations to illustrate how inference can differ under the “Ethopia-internal” versus “Ethiopia-external” analyses in the presence of population bottlenecks, differing levels of admixture from outside sources, and the intermixing of groups that contain previous admixture events. For example, we demonstrate how two populations splitting 45 generations (~1200-1300 years) ago, with one of the populations subsequently experiencing a strong bottleneck, can lead to high genetic differentiation under the “Ethiopia-internal” analysis but relatively little differentiation under the “Ethiopia-external” analysis. In contrast, two populations that have experienced admixture from the same sources, but at proportions differing by 20%, look genetically different under both analyses. We also show how the detection of admixture events when running GLOBETROTTER can depend on the surrogate populations used, in a manner consistent with our observation of how inferred admixture dates differ between the “Ethiopia-internal” and “Ethiopia-external” analyses in the real Ethiopian data.

Specifically, we generated simulated admixed populations by mixing two populations A and B that consisted of the following individuals:

A: BedouinA (25 individuals), Brahui (20), Egyptian_Comas (11), Palestinian (30), all from Lazaridis et al 2014

B: Baganda (96 individuals) from Gurdasani et al 2016

This admixture scenario is meant to reflect mixture between an East African source (B) and a West Eurasian source (A), as is seen in many of our Ethiopian populations. To simulate each admixed haploid genome, we employed both the tract-length generation technique described in Price et al 2006 and the forwards-in-time simulation approach described in Hellenthal et al 2014. For the former, we generated each admixed haploid as a sequence of tracts, with tract sizes in centimorgans sampled from an exponential distribution with rate equal to the time (in generations ago) that admixture occurred. Each tract was copied intact from a single source population haploid randomly selected according to the simulated admixture proportions. For the latter, in each generation we simulated each haploid genome as a mosaic of tracts from two randomly selected parent haploids from the previous generation, with tract sizes based on the build 37 recombination map used for the real data analyses. Here the number of tracts per chromosome is equal to a B1 +

B2 + 1, where B1 = {0,1} with probability {0.5,0.5}, and B2 is a random sample from a Poisson distribution with rate equal to the total Morgan length of the chromosome minus 0.5. B1 models the expected obligate crossover per generation on a chromosome, and B2 models the remaining crossovers.

We generated four populations of admixed individuals, where *g* below refers to the admixture time in generations ago and *p* refers to the proportion of DNA inherited from population A:

(1) **"40% (Exp)"**: Admixture with *g*=75 and *p*=0.4, with exponential growth. To do so, we first generated 200 admixed haploids by intermixing populations A and B with an admixture date of 30 generations and *p*=0.4 using the Price et al 2006 technique. We then forwards-in-time simulated this admixed population for 45 generations, assuming a constant population size of 1000 haploids. This reflects an instantaneous growth from 200 to 1000 haploids at generation 1.

(2) **"40% (BN)"**: Admixture with *g*=75 and *p*=0.4, with a severe bottleneck. To do so, we used the same population of 200 admixed haploids generated for (1), and forwards-in-time simulated this admixed population for 45 generations assuming a constant population size of 100 haploids. This reflects an instantaneous decline from 200 to 100 haploids at generation 1.

(3) **"20% (no Dem)"**: Admixture with g=30 and p=0.2, with no additional demography. I.e we generated 200 admixed haploids by intermixing populations A and B with an admixture date of 30 generations and *p*=0.2 using the Price et al 2006 technique.

(4) **"26% (3-date)"**: Admixture at *g*=10, with *p*=0.26 and multiple dates of admixture. To do so, we generated 40 admixed haploids using the Price et al 2006 technique, with 30% of the DNA derived from a population consisting of 160 (out of the total 200) simulated haploids from (1), and the remaining 70% from a population consisting of 160 of the simulated haploids from (3). Thus *p* = 0.3*0.4 + 0.7*0.2 = 0.26. Note that this admixed population has three pulses of admixture among its simulated ancestors: at *g* = 10 between sources (1) and (3), at *g* = 10+30 = 40 between sources A and B due to the admixture in (3) accounting for 70% of the ancestry, and at *g* = 10+75 = 85 generations between sources A and B due to the admixture in (1) accounting for 30% of the ancestry.

For analyses of these simulations, for each of (1)-(4) we made 20 diploid simulated individuals that consisted of 40 randomly selected (without replacement) simulated haploid genomes that were randomly paired. We performed analogous CHROMOPAINTER, SOURCEFIND and

GLOBETROTTER analyses to those performed in the “Ethiopia-internal” and “Ethiopia-external” analyses of the real data. In particular, for the simulations' “Ethiopia-external” analogue, we used CHROMOPAINTER to paint each simulated individual against 259 donor populations that included all of those used in the real “Ethiopia-external” analysis but excluding the five populations in A and B (i.e. $K$=259 here). For the simulations' “Ethiopia-internal” analogue, we painted each simulated individual against these same 259 donor populations plus the four simulated groups comprising the 80 sampled individuals from (1)-(4) (i.e $K$=263 here). (We note that the sampled individuals for (1) and (3) did not include any of the 320 haplotypes used to simulate (4).) Mimicking our real data “Ethiopia internal” analysis, we excluded matching to individuals from the same labeled group when generating painting samples files for each simulated population (1)-(4) for the simulations' “Ethiopia-internal” GLOBETROTTER analogue. For all CHROMOPAINTER analyses of simulated data, we used E-M estimated values of the global mutation (-M) and switch rates (-n) from the “Ethiopia-external” real data analysis.

To ease computational burden, all surrogate individuals used in our GLOBETROTTER/SOURCEFIND simulation analyses were represented by slightly modifying the “Ethiopian-external” paintings we had already generated for the real data analyses. In particular, as this real data analysis painting had allowed each surrogate to match to the five populations from A and B, we set any such matching to 0 (rather than re-painting). Furthermore, for the simulations' “Ethiopia-internal” analogue, we assumed each surrogate population matched 0 to the simulated individuals from (1)-(4). This could potentially lead to a reduction in power for our SOURCEFIND and GLOBETROTTER analyses, but we show that it seems to have made little difference here.

For the “Ethiopia-external” GLOBETROTTER analogue, each of the four populations used 116 surrogate present-day groups, mimicking our real data version of the “Ethiopian-external” GLOBETROTTER analysis but excluding 11 ancient groups and the present-day groups used to simulate. For the “Ethiopia-internal” GLOBETROTTER analogue, each of the four populations used 119 surrogate groups, consisting of these 116 groups plus the three other simulated populations. For the “Ethiopia-external” analogue, we also applied SOURCEFIND to each simulated population, again mimicking our real data analysis but removing the five groups used to simulate and hence using 271 surrogates in total.

Our simulation results are summarized in Figure S5. Simulations (1) and (2), which have the same ancestry before splitting 45 generations ago, are very genetically different under the

“Ethiopia-internal” analogue, e.g. with simulation (1) more similar to simulations (3) and (4) (Fig S5b, top left). However, this difference disappears under the “Ethiopia-external” analogue, with these two populations more similar to each other than any other pairing of the four simulated populations (Fig 4b, bottom right). This mimics our observations in the real data for the caste-like occupational groupings among the Ari and Wolayta (Fig 2a), and other marginalized groups (Chabu, Manja, Manjo, Negede-Woyto), indicating how the “Ethiopia-external” analysis can be used to uncover recent shared ancestry that has been masked by recent endogamy effects. On the other hand, under both analyses simulations (1) and (2) are genetically differentiated from simulations (3) and (4) that have different proportions of admixture from non-Ethiopian sources.

Using the “Ethiopia-external” analogue painting, SOURCEFIND accurately infers the admixing sources and proportions for each simulation (Fig S5d), despite removing the original admixing sources from analysis. Out of 271 surrogate populations, only 9 contribute >1% to any of the four simulated populations: Uganda_Muganda (unpublished Baganda newly released in this study), Egyptian_Metspalu, Balochi, Syria, Jordanian, Makrani, PalestinianArabs, Yemen, Lebanese (Fig S5c). Furthermore, under this “Ethiopia-external” analogue GLOBETROTTER's inferred admixture dates closely match the truth for simulations (1)-(3) (Fig S5d). Notably, simulations (1) and (2) show very similar inferred proportions, sources and dates, reflecting their common recent ancestry and in particular consistent with them having split more recently than their common inferred admixture date of 65-85 generations ago. Under the “Ethiopia-internal” analogue, inferred dates for simulations (2) and (3) closely match the truth, while GLOBETROTTER failed to detect admixture in simulation (1), presumably due to masking since its ancestry patterns are similar to those in simulation (4).

For simulation (4), the inferred admixture date for the “Ethiopia-external” analogue reflects the admixture inherent in simulated population (3), which contributed 70% of the ancestry for simulation (4) (Fig S5d). In contrast, the inferred date for simulation (4) under the “Ethiopia-internal” analogue captures the admixture between simulated populations (1) and (3) (Fig S5d). This reflects our observations in the real data of more recent inferred dates under the “Ethiopia-internal” analysis relative to the “Ethiopia-external analysis” (Fig 4A), indicating how the former can capture intermixing among Ethiopian groups that is missed by the latter, because only the “Ethiopia-internal” analysis includes Ethiopian surrogate groups. The oldest admixture date of 85 generations in simulated population (4), i.e. the admixture inherent in simulated population (1) that contributed only 30% of the ancestry to (4), is missed under both analyses, indicating that older events may be masked by more recent ones in our analyses.

### SI section 4. The association of genetic similarity and language classifications

This section provides further insights into the association between genetics and linguistic classifications. Our study contained Ethiopian individuals from ethnic groups classified as belonging to the Nilo-Saharan (NS) and Afroasiatic (AA) language families. These are classified into four different within-family branches (www.ethnologue.com): the NS Satellite-Core (179 individuals after quality control), AA Cushitic (383 individuals), AA Omotic (536 individuals) and AA Semitic (96 individuals) branches. In addition, our study included 20 individuals from two linguistic isolates (NegedeWoyto, Chabu) not classified into these families (www.ethnologue.com). Genetic differences among individuals from these different categories are summarized in Fig 1b, Fig 2b, Fig S8 and Fig S9. In this section we describe genetic similarities between individuals of different sub-branch classifications, which are summarized in Fig S9 and Extended Tables 7-8.

We had individuals representing two distinct sub-branches within each of the AA Cushitic, AA Omotic and NS Core-Satellite branches, as well as additional classifications within most of these sub-branches (see Extended Table 1). Sub-branches within each of the above are genetically differentiable (p-val < 0.001) under both the "Ethiopia-internal" and "Ethiopia-external" analyses (Fig S9). Therefore on average people from the same language sub-branch classification (i.e. to the third tier of classification provided in www.ethnologue.com) share more recent ancestry with each other than they do with people from other language classifications, with these effects not solely resulting from recent isolation.

We also find that individuals from different linguistic classifications within each sub-branch can be significantly more genetically similar, though some genetic patterns are not consistent with linguistic classifications (Fig S9). For example, within the AA Cushitic East sub-branch, it has been suggested that individuals from Highland and Lowland linguistic categories diverged before 3,000BCE, and that Werizoid (or Dullay) languages diverged from other Lowland languages between this time and 1,000BCE (Ehret, 1976). Conflicting with this, on average Highland speakers are more genetically similar to individuals from particular Lowland groups than individuals from different Lowland groups are to each other, and Lowland speakers from the Dullay and Konso-Gidolo groups are not differentiable from each other (p-val > 0.05) while each being significantly different from other Lowland groups (Fig S9). However, if linguistic trees correlated with the order in which groups became isolated from one another, these genetic discrepancies could be driven by subsequent factors, such as more recent admixture events shared among Dully and Konso-Gidolo speakers that did not affect other Lowland speakers, as suggested by Black (1975).

Within the AA Omotic North sub-branch, out of six linguistic classifications only individuals from ethnicities speaking Janjer and Gonga languages are genetically distinct from all other classifications under both analyses, while individuals from ethnicities speaking the other four languages (Chara, Dizoid, Gimira, Ometo) are not always distinguishable from one another (p-val > 0.05) particularly under the "Ethiopia-external" analysis (Fig S9). All three language classifications within the AA Omotic South sub-branch are genetically distinguishable under the "Ethiopia-internal" analysis (p-val < 0.001) but not all are under the "Ethiopia-external" analysis (Fig S9), suggesting some differences may be attributable to recent isolation.

Similarly, all three language classifications (Surmic, Nilotic, Koman) within the NS Core sub-branch are genetically distinguishable under the "Ethiopia-internal" analysis (Fig S9). The Gumuz have a disputed language classification, B'aga in Ethnologue, but in Bender (1976) it is suggested that the Gumuz language may be classified as Koman. Genetically, the Gumuz are significantly most similar to Komo speakers under the "Ethiopia-external" analysis (Fig S9, Fig S8b, Extended Table 4), suggesting they share more recent ancestry with Koman speakers than with other NS Core sub-classifications included in this study. Out of these NS Core sub-branches, the linguistically-unclassified Chabu are most similar to the Koman speakers and Gumuz (Fig S9, Extended Table 4) rather than the Surmic classification that contains the Mezhenger. However, this likely reflects the relatively high degree of genetic variability among the multiple sampled Surmic groups.

### SI Section 5. Inferring recent ancestry/admixture in Ethiopian groups

In this section we provide further details of the SOURCEFIND analysis used to infer ancestry and GLOBETROTTER analyses used to identify and date admixture events in the Ethiopian groups.

#### Inferring recent ancestry and dating admixture by comparing Ethiopian clusters to non-Ethiopian groups ("Ethiopia-external" analysis)

We used SOURCEFIND (Chacón-Duque et al., 2018) to compare haplotype sharing patterns (inferred by CHROMOPAINTER under the "Ethiopia-external" analysis) in each Ethiopian target cluster to those in 264 present-day world-wide reference populations (Table S2a, with a subset shown in Fig 3, top-left) and 11 ancient populations (Table S2b): the ~4.5kya East_Africa_forager Ethiopian Mota, plus Anatolia_Neolithic, South_Africa_Stone_Age, Iberia_Chalcolithic, Iberia_Neolithic, Iranian_Neolithic, Karasuk, LaBrana, Loschbour_Hunter_Gatherer, Srubnaya, WC1, Yamnaya. We excluded six African ancient groups, including the South_Africa_Iron_Age and West_Africa_Stone_to_Metal_Age groups that were inferred not to contribute in earlier analyses, plus the Prendergast et al 2019 samples as described in the next paragraph. Using a Bayesian approach, SOURCEFIND infers how best to describe the sharing patterns in the target group as a mixture of those of the reference populations. This ancestry inference is summarized by the barplots in Fig 3, Fig S12 and in Extended Table 5. For the 68 clusters where we infer admixture using GLOBETROTTER, only 13 out of 275 reference populations contributed >5% to any Ethiopian cluster under SOURCEFIND: the 4,500-year-old Ethiopian Mota and 12 present-day groups: Baganda from Uganda, Chad_Bulala, Dinka from Sudan, Egyptian_Metspalu, Kenya_Elmolo, Kenya_Rendille, Kenya_Sengwer, Saudi, Somali, Tanzania_Iraqw, Uganda_Muganda and Yemen.

We caution that this ancestry composition is not implying that each Ethiopian group is a mixture of these reference populations, or that the total proportions contributed by the 2-3 sources per cluster outlined in Fig 3 and Fig S12 accurately reflect the proportion of DNA inherited from those sources (though they can do – see simulations in SI section 3). Instead Ethiopian groups carry haplotype patterns that match those carried in these reference populations, suggesting more recent shared ancestry with those reference populations relative to the other reference populations. There are two important caveats/limitations of our approach. The first is that comparing to different reference populations potentially can give quite different results, and it is unclear what the “best” reference population set is. Our choice of reference populations is informed by findings reported by Prenderagast 2019 that studied ancestry in East Africans (including Ethiopians), in that we included surrogates related to the four primary sources of ancestry they detected, i.e. populations from West Eurasia (representing what Prendergast et al 2019 refer to as “EN2” in their Figure 3), the Sudanese Dinka (“EN1” in their Figure 3), the 4.5kya Ethiopian “Mota” (“E.African forager-related” in their Figure 3) and Bantu-speaking groups (“W.African-related” in their Figure 3). We excluded the aDNA samples from Kenya and Tanzania reported in Prendergast et al 2019 in our SOURCEFIND analysis, because (i) the authors inferred each of those aDNA samples to be admixed descendents of the four primary sources mentioned above and (ii) to enable comparison to the Prendergast et al 2019 findings (e.g. their Figure 3).

A second caveat of our SOURCEFIND analysis is that five of the 12 present-day groups contributing >5% (Chad_Bulala, Kenya_Elmolo, Kenya_Sengwer, Kenya_Rendille, Tanzania_Iraqw) had only two samples. When painting an individuals' genome using CHROMOPAINTER, an individual cannot match to itself. Thus each of these four populations matched to only one sample from their own population via CHROMOPAINTER, which may mitigate signals of isolation (e.g. due to endogamy) in that population, relative to groups that can match to a greater number of individuals from their own labeled group. By mitigating signals of endogamy effects, such reference populations can potentially be favored as an ancestral source in SOURCEFIND analysis, which aims to find the reference populations with painting patterns that most closely match those of the target (in this case Ethiopian) cluster. This may also explain why Mota is favored, as it has only a single sample and hence no means of measuring endogamy under this approach. Nonetheless, comparisons among Ethiopian clusters are still meaningful when conditioning on this set of references, as each cluster was analysed in the same way. In general, we note that surrogate populations with high degrees of isolation (e.g. due to endogamy) may be less likely to be selected as representative of an ancestral source, which is one way SOURCEFIND likely differs from e.g. a f3 outgroup test (Patterson et al 2012). But arguably such surrogates should be downweighted, as – due to recent isolation – the genetic make-up of such surrogates likely no longer well-reflects the ancestral source population.

We also applied GLOBETROTTER (Hellenthal et al 2014) as described in the main text separately to each Ethiopian cluster in order to identify and date admixture events, under a model that assumes one or two pulses of admixture where two or more sources intermixed. Analogously to SOURCEFIND, GLOBETROTTER uses surrogates to the putative admixing sources. Briefly, first CHROMOPAINTER is used to match haplotype patterns within individuals from the target (i.e. putatively admixed) group to those in a set of reference individuals. In doing so, each target individual's genome is composed of a series of DNA segments, with each segment matched to (i.e. inferred to share a most recent common ancestor with) a specific reference population. GLOBETROTTER then infers admixture in the target group by modelling the decay in linkage disequilibrium among segments within each target individual that match to different surrogate populations using the CHROMOPAINTER results.

For our GLOBETROTTER analysis, we excluded as surrogates 145 present-day and six ancient non-African groups (Anatolia_Neolithic, Iberia_Chalcolithic, Karasuk, LaBrana, Srubnaya, Yamnaya) that did not contribute to any Ethiopian cluster in our SOURCEFIND analysis. Instead as surrogates we included the remaining 119 present-day groups (including all from Africa) and 11 aDNA groups (hence 130 surrogates total), including all eight African aDNA groups and the three non-African groups Iberia_Neolithic, Iranian_Neolithic, Loschbour_Hunter_Gatherer. In general date estimation is robust to the surrogate groups included, so long as a surrogate is not substantially related to the target and hence masks the admixture signal, which is easy to diagnose (Hellenthal et al 2014). For every pairing of surrogate populations, GLOBETROTTER infers the probability that two DNA segments on the same chromosome within a target individual share a most recent ancestor with those two surrogate groups, with one DNA segment matched to each surrogate, versus the centimorgan distance between the two segments. Examples of these probability curves (after some scaling) are provided in Fig S15. Importantly, under the pulses of admixture model assumed by GLOBETROTTER, if two surrogate groups represent the same (unknown) admixing source, the inferred probability for that pair will decrease exponentially with increasing genetic distance. In contrast, if two surrogate groups represent different admixing sources, their inferred probability will increase exponentially with increasing genetic distance (Hellenthal et al., 2014). Therefore, by studying the probability patterns among all pairs of surrogates, GLOBETROTTER can automatically infer the number of admixture events (though attempts only to characterize up to two events in practice; see below), as well as which surrogates best match genetically to each putative source involved in each event.

For each Ethiopian cluster, GLOBETROTTER assigns any inferred admixture to one of four categories: (i) “one-date” involving a single purse of admixture between two sources, (ii) “one-date-multiway” involving a single pulse of admixture between greater than two sources, (iii) “multiple-dates” involving more than a single pulse of admixture at different times, potentially between greater than two sources, and (iv) “uncertain” where the probability curves described above are challenging to categorize into (i)-(iii). For (iii), GLOBETROTTER attempts to only date and describe two distinct pulses of admixture, though we note these signals can be consistent with more than two pulses of admixture or continuous admixture (Hellenthal et al., 2014). Furthermore, signals concluding (i),(ii),(iv) may reflect a failure of GLOBETROTTER to identify genuine older admixture events and/or have inferred dates biased towards more recent admixing in the case of continuous or multiple admixture events (Hellenthal et al., 2014), as illustrated in our simulation results (Fig S5).

We also visually inspected the probability curves (e.g. Fig S15) to assess whether the conclusions (i)-(iv) that GLOBETROTTER reports fit the data. Based on this visual inspection, and using the parameters highlighted below from the GLOBETROTTER *main.txt output files, we made some slight alterations to GLOBTROTTER's reported conclusions. In particular, to be conservative we do not report GLOBETROTTER results for clusters where “r2.oneevent”, which assess the overall evidence of admixture (on a 0-1 scale), was <0.34, as such clusters had noisy probability curves. Also, for the “Ethiopia-internal' analysis, which is picking up more subtle admixture between genetically similar groups (i.e. between Ethiopian groups), we did not analyse clusters with <=5 individuals, and we do not report results for two clusters (Eth_al, Eth_cr) with “r2.oneevent” < 0.5 that had noisy probability curves. In addition to these omissions, we slightly altered GLOBETROTTER's default threshold for concluding “one-date, multiway” over “one-date” from “fit.1event < 0.975” to “fit.1event < 0.98”, which changed the conclusion from “one-date” to “one-date, multiway” for four clusters (Eth_ab, Eth_ap, Eth_ar, Eth_bh) in the “Ethiopia-external” analysis and one cluster (Eth_ax) in the “Ethiopia-internal” analysis (Extended Table 5-6). Finally, we visually inspected clusters for which “maxScores.2events” ε (0.3,0.35], which is indicative of multiple-dates of admixture but does not meet GLOBETROTTER's default criterion of “maxScores.2events” > 0.35 for concluding “multiple-dates”. In some of these cases, two admixture dates appeared to fit the data notably better than one date; i.e. the red line in the GLOBETROTTER *pdf file output was a better fit to many probability curves relative to the green line. Thus we changed the conclusion from “one-date” of admixture to “multiple-dates” of admixture for three clusters (Eth_ag, Eth_ak, Eth_as) under the “Ethiopia-external” analysis and for three clusters (Eth_ag, Eth_ak, Eth_bi) under the “Ethiopia-internal” analysis (Extended Table 5-6). We note that other clusters may have multiple dates of admixture that we miss here, and that more data from Ethiopians will help to clarify these admixture signals in the future.

For each Ethiopian cluster for which GLOBETROTTER infers admixture, we use 100 bootstrap re-samples of individuals to infer confidence intervals around inferred dates.

GLOBETROTTER infers admixture events in 68 of the 78 Ethiopian clusters, with dates ranging from ~100 to 4200 years ago. The type of admixture events inferred among these 68 include 31 “one-date”, 27 “one-date-multiway” and ten “multiple-dates”. Based on visual inspection of each cluster's GLOBETROTTER probability curves, i.e. the probability that two DNA segments separated by increasing centimorgan distance are matched to two particular surrogate groups as described above (see Fig S15), we determined that admixing sources broadly could be defined by contribution patterns from the following six reference groups:

- the Nilotic-speaking Dinka from Sudan  (sometimes including the Bulala from Chad)
- the Nilotic-speaking Sengwer from Kenya
- the Bantu-speaking Baganda from Uganda
- the 4.5kya Ethiopian Mota
- the Cushitic-speaking Rendille from Kenya (sometimes including Somalians)
- Egyptians and two West Eurasian groups (Saudi Arabia, Yemen)

This is consistent with our SOURCEFIND results (Fig 3, Fig S12, Extended Table 5), for which these groups were the highest contributors to inferred ancestry. Various combinations of these different groups define the six admixture signatures we report in Fig 3 and Fig S12:

(1) A cluster containing the NS speaking Murle and Nyangatom shows multiway admixture at one date (10-16 gen ago) betweent three distinct sources best represented by the Muganda, Dinka and Sengwer.

(2) Seven clusters of NS speakers, plus separate clusters of the AA Omotic speaking Karo and linguistically-unclassified Chabu, show relatively recent admixture (typically < 30 gen ago) between two sources best represented by Mota and Dinka/Sengwer.

(3) Four clusters of AA speakign groups show admixture between two sources represented by Rendille and either Sengwer (seen in the Dasanech and Arbore clusters) or Mota (seen in the Manja and Manjo clusters).

(4) Three clusters containing primarily AA Omotic-speaking Hamar, AA Cushitic-speaking Tsamay, or NS speaking Meinit each show multiple sources and/or dates of admixture between three sources represented by Sengwer, Mota and Rendille.

(5) Thirty clusters containing primarily AA Cushitic and AA Omotic individuals show multiway admixture, at a wide range of inferred dates consistent with continuous or multiple pulses of admixture, between three sources represented by Egypt, Rendille and Mota.

(6) Twenty-one clusters show intermixing of two sources by Egypt versus Mota/Rendille, at either one or two separate admixture dates, with two dates consistent with continuous or multiple pulses of admixture. Among these, a cluster of the linguistically-unclassified Negede Woyto show similar inferred ancestry to the 18 of these 21 clusters consisting almost exclusively of AA speakers. The remaining three clusters contain NS speaking groups (Berta, Nyangatom) and show different patterns that include a substantial amount of Dinka-like ancestry. Consistent with their geography, the five of these 21 clusters containing AA Semitic-speakers and the AA Cushitic-speaking Agaw, plus two clusters containing the AA Cushitic-speaking Qimant and AA Omotic-speaking Shinasha, show the highest amounts of Egypt-like ancestry, with inferred admixture dates spanning 59-100 gen ago.

Though using different surrogates and techniques complicates direct comparisons, our inferred sources of ancestry broadly agree with those reported for present-day Ethiopian groups by Prendergast et al 2019, in particular results reported in their Figure 3. For example, clusters containing the Agaw (clusters 66, 67 in Fig 3, Fig S12) have relatively more Levant-like ancestry (which we represent with matching to Egypt), the Ari (clusters 22, 24, 25; called Aari in Prendergast et al 2019) have relatively more Mota-like ancestry, and the Mursi (cluster 2) have relatively more Dinka-like ancestry. Also consistent with their results, in general we find NS-speakers to have more

Dinka-related ancestry than AA speakers. Furthermore, we infer mixture between Mota-like and Levant-like sources as old as around 4000 years ago among some AA speaking populations, though we note our study of present-day populations may miss some of the older admixture events reported in that study.

Exploring recent admixture among Ethiopian clusters (“Ethiopia-internal” analysis)

We next set to determine whether there has been intermixing among Ethiopian groups. To do so, we applied GLOBETROTTER to each Ethiopian cluster, using 64 Ethiopian clusters as potential surrogates for the admixing sources in addition to the 130 surrogates used in the “Ethiopia-external” analysis. Fourteen of the 78 Ethiopian clusters (marked by asterisks in the first column of Extended Table 2) were not included as surrogates, because they each contained small numbers of individuals from several ethnic groups that hence would confuse interpretation of results. These GLOBETROTTER analyses inferred admixture in 61 of the 78 Ethiopian clusters, with 32 “one-date”, 19 “one-date-multiway”, 6 “multiple-dates” and 4 “uncertain” events. The inferred dates of events were much more recent relative to the analysis that excluded Ethiopian surrogates (Fig 4a). Overall 43 (84.3%) of 51 groups that concluded “one-date” and “one-date-multiway” events had inferred point estimate dates <30 generations ago (~750-850 ya) under this analysis. This demonstrates how this analysis is capturing more recent intermixing by including Ethiopian surrogates, likely because different Ethiopian groups have been intermixing more recently.

To assess whether Ethiopian groups are intermixing with geographically nearby groups, we first arranged our clusters (and the 4.5kya Ethiopian Mota) along a line (represented by the circle in Fig 4b) based on the geographic distance between them. To do so, we ordered groups according to their order along the first component of a principal-components-analysis (PCA) of the geographic (Haversine) distance matrix between clusters, where the latitude/longitude of each cluster is the average of that among all individuals within that cluster. For each of the 57 clusters that did not infer “uncertain” admixture, we took the GLOBETROTTER inference of the most strongly signalled event (“firstevent” in Extended Table 6). This is the event that explains the most variation in the set of GLOBETROTTER probability curve pairings across all reference populations (examples provided in Fig S15), i.e. as determined using the first (main) principal component of this probability curve set (see Hellenthal et al 2014 for details). (For clusters where admixture between more than 2 sources is inferred, the second, less clearly inferred admixture event is only partially captured by this first principal component if at all.) This event infers that two distinct sources intermixed, one contributing a majority of ancestry and the other contributing a minority of ancestry to the cluster's individuals.  We calculated two "geographic proximity scores" for each cluster by finding the ordinal distances, along the PCA-line, between that cluster and (A) the surrogate group that GLOBETROTTER inferred as the best genetic match to the minority-contributing source and (B) the surrogate group that GLOBETROTTER inferred as the best genetic match to the majority-contributing source. For each of (A) and (B), we did not include clusters where the surrogate was a non-Ethiopian group, and we averaged scores across all included clusters. This gave final values of 14.02 and 9.60 for (A) and (B), respectively. To be conservative, we took the highest of these two scores, i.e. that based on the ordinal distance between the cluster and the minority-contributing source. The lower score of (B) makes intuitive sense, as typically the majority contributing source is more genetically similar and geographically closer to the target group than the minority contributing source. Indeed, for this reason, the majority source is often presumed to reflect the ancestors who lived in the same region as the present-day target individuals, while the minority source is presumed to have admixed with these ancestors, e.g. after migrating into the region. We then permuted labels around the line 50,000 times, and recalculated our average proximity score (i.e. to the same minority contributing source labels) for each permutation. The permutation-based p-value calculating the proportion of permutations whose average proximity score was less than or equal to that of the observed average proximity score was highly significant ($p\text{-val} < 0.0002$). Overall these results suggest that geographically nearby groups in Ethiopia have intermixed with each other more recently than the W.Eurasian and W.African-source events inferred in our admixture analysis that excludes Ethiopians as surrogates.

<u>Alternative approach to explore recent admixture among Ethiopian clusters used in Figure 5</u>

Very recent admixture may be challenging to detect via GLOBETROTTER, which e.g. has no power to see admixture occurring one generation ago. Therefore, for each of these six cultural practices shown in Fig 5, we used an alternative means of assessing whether there was evidence of recent intermixing among people from pairs of groups that both reported the given practice. To do so, we exploit the fact that if two groups have recently intermixed, and/or if some individuals from one group have taken on the label of the other group, it is expected that some -- but not all -- pairings of individuals from the two groups will share a most recent common ancestor (MRCA) for atypically long stretches of DNA. Using this idea, the green colors in the upper left triangles of the heatmaps in Fig 5 indicate whether at least some (>=1) pairings of individuals, one from each group, have average inferred MRCA segments (inferred by CHROMOPAINTER under the "Ethiopia-internal" analysis) that are >2.5 cM longer than the median length of average inferred MRCA segments across all such pairings of individuals from the separate groups. For comparison, we see such a trend in 122 (11.8%) of 1035 (= 46 choose 2) total pairings of groups considered in this analysis, versus in 10 (18.2%) of 55 pairings of the 11 groups that reported practicing male circumcision, 5 (23.8%) of 21 pairings of seven groups that reported Sororate/Cousin marriages, and 2 (66.7%) of the 3 pairings of three groups that reported female circumcision. In Table S11, we report the proportion out of 10,000 random samples of 3, 7 or 11 groups (sampled from the 46 SNNPR groups analysed here) where a greater or equal number of group pairings showed this trend, also considering various different values of excess average MRCA segment size.

### SI section 6. Descriptions of cultural traits

The practices listed below are reported in (The Council of Nationalities, Southern Nations and Peoples Region, 2017), with groups' reports regarding them provided in Extended Table 10.

<u>Arranged marriage or marriage arranged by parents</u>: marriage arranged by the parents of the couple with little or no involvement of the couple.

<u>Abduction</u>: involves a man, often assisted by members of his family and/or friends, forcibly taking a young girl or a mature woman as a wife. No parental consent is obtained.

<u>Swift/spontaneous unions</u>: said to occur only occasionally in southwestern parts of Wollo province to the northeast of the south-westerly bend of the Blue Nile but also elsewhere including in the SNNPR (e.g. among the Halaba). Involves a girl, well-past normal marriageable age and spur of the moment agreement.

<u>Sororate/cousin marriages</u>: a widower marries a sister or cousin of his deceased wife.

Wife-replacement: a widower marries a sister or close female relative of his deceased wife.

Wife-inheritance: a married man "inherits" i.e. takes as an additional wife a widow of his deceased elder brother, cousin or close kinsman with the primary objective of providing trusteeship for children and assets left behind by his deceased relative.

Belt-giving: a form of marriage that involves offering the intended bride ladies' belts as a symbolic gesture of the young man's desire to marry her. If the girl does not wish to marry the man she refuses to accept the belt. The belt may be considered a token of love and may form a small portion of the bride-price. Parents may not be able to object to the marriage.

Bead-giving: a young man offers his future bride beads as a token of his love and affection. A variant of this practice involves a young man forcibly tying beadwork round a girl's neck despite her resistance (parents may not be able to object to marriage following such an event). The beads may be considered a small pre-marriage instalment of bride-price.

Beaded necklace snatching: a form of marriage that involves an earlier snatching of a young girl's beadwork necklace. (The act of snatching the beadwork is a symbolic gesture of the man's wish to marry the woman.) If adult male members of the girl's family apprehend the snatcher, he may be beaten and dispossessed of the bead necklace, in which case he cannot lay customary claim to the girl as his future wife. If the man escapes with the girl's bead necklace the girl's family will be forced to give the girl up to the man for marriage.

Note: many groups in the SNNPR accord ideological/symbolic significance to belts and beads that revolves around 'omens and a person's fate and fortunes'.

Men moving in to marry women: the married couple move into the bride's home after marriage. This is uncommon. A women most commonly moves into a home built and owned by her husband.

Women moving in to marry men: suddenly (in circumstances in which a man and woman are in love), completely unannounced, a young woman enters the family home of a young man and, clinging to the central pole of the house, pleads with the boy's parents to allow their son to marry her.

Repeat marriages: marriages following divorce or annulment of a previous union.

Bride’s butter anointment: an important part of a marriage ceremony that takes place at the groom’s family home during a wedding. It involves the future mother-in-law anointing the hair of the bride with a generous amount of butter.

Male circumcision: removal of the foreskin. (May be performed on babies, young boys, teenagers and adult males, individually or in groups of similar age. May be part of initiation ceremonies.)

Female circumcision: cutting of the labia minora and/or the labia majora, with or without excision of the clitoris of young girls and women. In northern Ethiopia it is performed at an early age, while in the south-western parts of the country it takes place at a later age and is related to marriage.

Pre-marital sex: sexual intercourse with the opposite sex prior to marriage (considered unacceptable in most communities in Ethiopia but accepted as the norm by a few groups in the SNNPR).

Pre-marital pregnancy or birth: a woman becoming pregnant or having a baby prior to a marital union.

Endo/exogamy: marriage of a man or a woman to the opposite sex, respectively, within/outside their clan or lineage.

Poly/monogamy: marriage of a man to many wives or just one wife, respectively.

Marriage with bride’s parental consent: marriage of a woman to a man with her parents’ consent.

Marriage with brides encouraged by parents: marriage of a woman to a man whom the woman’s parents prefer to other men.

Groom’s parental choice marriage: marriage of a man to a woman whom the man’s parents prefer to other women.

Groom's aunt arranged marriage: marriage of a man to a woman whom the man's aunt prefers to other women.

Marriage involving third-party agents: marriage between a man and a woman arranged by third-party agents (might be cousins, aunts, uncles, friends, acquaintances, any other person, related or unrelated, to the couple).

Marriage involving women intermediaries: marriage between a man and a woman arranged solely by female intermediaries (they may or may not be related to the couple).

Special unions: marriage unions between a man and a woman that do not conform to commonly accepted cultural practice followed by most members of a group (important examples of such marriages include community or religious/spiritual leaders, chiefs and kings).

Minors' marriage: marital union between underage children of the opposite sex arranged by their parents.

Mate-selection: marriage in which the couple themselves decide to marry (with little involvement of either set of parents in marriage negotiations).

Marriage with mutual spouse consent: as mate-selection.

Marriage with spouse and parental consent: marriage of a man and a woman with the consent of the couple and their parents.

Marriage by elopement/persuasive absconding: marriage of a man and woman usually without parental consent after the man persuades the woman to abscond with him.

### Supplementary Tables

**Table S1** Features of the 1,214 Ethiopian samples included in this work, including ethnicity, language group and self-reported first and second languages (n = sample size). “Language group” is given at the first (“Nilo-Saharan”) and to the second (other languages) classification level at www.ethnologue.com; all Nilo-Saharan speakers are from the “Satellite-Core” second classification. Labels in parentheses of column 1 are used in Fig 5; labels in parenthesis of column 3 are used in Fig S7ab. Groups from the SSNPR with information about culture practices are indicated with an asterisk.

| Ethnicity | Language group | Self-reported 1st language | Self-reported 2nd language | n |
|---|---|---|---|---|
| Afar (AFA) | Afroasiatic_Cushitic | Afar (AF) | Amharic | 9 |
| | | | Tigrinya | 1 |
| Agaw (AGA) | Afroasiatic_Cushitic | Agaw (AG) | Amharic | 14 |
| | | Amharic (AM) | Agaw | 6 |
| | | | None | 10 |
| | | unknown | unknown | 2 |
| Alae (ALA) | Afroasiatic_Cushitic | Alae (GB) | Amharic | 16 |
| | | | Bana | 1 |
| | | | Affa Honso | 7 |
| | | | None | 8 |
| | | | Oromiffa | 2 |
| | | Masholae (MH) | Oromiffa | 5 |
| | | Oromiffa (OR) | Amharic | 3 |
| | | | Alae | 3 |
| | | | Masholae | 1 |
| Amhara (AMH) | Afroasiatic_Semitic | Amharic | None | 25 |
| | | | Oromiffa | 2 |
| | | Oromiffa | Amharic | 1 |
| | | unknown | unknown | 26 |

| | | | | |
|---|---|---|---|---|
| Anuak (ANU) | Nilo-Saharan | Anuak (AN) | Amharic | 9 |
| Arbore (ARB) | Afroasiatic_Cushitic | Arbore (AE) | Amharic | 4 |
| | | | None | 3 |
| | | | Oromiffa | 7 |
| Ari (Aari) (ARI) | Afroasiatic_Omotic | unknown | unknown | 2 |
| Ari_Cultivator (ARI_C) | Afroasiatic_Omotic | Ari (AA) | Amharic | 9 |
| | | | None | 5 |
| Ari_Potter (ARI_P) | Afroasiatic_Omotic | Ari | Amharic | 4 |
| | | | Bana | 1 |
| | | | None | 19 |
| Ari_Smith (ARI_S) | Afroasiatic_Omotic | Ari | Amharic | 4 |
| | | | None | 10 |
| Bana (BAN) | Afroasiatic_Omotic | Amharic | Bana | 1 |
| | | Bana (BE) | Ari | 4 |
| | | | Amharic | 14 |
| | | | Hamar | 1 |
| | | | None | 8 |
| Basket (BAS) | Afroasiatic_Omotic | Amharic | Basket | 1 |
| | | Basket (BK) | Amharic | 13 |
| Bench (BEN) | Afroasiatic_Omotic | Bench (BN) | Amharic | 9 |
| | | | Kafacho | 2 |
| | | | None | 1 |
| Berta (BER) | Nilo-Saharan | Berta (BT) | Amharic | 8 |
| | | | Oromiffa | 3 |
| | | | unknown | 2 |
| BetaIsrael (BET) | Afroasiatic_Semitic | unknown | unknown | 13 |
| Bodi (BOD) | Nilo-Saharan | Bodi (BD) | Amharic | 9 |
| | | | None | 5 |
| Burji (BUR) | Afroasiatic_Cushitic | Burji (BR) | Amharic | 2 |
| | | | None | 3 |
| | | | Oromiffa | 10 |
| | | Oromiffa | Amharic | 8 |
| | | | Burji | 1 |
| Chabu (CHA) | Unclassified | Chabu (CH) | Kafa/Shekacho | 11 |
| Chara (CHR) | Afroasiatic_Omotic | Chara (CR) | Kafacho | 17 |
| Dasanech (DAS) | Afroasiatic_Cushitic | Dasenech (DS) | Amharic | 7 |
| | | | None | 8 |
| Dawro (DAW) | Afroasiatic_Omotic | Dawro (DW) | Amharic | 13 |
| | | | None | 1 |
| Dhime (DHI) | Afroasiatic_Omotic | Dhime (DM) | Amharic | 11 |

| | | | | |
|---|---|---|---|---|
| | | | Bodi | 9 |
| | | Gofa (GF) | Amharic | 1 |
| Dirasha (DIR) | Afroasiatic_Cushitic | Amharic | Dirasha | 2 |
| | | Dirasha (DR) | Amharic | 10 |
| | | | Oromiffa | 1 |
| | | Oromiffa | Amharic | 3 |
| | | | Dirasha | 1 |
| Dizi (DIZ) | Afroasiatic_Omotic | Amharic | Dizi | 3 |
| | | Dizi (DI) | Amharic | 10 |
| | | | None | 1 |
| Dorze (DOR) | Afroasiatic_Omotic | Dorze (DZ) | Gamo (other) | 15 |
| Gedeo (GED) | Afroasiatic_Cushitic | Gedeo (GE) | Amharic | 21 |
| GentaGamo (GEN) | Afroasiatic_Omotic | Amharic | Gamo (other) | 1 |
| | | | GentaGamo | 2 |
| | | | None | 1 |
| | | Gamo (other) (GM) | Amharic | 2 |
| | | GentaGamo (GN) | Amharic | 4 |
| | | | Gidicho | 2 |
| | | | Gamo (other) | 3 |
| Gidicho (GID) | Afroasiatic_Omotic | Gidicho (GG) | Amharic | 6 |
| | | | GentaGamo | 5 |
| Gofa (GOF) | Afroasiatic_Omotic | Amharic | Gofa | 3 |
| | | | Gamo (other) | 3 |
| | | | None | 3 |
| | | Gofa | Amharic | 2 |
| | | | Gamo (other) | 2 |
| | | Gamo (other) | Amharic | 2 |
| Gumuz (GUM) | Nilo-Saharan | Gumuz (GU) | Amharic | 2 |
| | | unknown | unknown | 20 |
| Gurage (GUR) | Afroasiatic_Semitic | Amharic | Gamo (other) | 1 |
| | | | Gurage | 2 |
| | | | None | 3 |
| | | | Oromiffa | 1 |
| | | Gurage (GR) | Amharic | 7 |
| | | | Sidama | 1 |
| | | Oromiffa | Amharic | 1 |
| Hadiya (HAD) | Afroasiatic_Cushitic | Amharic | Hadiya | 2 |
| | | | None | 1 |
| | | Hadiya (HD) | Amharic | 11 |
| Halaba (HAL) | Afroasiatic_Cushitic | Halaba (AL) | Amharic | 13 |

| | | | | |
|---|---|---|---|---|
| | | Amharic | Halaba | 1 |
| Hamar (HAM) | Afroasiatic_Omotic | Amharic | Hamar | 1 |
| | | Hamar (HM) | Amharic | 7 |
| | | | Karo | 1 |
| | | | None | 5 |
| Honsita (HON) | Afroasiatic_Cushitic | Amharic | Affa Honso | 3 |
| | | | None | 1 |
| | | Affa Honso (KS) | Amharic | 10 |
| | | | Dirasha | 2 |
| | | | Gamo (other) | 1 |
| Kafacho (KAF) | Afroasiatic_Omotic | Kafacho (KF) | Amharic | 16 |
| Kambata (KAM) | Afroasiatic_Cushitic | Amharic | Kambata | 1 |
| | | Kambata (KM) | Amharic | 12 |
| Karo (KAR) | Afroasiatic_Omotic | Karo (KO) | Hamar | 14 |
| Komo (KOM) | Nilo-Saharan | Komo (KE) | Mao | 1 |
| | | | None | 4 |
| | | | Oromiffa | 3 |
| Konta (KON) | Afroasiatic_Omotic | Konta (KN) | Amharic | 16 |
| Kore (KOR) | Afroasiatic_Omotic | Amharic | Kore | 3 |
| | | Kore (KR) | Amharic | 5 |
| | | | Oromiffa | 2 |
| | | Oromiffa | Amharic | 6 |
| Kwegu (KWE) | Nilo-Saharan | Karo | Kwegu | 1 |
| | | Kwegu (KW) | Karo | 6 |
| | | | None | 3 |
| Maleh (MAL) | Afroasiatic_Omotic | Maleh (ML) | Amharic | 9 |
| | | | Gofa | 2 |
| Manja (MAN) | Afroasiatic_Omotic | Manja (MA) | Dawro | 14 |
| Manjo (MAJ) | Afroasiatic_Omotic | Kafa/Shekacho (SC) | Amharic | 11 |
| Mao (MAO) | Afroasiatic_Omotic | Mao (ME) | Oromiffa | 1 |
| | | Oromiffa | Amharic | 5 |
| | | | Mao | 2 |
| | | | None | 1 |
| Masholae (MAS) | Afroasiatic_Cushitic | Alae | Oromiffa | 1 |
| | | Masholae | Amharic | 8 |
| | | | Dirasha | 3 |
| | | | Oromiffa | 2 |
| | | Oromiffa | Amharic | 4 |
| | | | Masholae | 1 |
| Meinit (MEI) | Nilo-Saharan | Meinit (MT) | Amharic | 14 |

| | | | | |
|---|---|---|---|---|
| | | | Bench | 1 |
| Mezhenger (MEZ) | Nilo-Saharan | Mezhenger (MG) | Amharic | 9 |
| | | | None | 5 |
| Mossiye (MOS) | Afroasiatic_Cushitic | Mossiye (BU) | Amharic | 2 |
| | | | Gamo (other) | 2 |
| | | | None | 1 |
| | | Gamo (other) | Amharic | 3 |
| | | | Mossiye | 1 |
| | | | Dirasha | 1 |
| Murle (MUR) | Nilo-Saharan | Nyangatom (BM) | Amharic | 6 |
| | | | None | 6 |
| | | Murle (MB) | Nyangatom | 1 |
| Mursi (MUS) | Nilo-Saharan | Mursi (MU) | None | 10 |
| | | unknown | unknown | 2 |
| Nao (NAO) | Afroasiatic_Omotic | Kafacho | Amharic | 1 |
| | | Nao (NO) | Kafacho | 16 |
| NegedeWoyto (NEG) | Unclassified | Amharic | None | 9 |
| Nuer (NUE) | Nilo-Saharan | Nuer (NR) | Amharic | 2 |
| | | | English | 1 |
| | | | None | 8 |
| Nyangatom (NYA) | Nilo-Saharan | Nyangatom | Amharic | 7 |
| | | | None | 5 |
| Oromo (ORO) | Afroasiatic_Cushitic | Oromiffa | Amharic | 7 |
| | | unknown | unknown | 28 |
| OtherGamo (OTH) | Afroasiatic_Omotic | Amharic | Gamo (other) | 10 |
| | | Gamo (other) | Amharic | 6 |
| Qimant (QIM) | Afroasiatic_Cushitic | Amharic | None | 16 |
| | | | QM | 1 |
| Shekacho (SHK) | Afroasiatic_Omotic | Kafa/Shekacho | Amharic | 16 |
| Sheko (SHE) | Afroasiatic_Omotic | Sheko (SK) | Amharic | 15 |
| Shinasha (SHI) | Afroasiatic_Omotic | Oromiffa | Amharic | 1 |
| | | Shinasha (SA) | Amharic | 16 |
| | | | Oromiffa | 1 |
| Sidama (SID) | Afroasiatic_Cushitic | Amharic | Sidama | 2 |
| | | Sidama (SD) | Amharic | 19 |
| Somali (SOM) | Afroasiatic_Cushitic | Amharic | None | 2 |
| | | unknown | unknown | 24 |
| Suri (SUR) | Nilo-Saharan | Suri (SR) | Amharic | 14 |
| Tigraway (TIG) | Afroasiatic_Semitic | Amharic | None | 2 |
| | | | Oromiffa | 1 |

| | | | | |
|---|---|---|---|---|
| | | | Tigrinya | 2 |
| | | Tigrinya (TG) | Amharic | 8 |
| Tsamay (TSA) | Afroasiatic_Cushitic | Bana | Amharic | 1 |
| | | Alae | Amharic | 1 |
| | | Tsamay (TY) | Amharic | 3 |
| | | | Bana | 12 |
| | | | None | 1 |
| Wolayta (WOL) | Afroasiatic_Omotic | Gamo (other) | Amharic | 1 |
| | | unknown | unknown | 21 |
| | | Wolayta (WL) | Amharic | 3 |
| Wolayta_Cultivator (WOL_C) | Afroasiatic_Omotic | Wolayta | Amharic | 6 |
| Wolayta_Potter (WOL_P) | Afroasiatic_Omotic | Wolayta | Amharic | 1 |
| | | | None | 9 |
| Wolayta_Smith (WOL_S) | Afroasiatic_Omotic | Wolayta | Amharic | 1 |
| | | | None | 11 |
| Wolayta_Tanner (WOL_T) | Afroasiatic_Omotic | Wolayta | Amharic | 2 |
| | | | None | 6 |
| Wolayta_Weaver (WOL_W) | Afroasiatic_Omotic | Wolayta | None | 12 |
| Yem (YEM) | Afroasiatic_Omotic | Yem (YM) | Amharic | 13 |
| Zayse (ZAY) | Afroasiatic_Omotic | Amharic | None | 1 |
| | | | Zayse | 4 |
| | | Zayse (ZS) | Amharic | 11 |
| | | | None | 1 |
| Zilmamo (ZIL) | Nilo-Saharan | Zilmamo (ZL) | Suri | 12 |

**Table S2**. Non-Ethiopian samples included in this work (n = 2,718), for both the “Ethiopia-internal” and “Ethiopia-external” analyses. Reg=Region, with NA=North Africa, WA=West Africa, CA=Central Africa, EA=East Africa, SO=Somalia, SA=South Africa, EG=Egypt, WE=West Eurasia, SS=South Asia, ES=East Asia, CS=Central Asia/Siberia, OC=Oceania, AM=Americas

(a) present-day individuals:

| Reg | Population label | n | Reference | Reg | Population label | n | Reference |
|---|---|---|---|---|---|---|---|
| AM | Algonquin | 8 | Lazaridis et al 2014 | SA | Malawi_Chewa | 11 | Skoglund et al 2017 |

| | | | | | | | |
|---|---|---|---|---|---|---|---|
| AM | Aymara | 4 | Lazaridis et al 2014 | SA | Malawi_Ngoni | 4 | Skoglund et al 2017 |
| AM | Bolivian_LaPaz | 3 | Lazaridis et al 2014 | SA | Malawi_Tumbuka | 10 | Skoglund et al 2017 |
| AM | Bolivian_Pando | 3 | Lazaridis et al 2014 | SA | Malawi_Yao | 9 | Skoglund et al 2017 |
| AM | Cabecar | 6 | Lazaridis et al 2014 | SA | Nama | 16 | Lazaridis et al 2014 |
| AM | Chilote | 3 | Lazaridis et al 2014 | SA | Naro | 8 | Lazaridis et al 2014 |
| AM | Chipewyan | 19 | Lazaridis et al 2014 | SA | Shua | 9 | Lazaridis et al 2014 |
| AM | Cree | 11 | Lazaridis et al 2014 | SA | Taa_East | 6 | Lazaridis et al 2014 |
| AM | Guarani | 4 | Lazaridis et al 2014 | SA | Taa_North | 9 | Lazaridis et al 2014 |
| AM | Kaqchikel | 5 | Lazaridis et al 2014 | SA | Taa_West | 15 | Lazaridis et al 2014 |
| AM | Karitiana | 10 | Lazaridis et al 2014 | SA | Tshwa | 4 | Lazaridis et al 2014 |
| AM | Mayan | 17 | Lazaridis et al 2014 | SA | Tswana | 5 | Lazaridis et al 2014 |
| AM | Mixe | 10 | Lazaridis et al 2014 | SA | Wambo | 5 | Lazaridis et al 2014 |
| AM | Mixtec | 10 | Lazaridis et al 2014 | SA | Xuun | 13 | Lazaridis et al 2014 |
| AM | Ojibwa | 12 | Lazaridis et al 2014 | SA | Zulu | 100 | Gurdasani et al 2015 |
| AM | Piapoco | 3 | Lazaridis et al 2014 | SO | Somali | 13 | Lazaridis et al 2014 |
| AM | Pima | 11 | Lazaridis et al 2014 | SS | Balochi | 5 | Lazaridis et al 2014 |
| AM | Quechua_Coriell | 5 | Lazaridis et al 2014 | SS | Bengali_Bangladesh_BEB | 7 | Lazaridis et al 2014 |
| AM | Surui | 8 | Lazaridis et al 2014 | SS | Brahui | 20 | Lazaridis et al 2014 |
| AM | Zapotec | 10 | Lazaridis et al 2014 | SS | Burusho | 23 | Lazaridis et al 2014 |
| CA | BiakaPygmy | 20 | Lazaridis et al 2014 | SS | Cochin_Jew | 5 | Lazaridis et al 2014 |
| CA | Cameroon_Aghem | 28 | Lipson et al 2020 | SS | GujaratiA_GIH | 5 | Lazaridis et al 2014 |
| CA | Cameroon_Bafut | 15 | Lipson et al 2020 | SS | GujaratiB_GIH | 5 | Lazaridis et al 2014 |
| CA | Cameroon_Baka | 2 | Mallick et al 2016 | SS | GujaratiC_GIH | 5 | Lazaridis et al 2014 |
| CA | Cameroon_Bakoko | 1 | Lipson et al 2020 | SS | GujaratiD_GIH | 5 | Lazaridis et al 2014 |
| CA | Cameroon_Bakola | 2 | Mallick et al 2016 | SS | Hazara | 13 | Lazaridis et al 2014 |
| CA | Cameroon_Bangwa | 3 | Lipson et al 2020 | SS | India_Hindu | 12 | Lopez et al 2017 |
| CA | Cameroon_Bedzan | 2 | Mallick et al 2016 | SS | India_Zoroastrian | 13 | Lopez et al 2017 |
| CA | Cameroon_Fulani | 2 | Mallick et al 2016 | SS | Kalash | 16 | Lazaridis et al 2014 |
| CA | Cameroon_Kotoko | 7 | this study | SS | Kharia | 8 | Lazaridis et al 2014 |
| CA | Cameroon_Mada | 2 | Mallick et al 2016 | SS | Kusunda | 7 | Lazaridis et al 2014 |
| CA | Cameroon_Mbo | 28 | Lipson et al 2020 | SS | Lodhi | 13 | Lazaridis et al 2014 |
| CA | Cameroon_Ngumba | 2 | Mallick et al 2016 | SS | Makrani | 9 | Lazaridis et al 2014 |
| CA | Cameroon_Tikar_South | 2 | Mallick et al 2016 | SS | Mala | 13 | Lazaridis et al 2014 |
| CA | Esan_Nigeria_ESN | 8 | Lazaridis et al 2014 | SS | Onge | 6 | Lazaridis et al 2014 |
| CA | MbutiPygmy | 10 | Lazaridis et al 2014 | SS | Pathan | 19 | Lazaridis et al 2014 |
| CA | Yoruba | 70 | Lazaridis et al 2014 | SS | Punjabi_Lahore_PJL | 8 | Lazaridis et al 2014 |
| CS | Aleut | 5 | Lazaridis et al 2014 | SS | Sindhi | 18 | Lazaridis et al 2014 |
| CS | Altaian | 7 | Lazaridis et al 2014 | SS | Tiwari | 14 | Lazaridis et al 2014 |
| CS | Chukchi | 18 | Lazaridis et al 2014 | SS | Vishwabrahmin | 13 | Lazaridis et al 2014 |
| CS | Dolgan | 3 | Lazaridis et al 2014 | WA | CentralAfricanRepubli | 2 | Mallick et al 2016 |

| | | | | | | | |
|---|---|---|---|---|---|---|---|
| | | | | | c_Kaba | | |
| CS | Eskimo_Chaplin | 4 | Lazaridis et al 2014 | WA | Chad_Bulala | 2 | Mallick et al 2016 |
| CS | Eskimo_Naukan | 11 | Lazaridis et al 2014 | WA | Chad_Laka | 2 | Mallick et al 2016 |
| CS | Eskimo_Sireniki | 4 | Lazaridis et al 2014 | WA | Gambian_GWD | 6 | Lazaridis et al 2014 |
| CS | Even | 8 | Lazaridis et al 2014 | WA | Mandenka | 17 | Lazaridis et al 2014 |
| CS | Itelmen | 5 | Lazaridis et al 2014 | WA | Mende_Sierra_Leone_ MSL | 8 | Lazaridis et al 2014 |
| CS | Kalmyk | 10 | Lazaridis et al 2014 | WA | Senegal | 13 | this study |
| CS | Koryak | 9 | Lazaridis et al 2014 | WE | Abkhasian | 9 | Lazaridis et al 2014 |
| CS | Kyrgyz | 9 | Lazaridis et al 2014 | WE | Adygei | 16 | Lazaridis et al 2014 |
| CS | Mansi | 8 | Lazaridis et al 2014 | WE | Albanian | 6 | Lazaridis et al 2014 |
| CS | Nganasan | 10 | Lazaridis et al 2014 | WE | Armenian | 9 | Lazaridis et al 2014 |
| CS | Russian | 22 | Lazaridis et al 2014 | WE | Ashkenazi_Jew | 7 | Lazaridis et al 2014 |
| CS | Selkup | 10 | Lazaridis et al 2014 | WE | Balkar | 10 | Lazaridis et al 2014 |
| CS | Tajik_Pomiri | 8 | Lazaridis et al 2014 | WE | Basque_French | 20 | Lazaridis et al 2014 |
| CS | Tlingit | 4 | Lazaridis et al 2014 | WE | Basque_Spanish | 9 | Lazaridis et al 2014 |
| CS | Tubalar | 18 | Lazaridis et al 2014 | WE | BedouinA | 25 | Lazaridis et al 2014 |
| CS | Turkmen | 7 | Lazaridis et al 2014 | WE | BedouinB | 19 | Lazaridis et al 2014 |
| CS | Tuvinian | 9 | Lazaridis et al 2014 | WE | Belarusian | 10 | Lazaridis et al 2014 |
| CS | Ulchi | 23 | Lazaridis et al 2014 | WE | Bulgarian | 9 | Lazaridis et al 2014 |
| CS | Yakut | 20 | Lazaridis et al 2014 | WE | Chechen | 9 | Lazaridis et al 2014 |
| CS | Yukagir_Forest | 4 | Lazaridis et al 2014 | WE | Chuvash | 10 | Lazaridis et al 2014 |
| CS | Yukagir_Tundra | 13 | Lazaridis et al 2014 | WE | Croatian | 10 | Lazaridis et al 2014 |
| EA | BantuKenya | 6 | Lazaridis et al 2014 | WE | Cypriot | 8 | Lazaridis et al 2014 |
| EA | Buganda | 96 | Gurdasani et al 2015 | WE | Czech | 10 | Lazaridis et al 2014 |
| EA | Datog | 3 | Lazaridis et al 2014 | WE | Druze | 35 | Lazaridis et al 2014 |
| EA | Dinka | 7 | Lazaridis et al 2014 | WE | English_Cornwall_GB R | 5 | Lazaridis et al 2014 |
| EA | Hadza | 14 | Lazaridis et al 2014 | WE | English_Kent_GBR | 5 | Lazaridis et al 2014 |
| EA | Hadza_Henn | 3 | Lazaridis et al 2014 | WE | Estonian | 10 | Lazaridis et al 2014 |
| EA | Kenya_Elmolo | 2 | Mallick et al 2016 | WE | Finnish_FIN | 7 | Lazaridis et al 2014 |
| EA | Kenya_Kikuyu | 2 | Mallick et al 2016 | WE | French | 25 | Lazaridis et al 2014 |
| EA | Kenya_Ogiek | 2 | Mallick et al 2016 | WE | French_South | 7 | Lazaridis et al 2014 |
| EA | Kenya_Rendille | 2 | Mallick et al 2016 | WE | Georgian_Jew | 7 | Lazaridis et al 2014 |
| EA | Kenya_Sengwer | 2 | Mallick et al 2016 | WE | Georgian_Megrels | 10 | Lazaridis et al 2014 |
| EA | Kikuyu | 4 | Lazaridis et al 2014 | WE | Greek_Comas | 14 | Lazaridis et al 2014 |
| EA | Luhya_Kenya_LWK | 8 | Lazaridis et al 2014 | WE | Greek_Coriell | 6 | Lazaridis et al 2014 |
| EA | Luo | 8 | Lazaridis et al 2014 | WE | Hungarian_Coriell | 10 | Lazaridis et al 2014 |
| EA | Masai_Ayodo | 2 | Lazaridis et al 2014 | WE | Hungarian_Metspalu | 10 | Lazaridis et al 2014 |
| EA | Masai_Kinyawa_MKK | 9 | Lazaridis et al 2014 | WE | Icelandic | 12 | Lazaridis et al 2014 |
| EA | Sandawe | 22 | Lazaridis et al 2014 | WE | Iran_Fars | 17 | Lopez et al 2017 |

| | | | | | | | |
|---|---|---|---|---|---|---|---|
| EA | Tanzania_Hadza | 2 | Mallick et al 2016 | WE | Iranian | 8 | Lazaridis et al 2014 |
| EA | Tanzania_Iraqw | 2 | Mallick et al 2016 | WE | Iranian_Jew | 9 | Lazaridis et al 2014 |
| EA | Tanzania_Sandawe | 1 | Mallick et al 2016 | WE | Iran_Zoroastrian | 27 | Lopez et al 2017 |
| EA | Uganda_Muganda | 6 | this study | WE | Iraqi_Jew | 6 | Lazaridis et al 2014 |
| EA | Uganda_Mussese | 6 | this study | WE | Israeli_Arabs | 23 | this study |
| ES | Ami_Coriell | 10 | Lazaridis et al 2014 | WE | Italian_Bergamo | 12 | Lazaridis et al 2014 |
| ES | Atayal_Coriell | 9 | Lazaridis et al 2014 | WE | Italian_EastSicilian | 5 | Lazaridis et al 2014 |
| ES | Cambodian | 8 | Lazaridis et al 2014 | WE | Italian_Tuscan | 8 | Lazaridis et al 2014 |
| ES | Dai | 10 | Lazaridis et al 2014 | WE | Italian_WestSicilian | 6 | Lazaridis et al 2014 |
| ES | Daur | 9 | Lazaridis et al 2014 | WE | Jordanian | 4 | Lazaridis et al 2014 |
| ES | Han | 33 | Lazaridis et al 2014 | WE | Kumyk | 8 | Lazaridis et al 2014 |
| ES | Han_NChina | 10 | Lazaridis et al 2014 | WE | Lebanese | 8 | Lazaridis et al 2014 |
| ES | Hezhen | 7 | Lazaridis et al 2014 | WE | Lezgin | 9 | Lazaridis et al 2014 |
| ES | Japanese | 29 | Lazaridis et al 2014 | WE | Lithuanian | 10 | Lazaridis et al 2014 |
| ES | Kinh_Vietnam_KHV | 8 | Lazaridis et al 2014 | WE | Maltese | 8 | Lazaridis et al 2014 |
| ES | Korean | 6 | Lazaridis et al 2014 | WE | Mordovian | 10 | Lazaridis et al 2014 |
| ES | Lahu | 7 | Lazaridis et al 2014 | WE | Nogai | 9 | Lazaridis et al 2014 |
| ES | Miao | 10 | Lazaridis et al 2014 | WE | North_Ossetian | 10 | Lazaridis et al 2014 |
| ES | Mongola | 6 | Lazaridis et al 2014 | WE | Norwegian | 11 | Lazaridis et al 2014 |
| ES | Naxi | 8 | Lazaridis et al 2014 | WE | Orcadian | 12 | Lazaridis et al 2014 |
| ES | Oroqen | 9 | Lazaridis et al 2014 | WE | Palestinian | 33 | Lazaridis et al 2014 |
| ES | She | 9 | Lazaridis et al 2014 | WE | PalestinianArabs | 13 | this study |
| ES | Thai | 8 | Lazaridis et al 2014 | WE | Sardinian | 27 | Lazaridis et al 2014 |
| ES | Tu | 10 | Lazaridis et al 2014 | WE | Saudi | 8 | Lazaridis et al 2014 |
| ES | Tujia | 10 | Lazaridis et al 2014 | WE | SaudiBeduins | 8 | this study |
| ES | Uygur | 10 | Lazaridis et al 2014 | WE | Scottish_Argyll_Bute_GBR | 4 | Lazaridis et al 2014 |
| ES | Xibo | 7 | Lazaridis et al 2014 | WE | Spanish_Andalucia_IBS | 4 | Lazaridis et al 2014 |
| ES | Yi | 10 | Lazaridis et al 2014 | WE | Spanish_Aragon_IBS | 6 | Lazaridis et al 2014 |
| EG | Egyptian_Comas | 11 | Lazaridis et al 2014 | WE | Spanish_Baleares_IBS | 4 | Lazaridis et al 2014 |
| EG | Egyptian_Metspalu | 7 | Lazaridis et al 2014 | WE | Spanish_Cantabria_IBS | 5 | Lazaridis et al 2014 |
| NA | Algerian | 4 | Lazaridis et al 2014 | WE | Spanish_Castilla_la_Mancha_IBS | 5 | Lazaridis et al 2014 |
| NA | Libyan_Jew | 9 | Lazaridis et al 2014 | WE | Spanish_Castilla_y_Leon_IBS | 5 | Lazaridis et al 2014 |
| NA | Moroccan_Jew | 6 | Lazaridis et al 2014 | WE | Spanish_Cataluna_IBS | 5 | Lazaridis et al 2014 |
| NA | MoroccoBerber | 18 | this study | WE | Spanish_Extremadura_IBS | 5 | Lazaridis et al 2014 |
| NA | Mozabite | 21 | Lazaridis et al 2014 | WE | Spanish_Galicia_IBS | 5 | Lazaridis et al 2014 |
| NA | Saharawi | 6 | Lazaridis et al 2014 | WE | Spanish_Murcia_IBS | 4 | Lazaridis et al 2014 |
| NA | Tunisian | 8 | Lazaridis et al 2014 | WE | Spanish_Pais_Vasco_IBS | 5 | Lazaridis et al 2014 |

| | | | | | | | |
|---|---|---|---|---|---|---|---|
| NA | Tunisian_Jew | 7 | Lazaridis et al 2014 | WE | Spanish_Valencia_IBS | 5 | Lazaridis et al 2014 |
| OC | Australian_ECCAC | 2 | Lazaridis et al 2014 | WE | Syria | 12 | this study |
| OC | Bougainville | 10 | Lazaridis et al 2014 | WE | Syrian | 2 | Lazaridis et al 2014 |
| OC | Papuan | 13 | Lazaridis et al 2014 | WE | Turkish | 4 | Lazaridis et al 2014 |
| SA | Bantu_SA | 8 | Lazaridis et al 2014 | WE | Turkish_Adana | 10 | Lazaridis et al 2014 |
| SA | Damara | 12 | Lazaridis et al 2014 | WE | Turkish_Aydin | 7 | Lazaridis et al 2014 |
| SA | Gana | 7 | Lazaridis et al 2014 | WE | Turkish_Balikesir | 6 | Lazaridis et al 2014 |
| SA | Gui | 7 | Lazaridis et al 2014 | WE | Turkish_Istanbul | 10 | Lazaridis et al 2014 |
| SA | Haiom | 7 | Lazaridis et al 2014 | WE | Turkish_Jew | 8 | Lazaridis et al 2014 |
| SA | Himba | 4 | Lazaridis et al 2014 | WE | Turkish_Kayseri | 10 | Lazaridis et al 2014 |
| SA | Hoan | 6 | Lazaridis et al 2014 | WE | Turkish_Trabzon | 9 | Lazaridis et al 2014 |
| SA | Ju_hoan_North | 21 | Lazaridis et al 2014 | WE | Ukrainian_East | 6 | Lazaridis et al 2014 |
| SA | Ju_hoan_South | 5 | Lazaridis et al 2014 | WE | Ukrainian_West | 3 | Lazaridis et al 2014 |
| SA | Kgalagadi | 5 | Lazaridis et al 2014 | WE | Uzbek | 10 | Lazaridis et al 2014 |
| SA | Khomani | 9 | Lazaridis et al 2014 | WE | Yemen | 6 | Lazaridis et al 2014 |
| SA | Khwe | 8 | Lazaridis et al 2014 | WE | Yemenite_Jew | 8 | Lazaridis et al 2014 |

(b) ancient individuals:

| **Population** | **Country** | **Samples** | **Reference** |
|---|---|---|---|
| Anatolia_Neolithic | Turkey | Bar8, I0707, I0708, I0709, I0745, I0746, I1579, I1583, I1585 | Hofmanova et al, 2016; Mathieson et al, 2015 |
| South_Africa_Stone_Age | South Africa | BallitoBayA | Schlebusch et al 2017 |
| Iberia_Chalcolithic | Spain | I0581 | Mathieson et al, 2015 |
| Iceman_Neolithic | Italy | Iceman | Keller et al, 2012 |
| Karasuk | Russia | RISE493, RISE497 | Allentoft et al, 2015 |
| LaBrana (Iberian_Hunter_gatherer) | Iberia | LaBrana | Olalde et al, 2014 |
| Loschbour_Hunter_gatherer | Luxembourg | Loschbour | Lazaridis et al, 2014 |
| ~4.5kya East_Africa_forager (Ethiopia) | Ethiopia | Mota/GB20 | Gallego-Llorente et al., 2015 |
| West_Africa_Stone_to_Metal_Age | Cameroon | I10871_8 | Lipson et al, 2020 |
| Srubnaya | Russia | I0232 | Mathieson et al 2015 |
| East_Africa_Iron_Age | Kenya | I8802 | Prendergast et al 2019 |
| South_Africa_Iron_Age | South Africa | ElandCave, Mfongosi, Newcastle | Schlebusch et al 2017 |

| | | | |
|---|---|---|---|
| East_Africa_Later_Stone_Age | Kenya | I8808 | Prendergast et al 2019 |
| East_Africa_Pastoral_Iron_Age | Kenya | I12379 | Prendergast et al 2019 |
| East_Africa_Pastoral_Neolithic | Kenya / Tanzania | I10719, I13762, I13979, I13980, I8805, I8809, I8814, I8874, I8918, I8919, I8920, I8922 | Prendergast et al 2019 |
| Iranian_Neolithic | Iran | WC1 | Broushaki et al, 2016 |
| Yamnaya | Russia | I0231, Yamnaya | Haak et al, 2015 |

**Table S3 Mean and inner 95% empirical quantiles of genetic similarity (1-TVD) under the "Ethiopia-internal" analysis** between all pairs of individuals, or restricting to individuals that are from the same self-reported group label, whose group labels belong to the same major language group, or who speak the same first language, second language or have the same religious affiliation. Results are also shown after first conditioning on geographic distance ("Geo") or elevation difference ("Elev"), and rescaling each to span the same range as the first column.

| | **1-TVD** | **1-TVD \| Geo** | **1-TVD \| Elev** |
|---|---|---|---|
| **all** | 0.587 (0.35-0.888) | 0.559 (0.3-0.873) | 0.467 (0.208-0.779) |
| **same group** | 0.868 (0.637-0.942) | 0.728 (0.446-0.933) | 0.734 (0.483-0.846) |
| **same major language** | 0.654 (0.413-0.912) | 0.614 (0.356-0.886) | 0.531 (0.265-0.797) |
| **same 1st language** | 0.766 (0.46-0.935) | 0.683 (0.306-0.934) | 0.635 (0.301-0.834) |
| **same 2nd language** | 0.624 (0.386-0.893) | 0.594 (0.339-0.863) | 0.501 (0.243-0.783) |
| **same religion** | 0.606 (0.347-0.889) | 0.573 (0.296-0.863) | 0.482 (0.199-0.779) |

**Table S4 Parameters defining best-fit lines when testing for associations between genetic similarity and various factors (rows) while accounting for geographic distance, under the "Ethiopia-internal" analysis (first 3 columns, corresponding to Fig 1b) and the "Ethiopia-external" analysis (last 2 columns, corresponding to Fig S6b).**

| | rate | geo dist = ∞ | geo dist = 0 | | geo dist = 0 | slope (TVD / 100 km) |
|---|---|---|---|---|---|---|
| **Geo distance** | 0.013 | 0.562 | 0.779 | | 0.915 | -0.013 |
| **Group label** | 655.45 | 0.862 | 0.878 | | 0.954 | -0.004 |
| **Lang group** | 0.011 | 0.61 | 0.797 | | 0.938 | -0.011 |
| **1st lang** | 0.283 | 0.746 | 0.86 | | 0.954 | -0.009 |
| **2nd lang** | 0.01 | 0.585 | 0.803 | | 0.917 | -0.012 |
| **religion** | 0.012 | 0.571 | 0.782 | | 0.92 | -0.014 |

**Table S5 Parameters defining best-fit lines when testing for associations between genetic similarity and various factors (rows) while accounting for elevation difference, under the "Ethiopia-internal" analysis (first 2 columns, corresponding to Fig S6a) and the "Ethiopia-external" analysis (last 2 columns, corresponding to Fig S6c).**

| | geo dist = 0 | slope (TVD / km) | | geo dist = 0 | slope (TVD / km) |
|---|---|---|---|---|---|
| **Elev difference** | 0.638 | -0.067 | | 0.916 | -0.059 |
| **Group label** | 0.872 | -0.016 | | 0.953 | -0.006 |
| **Lang group** | 0.683 | -0.046 | | 0.929 | -0.032 |
| **1st lang** | 0.792 | -0.064 | | 0.943 | -0.024 |
| **2nd lang** | 0.668 | -0.065 | | 0.919 | -0.052 |
| **religion** | 0.649 | -0.064 | | 0.923 | -0.059 |

**Table S6 Associations of genetic similarity with various factors, while accounting for (a) geographic distance under the "Ethiopia-internal" analysis, (b) elevation difference under the "Ethiopia-internal" analysis, (c) geographic distance under the "Ethiopia-external" analysis and (d) elevation difference under the "Ethiopia-external" analysis.** In the second row ("Geo distance/Elevation") and second column ("All"), values give the proportions of 1000 permutations that were more associated with genetic similarity than the un-permuted data, when testing for an association with (a,c) geographic distance or (b,d) elevation difference (see Methods). Column 2 ("All") in the subsequent rows give analogous proportions when testing an additional factor (1st column) for association with genetic similarity after accounting for spatial distance: sharing a common group label ("Group label"), having ethnicities from the same language branch ("Lang group": AA Cushitic, AA Omotic, AA Semitic, NS Satellite-Core), sharing the same first language

("1st lang"), second language ("2nd lang") or religious affiliation ("religion"). Columns 3-7 depict results when permuting in a manner to account for each other factor. Fig 1b and Fig S6 give the maximum across values in each row (*=ignored when determining this final p-value; see Methods). For the first row in Table S6a-d, the p-value to the right of the "/" is after adjusting geographic distance and elevation for each other (see Methods).

**Table S6a: Genetic similarity versus geographic distance, "Ethiopia-internal" analysis**

| | **All** | **group label** | **lang group** | **1st lang** | **2nd lang** | **religion** |
|---|---|---|---|---|---|---|
| **Geo distance** | 0 / 0 | 0 / 0* | 0 / 0 | 0 / 0* | 0 / 0 | 0 / 0 |
| **Group label** | 0 | NA | 0 | 0* | 0 | 0 |
| **Lang group** | 0 | NA | NA | 0.155* | 0 | 0 |
| **1st lang** | 0 | 0* | 0.006 | NA | 0 | 0 |
| **2nd lang** | 0 | 1 | 0 | 0.992 | NA | 0 |
| **religion** | 0.998 | 1 | 0.445 | 1 | 0.982 | NA |

**Table S6b: Genetic similarity versus elevation difference, "Ethiopia-internal" analysis**

| | **All** | **group label** | **lang group** | **1st lang** | **2nd lang** | **religion** |
|---|---|---|---|---|---|---|
| **Elevation** | 0 / 0 | 0.015 / 0.337* | 0 / 0 | 0.066 / 1* | 0 / 0 | 0 / 0 |
| **Group label** | 0 | NA | 0 | 0* | 0 | 0 |
| **Lang group** | 0 | NA | NA | 0.003* | 0 | 0 |
| **1st lang** | 0 | 0.078* | 0 | NA | 0 | 0 |
| **2nd lang** | 0.422 | 1 | 0.667 | 1 | NA | 0.958 |
| **religion** | 0.992 | 1 | 1 | 1 | 0.895 | NA |

**Table S6c: Genetic similarity versus geographic distance, "Ethiopia-external" analysis**

| | **All** | **group label** | **lang group** | **1st lang** | **2nd lang** | **religion** |
|---|---|---|---|---|---|---|
| **Geo distance** | 0 / 0 | 0 / 0* | 0 / 0 | 0 / 0* | 0 / 0 | 0 / 0 |
| **Group label** | 0 | NA | 0 | 0.922* | 0 | 0 |
| **Lang group** | 0 | NA | NA | 0.36* | 0 | 0 |
| **1st lang** | 0 | 0.172* | 0 | NA | 0 | 0 |
| **2nd lang** | 0.852 | 1 | 0.999 | 1 | NA | 0.979 |
| **religion** | 1 | 0.991 | 1 | 0.998 | 0.999 | NA |

**Table S6d: Genetic similarity versus elevation difference, "Ethiopia-external" analysis**

| | **All** | **group label** | **lang group** | **1st lang** | **2nd lang** | **religion** |
|---|---|---|---|---|---|---|
| **Elevation** | 0 / 0 | 0 / 0* | 0 / 0 | 0 / 0* | 0 / 0 | 0 / 0 |
| **Group label** | 0 | NA | 0 | 0* | 0 | 0 |
| **Lang group** | 0 | NA | NA | 0.001* | 0 | 0 |
| **1st lang** | 0 | 0.809* | 0 | NA | 0 | 0 |
| **2nd lang** | 0.997 | 1 | 1 | 1 | NA | 0.996 |
| **religion** | 0.712 | 0.985 | 1 | 0.506 | 0.548 | NA |

**Table S7 Median and interquartile range (IQR) of spatial distance between individuals.** These values, are shown for all pairwise combinations of individuals (“All”), or as the median/IQR across median distances of all pairwise combinations of individuals within each group label (“group label”), major language group (“lang group”), first language (“1st lang”), second language (“2nd lang”) or religious affiliation (“religion”).

| | **All** | **group label** | **lang group** | **1st lang** | **2nd lang** | **religion** |
|---|---|---|---|---|---|---|
| **Geo distance (km)** | 252 (143-514) | 30 (0-46) | 223 (210-243) | 42 (0-50) | 59 (0-99) | 227 (103-340) |
| **Elevation (m)** | 643 (290-1134) | 180 (0-283) | 463 (354-536) | 228 (0-364) | 303 (0-561) | 368 (265-552) |

**Table S8 Genetic similarity among individuals across religious affiliations.** Average genetic similarity under the “Ethiopia-internal” analysis between two individuals with religious affiliation = Christian (“C”), Muslim (“M”) or Traditional (“T”), within each of 16 Ethiopian groups with n >= 5 sampled individuals from each of at least two of these religious affiliations. Also given is the average similarity between two individuals from separate religions (“C vs M”, “C vs T”) within each group. Asterisks denote that, within the ethnicity, the average genetically similarity is significantly higher between two individuals from that same religion versus individuals from different religions at p-value <0.05(*) or <0.001(**), based on 1000 permutations of the two compared religious affiliations within each group. Under the “Ethiopia-external” analysis, Christians within each of Tigraway and Murle, and Traditional within each of Bana and Chara, have p-value < 0.05.

| **group** | **C (n)** | **M (n)** | **T (n)** | **C vs M** | **C vs T** |
|---|---|---|---|---|---|
| **Alae** | 0.832 (32) | **0.9** (10)** | 0.906 (4) | 0.835 | -- |
| **Amhara** | 0.913 (11) | 0.923 (17) | -- (0) | 0.924 | -- |
| **Ari Potter** | 0.805 (12) | -- (0) | 0.881 (12) | -- | 0.824 |
| **Bana** | 0.75 (14) | -- (0) | 0.824 (14) | -- | 0.763 |
| **Bodi** | 0.873 (9) | -- (0) | 0.883 (5) | -- | 0.88 |

| | | | | | |
|---|---|---|---|---|---|
| **Chara** | 0.83 (9) | -- (0) | **0.916* (8)** | -- | 0.852 |
| **Dasanech** | 0.822 (5) | -- (0) | 0.874 (10) | -- | 0.849 |
| **Gedeo** | 0.903 (10) | 0.932 (11) | -- (0) | 0.929 | -- |
| **Gurage** | 0.89 (11) | 0.908 (5) | -- (0) | 0.895 | -- |
| **Hamar** | 0.807 (6) | -- (0) | 0.843 (8) | -- | 0.822 |
| **Honsita** | 0.846 (12) | **0.926* (5)** | -- (0) | 0.897 | -- |
| **Manja** | 0.935 (5) | -- (0) | 0.906 (9) | -- | 0.921 |
| **Murle** | **0.895* (7)** | -- (0) | 0.844 (6) | -- | 0.85 |
| **Sidama** | 0.872 (11) | **0.925* (10)** | -- (0) | 0.895 | -- |
| **Tigraway** | **0.921* (8)** | 0.907 (5) | -- (0) | 0.908 | -- |
| **Tsamay** | 0.78 (8) | -- (1) | 0.757 (9) | 0.688 | 0.753 |

**Table S9 Genetic association with cultural similarity.** P-values from Mantel tests for association between genetic and cultural similarity among ethnicities ("All"), and from partial Mantel tests that accounts for one of geographic distance, elevation, or language branch (AA Cushitic, AA Omotic, AA Semitic, NS Satellite-Core) when testing for an association between genetic and cultural similarity. Within each analysis ("Ethiopia-internal", "Ethiopia-external"), the first row measures cultural similarity as the number of matching reported cultural practices across ethnicities, while the second row up-weights sharing of rare cultural practices among ethnicities (see Methods).

| **Analysis** | | **All** | **Geographic distance** | **Elevation** | **Language** |
|---|---|---|---|---|---|
| **Eth-internal** | equal practice weight | 0.0227 | 0.0152 | 0.0448 | 0.0279 |
| | up-weight rare practices | 0.023 | 0.036 | 0.061 | 0.061 |
| **Eth-external** | equal practice weight | 0.214 | 0.173 | 0.440 | 0.280 |
| | up-weight rare practices | 0.114 | 0.181 | 0.300 | 0.233 |

**Table S10 Association of ancestry sharing with Mota to spatial distance.** Effect sizes, standard errors and p-values for a linear regression of SOURCEFIND-inferred ancestry matching to the 4.5kya Ethiopian Mota versus geographic/elevation distance of modern individuals from Mota (Fig 4cd).

| analysis | Effect size (% per km) | SE (% per km) | p-value |
|---|---|---|---|
| **geographic distance** | -4.64e-02 | 9.68e-03 | 0.000008 |
| **elevation difference** | -13.7 | 3.71 | 0.000418 |

**Table S11 Evidence of recent intermixing among groups reporting particular cultural practices** under the "Ethiopia-internal" analysis. Out of 46 SNNPR groups for which we analysed reported cultural traits, the number of groups sharing each trait (*n*), and the number of pairings of these *n* groups (all other columns) whereby individuals from separate groups share average inferred MRCA segments >*X*cM longer than the median segment lengths across all pairings of individuals from the separate groups, indicative of recent intermixing. Values in italics give the proportion of times, out of 10K re-samples, where a greater or equal number of group pairings among *n* randomly selected groups (out of the 46 SNNPR groups) showed this trend.

| practice | n | >2cM | >2.5 | >3 | >3.5 | >4 | >4.5 | >5cM |
|---|---|---|---|---|---|---|---|---|
| **Female circumcision** | 3 | 2 | 2 | 1 | 1 | 1 | 1 | 1 |
| | | *0.0757* | *0.02186* | *0.12684* | *0.08071* | *0.05573* | *0.03872* | *0.03038* |
| **Male circumcision** | 11 | 19 | 10 | 7 | 6 | 3 | 3 | 3 |
| | | *0.01096* | *0.03232* | *0.02587* | *0.01085* | *0.09501* | *0.04903* | *0.02985* |
| **Sororate/cousin marriages** | 7 | 10 | 5 | 3 | 2 | 2 | 1 | 1 |
| | | *0.00696* | *0.04072* | *0.07942* | *0.119* | *0.06363* | *0.24267* | *0.19449* |

**Table S12** For each group *X*={*Mursi, Suri, Zilmamo*} (columns), all of which traditionally wear decorative lip plates, each row gives average genetic similarity (based on TVD) between individuals from *X* versus those from other listed group **(a)** under the **"Ethiopia-internal"** analysis, **(b)** relative to that expected under geographic distance (fit as described in Methods) under the **"Ethiopia-internal"** analysis, **(c)** under the **"Ethiopia-external"** analysis, and **(d)** relative to that expected under geographic distance under the **"Ethiopia-external"** analysis. I.e. values reflect averages across all pairwise comparisons of individuals, with one individual from *X* and the other from the group in the row. For each of *X*={*Mursi*, *Suri*, *Zilmamo*} and the average across these three, the six groups with six highest such values are shown. Letters in parentheses give the language branch classification of that ethnicity (O = Afroasiatic Omotic, N = Nilo-Saharan Satellite-Core).

**Table S12a: Genetic similarity with other groups, "Ethiopia-internal" analysis**

| **Mursi** | | **Suri** | | **Zilmamo** | | **Average** | |
|---|---|---|---|---|---|---|---|
| **Mursi (N)** | **0.9** | **Suri (N)** | **0.914** | **Suri (N)** | **0.89** | **Suri (N)** | **0.873** |
| **Suri (N)** | **0.814** | **Zilmamo (N)** | **0.89** | **Zilmamo (N)** | **0.881** | **Zilmamo (N)** | **0.856** |
| **Zilmamo (N)** | **0.796** | **Mursi (N)** | **0.814** | **Mursi (N)** | **0.796** | **Mursi (N)** | **0.837** |
| Bodi (N) | 0.754 | Bodi (N) | 0.735 | Bodi (N) | 0.725 | Bodi (N) | 0.738 |
| Karo (O) | 0.719 | Karo (O) | 0.728 | Karo (O) | 0.723 | Karo (O) | 0.724 |
| Murle (N) | 0.685 | Anuak (N) | 0.682 | Anuak (N) | 0.681 | Neur (N) | 0.682 |

**Table S12b: Genetic similarity with other groups, relative to geographic distance, "Ethiopia-internal" analysis**

| **Mursi** | | **Suri** | | **Zilmamo** | | **Average** | |
|---|---|---|---|---|---|---|---|
| **Suri (N)** | **0.195** | **Mursi (N)** | **0.195** | **Mursi (N)** | **0.177** | **Mursi (N)** | **0.168** |
| **Zilmamo (N)** | **0.177** | Karo (O) | 0.137 | Karo (O) | 0.133 | **Suri (N)** | **0.148** |
| **Mursi (N)** | **0.133** | **Suri (N)** | **0.137** | **Suri (N)** | **0.112** | **Zilmamo (N)** | **0.131** |
| Nuer (N) | 0.121 | Nuer (N) | 0.114 | Nuer (N) | 0.11 | Nuer (N) | 0.115 |
| Anuak (N) | 0.113 | **Zilmamo (N)** | **0.112** | **Zilmamo (N)** | 0.103 | Karo (O) | 0.11 |
| Murle (N) | 0.077 | Anuak (N) | 0.104 | Anuak (N) | 0.103 | Anuak (N) | 0.107 |

**Table S12c: Genetic similarity with other groups, “Ethiopia-external” analysis**

| **Mursi** | | **Suri** | | **Zilmamo** | | **Average** | |
|---|---|---|---|---|---|---|---|
| **Mursi (N)** | **0.967** | **Suri (N)** | **0.966** | **Suri (N)** | **0.958** | **Suri (N)** | **0.961** |
| Bodi (N) | 0.963 | Kwegu (N) | 0.96 | **Zilmamo (N)** | **0.956** | **Mursi (N)** | **0.959** |
| **Suri (N)** | **0.958** | Bodi (N) | 0.959 | Kwegu (N) | 0.955 | Bodi (N) | 0.958 |
| **Zilmamo (N)** | **0.951** | **Zilmamo (N)** | **0.958** | Bodi (N) | 0.952 | **Zilmamo (N)** | **0.955** |
| Kwegu (N) | 0.95 | **Mursi (N)** | **0.958** | **Mursi (N)** | **0.951** | Kwegu (N) | 0.955 |
| Karo (O) | 0.944 | Mezhenger (N) | 0.953 | Mezhenger (N) | 0.949 | Mezhenger (N) | 0.948 |

**Table S12d: Genetic similarity with other groups, relative to geographic distance, “Ethiopia-external” analysis**

| **Mursi** | | **Suri** | | **Zilmamo** | | **Average** | |
|---|---|---|---|---|---|---|---|
| Gumuz (N) | 0.074 | Gumuz (N) | 0.079 | Gumuz (N) | 0.078 | Gumuz (N) | 0.077 |
| **Suri (N)** | **0.056** | Kwegu (N) | 0.064 | Kwegu (N) | 0.058 | Komo (N) | 0.058 |
| Komo (N) | 0.055 | Komo (N) | 0.059 | Komo (N) | 0.058 | Kwegu (N) | 0.055 |
| Bodi (N) | 0.054 | **Mursi (N)** | **0.056** | Karo (O) | 0.051 | **Mursi (N)** | **0.053** |
| **Mursi (N)** | **0.053** | Karo (O) | 0.056 | **Mursi (N)** | **0.049** | Bodi (N) | 0.052 |
| Nuer (N) | 0.053 | Bodi (N) | 0.053 | Mezhenger (N) | 0.048 | **Suri (N)** | **0.05** |

**Supplementary Figures**

a)

b)

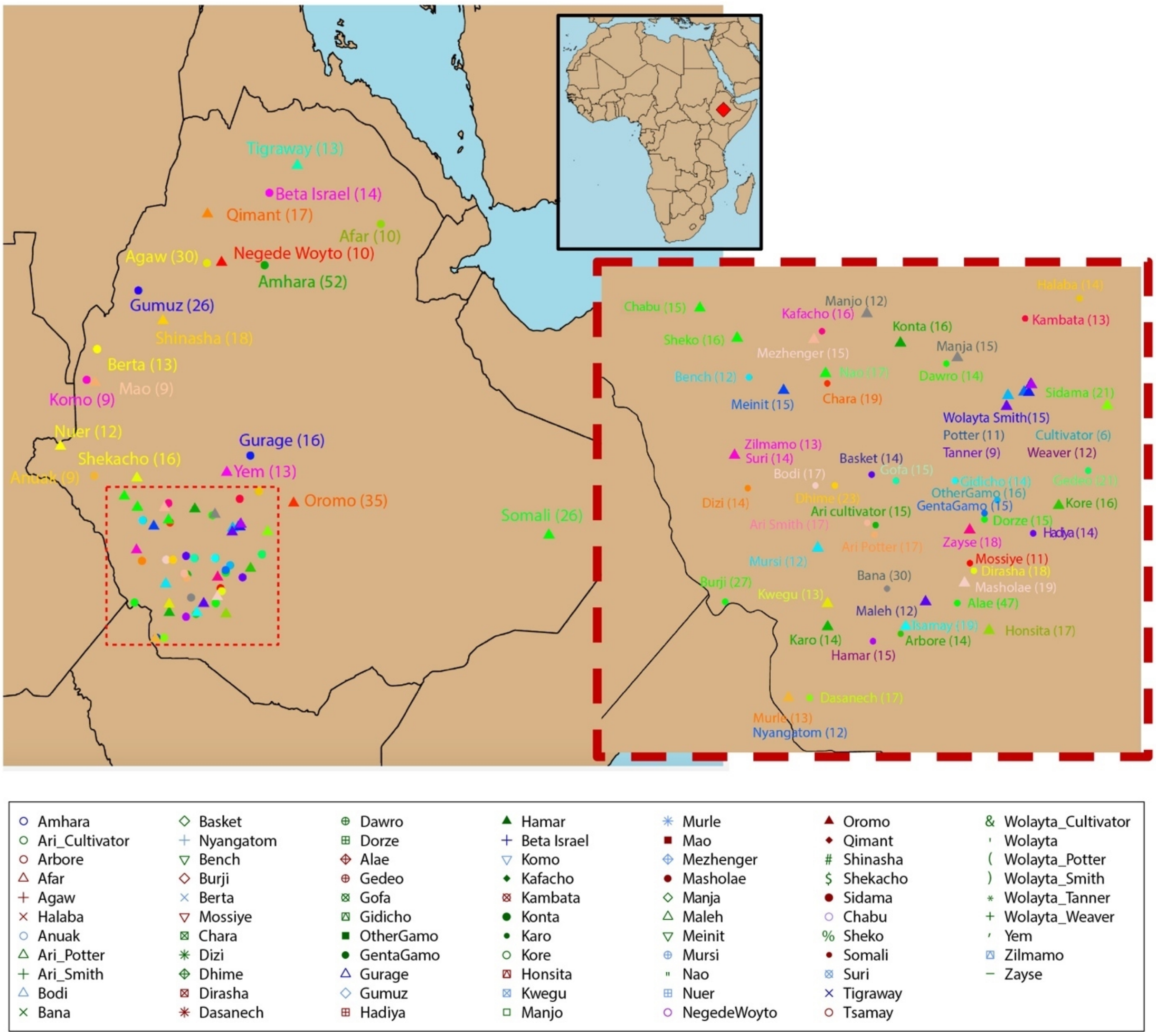

**Figure S1. (A) Location of all sampled Ethiopian groups included in this study.** Number of individuals from each self-reported group included in this work. Labels are placed at the average location (based on birthplace) of sampled individuals from that group. **(B)** Legend for Fig 1a of the main text.

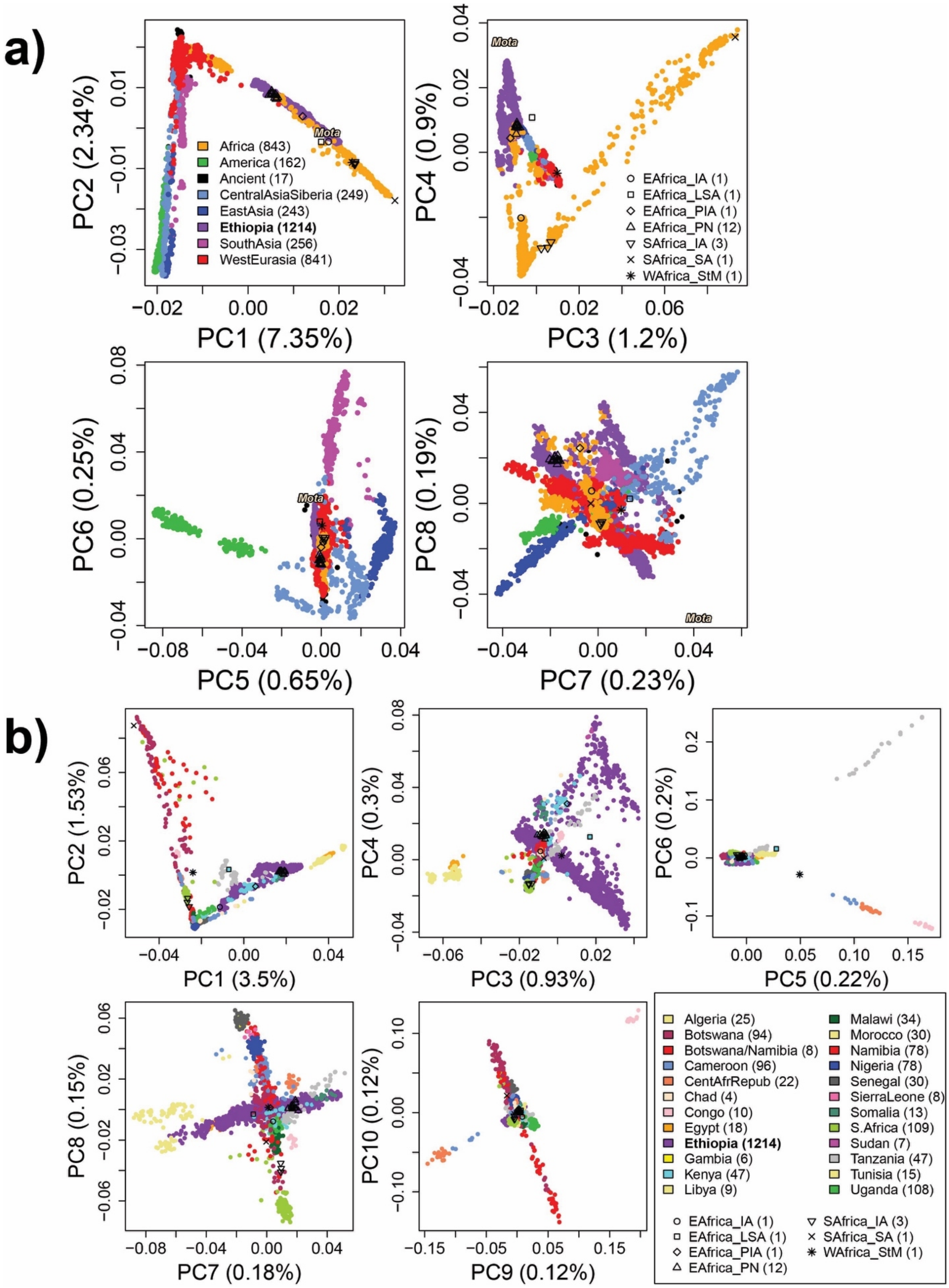
a)
PC1 (7.35%)
PC2 (2.34%)
Africa (843)
America (162)
Ancient (17)
CentralAsiaSiberia (249)
EastAsia (243)
Ethiopia (1214)
SouthAsia (256)
WestEurasia (841)
Mota
PC3 (1.2%)
PC4 (0.9%)
EAfrica_IA (1)
EAfrica_LSA (1)
EAfrica_PIA (1)
EAfrica_PN (12)
SAfrica_IA (3)
SAfrica_SA (1)
WAfrica_StM (1)
PC5 (0.65%)
PC6 (0.25%)
PC7 (0.23%)
PC8 (0.19%)
b)
PC1 (3.5%)
PC2 (1.53%)
PC3 (0.93%)
PC4 (0.3%)
PC5 (0.22%)
PC6 (0.2%)
PC7 (0.18%)
PC8 (0.15%)
PC9 (0.12%)
PC10 (0.12%)
Algeria (25)
Botswana (94)
Botswana/Namibia (8)
Cameroon (96)
CentAfrRepub (22)
Chad (4)
Congo (10)
Egypt (18)
Ethiopia (1214)
Gambia (6)
Kenya (47)
Libya (9)
Malawi (34)
Morocco (30)
Namibia (78)
Nigeria (78)
Senegal (30)
SierraLeone (8)
Somalia (13)
S.Africa (109)
Sudan (7)
Tanzania (47)
Tunisia (15)
Uganda (108)
EAfrica_IA (1)
EAfrica_LSA (1)
EAfrica_PIA (1)
EAfrica_PN (12)
SAfrica_IA (3)
SAfrica_SA (1)
WAfrica_StM (1)

**Figure S2. (A)** First eight principal components (PC) of a principal-components-analysis (PCA) of all the samples included in this study. Numbers from each label/region are given in parenthesis (note the unequal sample sizes). “Ancient” refers to non-African aDNA samples, other black symbols are aDNA samples from Kenya/Tanzania (“EAfrica”, Prendergast et al 2019), South Africa (“SAfrica”, Schlebusch et al 2017), and Cameroon (“WAfrica”, Lipson et al 2020), with IA = Iron Age, LSA = Later Stone Age, PIA = Pastoral Iron Age, PN = Pastoral Neolithic, SA = Stone Age, StM = Stone-to-Metal Age (Table S2). **(B)** PCA of all the African samples included in this study.

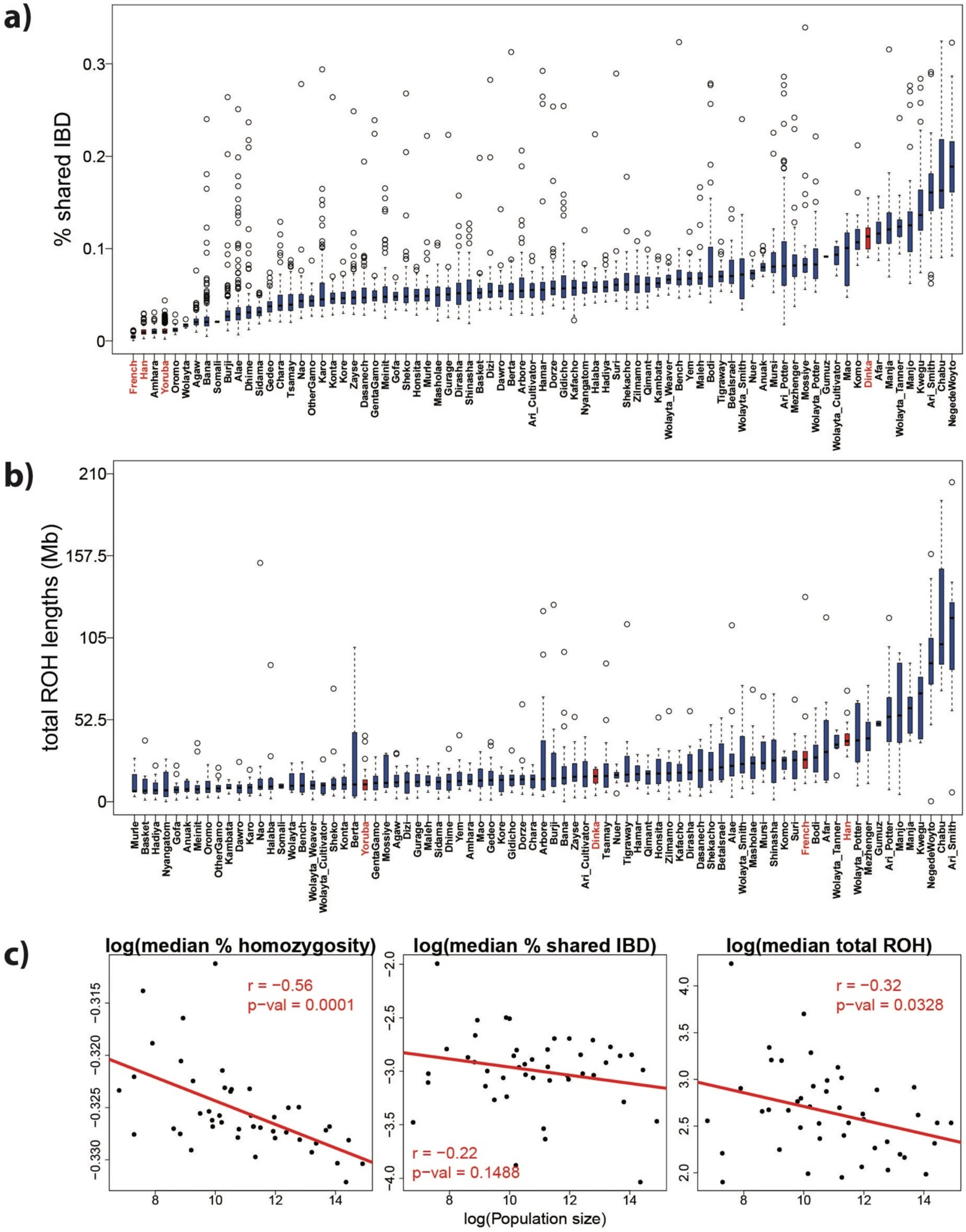
a)
% shared IBD
b)
total ROH lengths (Mb)
c)
log(median % homozygosity)
r = −0.56
p-val = 0.0001
log(median % shared IBD)
r = −0.22
p-val = 0.1488
log(median total ROH)
r = −0.32
p-val = 0.0328
log(Population size)

**Figure S3 Genetic homogeneity estimates for the Ethiopian groups, and correlation with population census size**. **(a)** Distribution of the proportion of genome shared identical-by-descent (IBD) across all pairs of (non-excluded) individuals within each Ethiopian group (blue) and other populations (red). **(b)** Distribution of the total runs of homozygosity (ROH) within (non-excluded) individuals from each Ethiopian group (blue) and other populations (red). **(c)** Median homozygosity, IBD and ROH values (on log-scale) versus log population census size, with the red line giving the best simple linear regression fit, and correlation (r) and the p-value of this fit provided. In general, we note an increase in genetic diversity (i.e. reduced homogeneity) of ethnic groups with larger census population sizes, though this is not always significant. As expected, the highest levels of homozygosity were detected in populations with low census sizes, likely reflecting an elevated degree of endogamy and marriage between close kin.

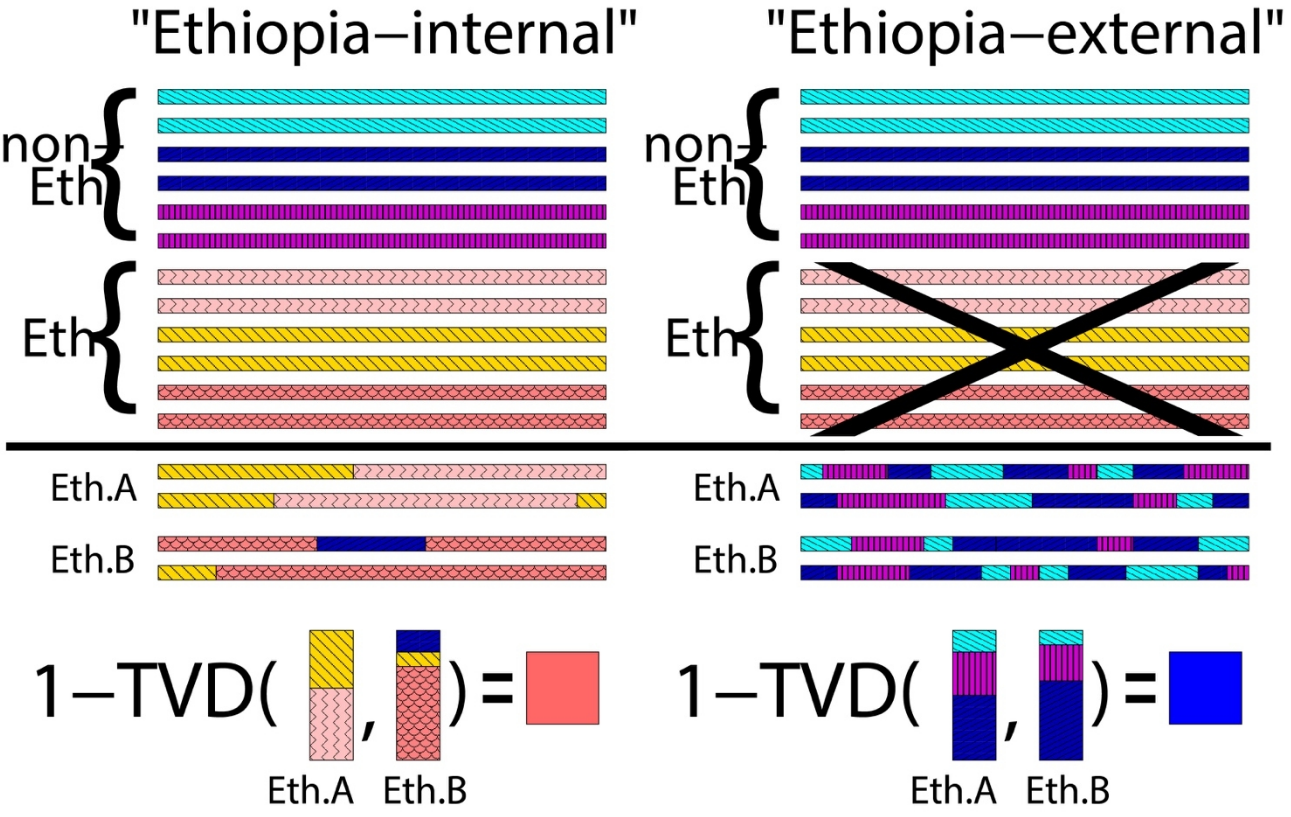


**Figure S4. Schematic of two CHROMOPAINTER analyses.** In the "Ethiopa-internal" analysis, haplotype patterns in Ethiopians *A* and *B* ("Eth.A","Eth.B") are matched to those of non-Ethiopians and other Ethiopians, giving the genome-wide paintings below the black line, which are then compared to give the genetic similarity (1-TVD) between *A* and *B* (red square). In the "Ethiopia-external" analysis, *A* and *B* are matched only to non-Ethiopians, typically giving a higher genetic similarity between their respective paintings (blue square) by substantially mitigating effects of recent isolation.

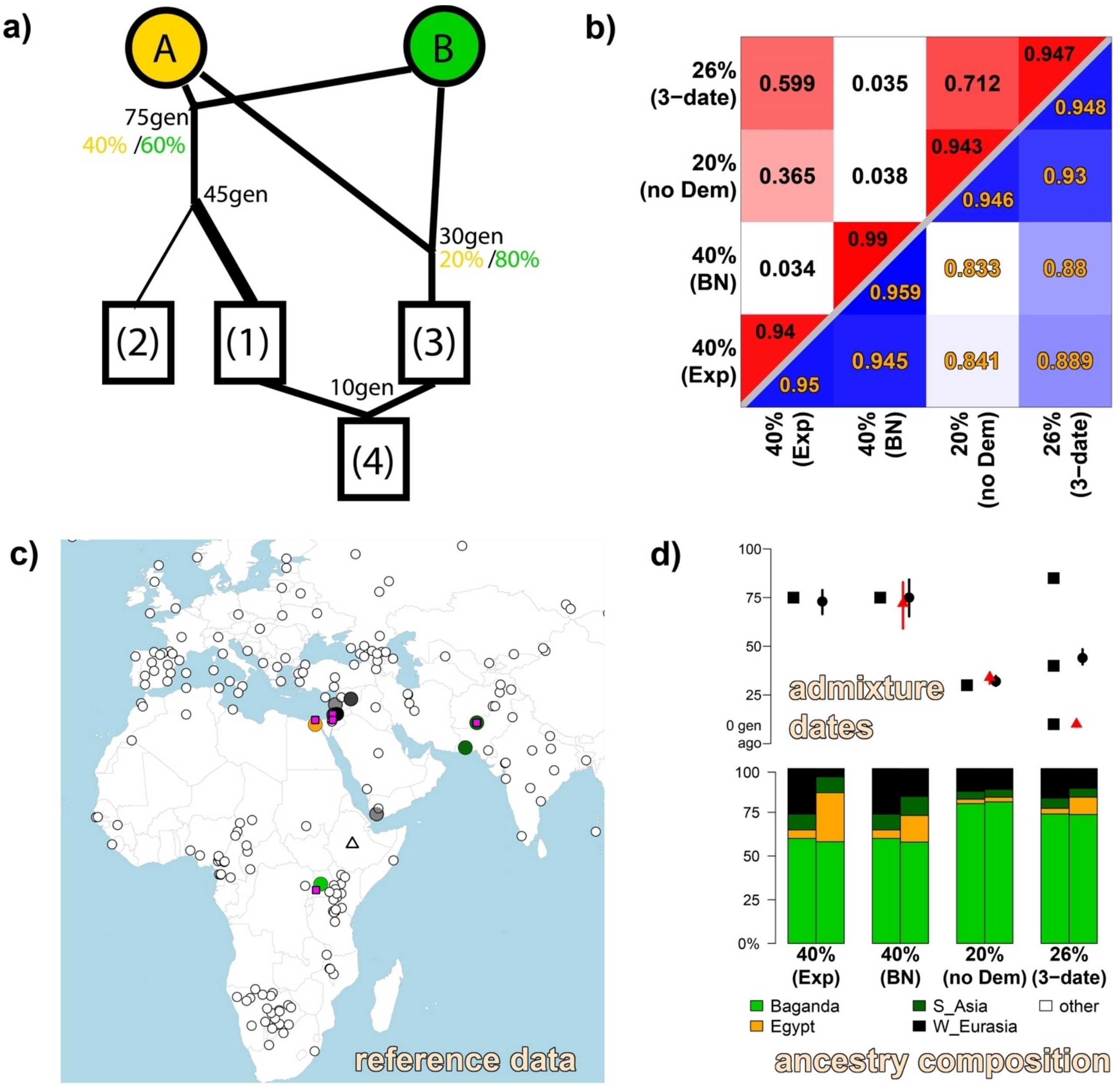


**Figure S5. Simulations. (A)** Schematic of simulations (1)-(4) formed as mixtures of original source populations A and B (see SI section 3). **(B)** Average pairwise genetic similarity (1-TVD) between individuals from different simulated populations, under the "Ethiopia-internal" analogue (top left, red color scale) versus the "Ethiopia-external" analogue (bottom right, blue color scale). **(C)** Non-Ethiopian populations, plus the 4.5kya Ethiopian Mota (Gallego-Llorente et al., 2015), that simulated groups were compared to under the "Ethiopian-external" SOURCEFIND analogue. Open circle populations contributed to describing <1% of the ancestry of any simulated population. Filled circles are darkened according to their maximal SOURCEFIND-inferred contribution to any simulated population relative to that of all other populations from that major geographic region. Pink dots denote the location of the true admixing sources (i.e. comprising sources A and B in Fig S5A). **(D)** Bottom: The true (left bar) and SOURCEFIND-inferred (right bar) ancestry proportions for each simulation. Top: The true (squares) and GLOBETROTTER inferred dates (dot/triangle=mean, line=95% CI) when testing for admixture within each simulation. The black dot and red triangle depict date inference under the “Ethiopia-external” and “Ethiopia-internal” analogues, respectively.

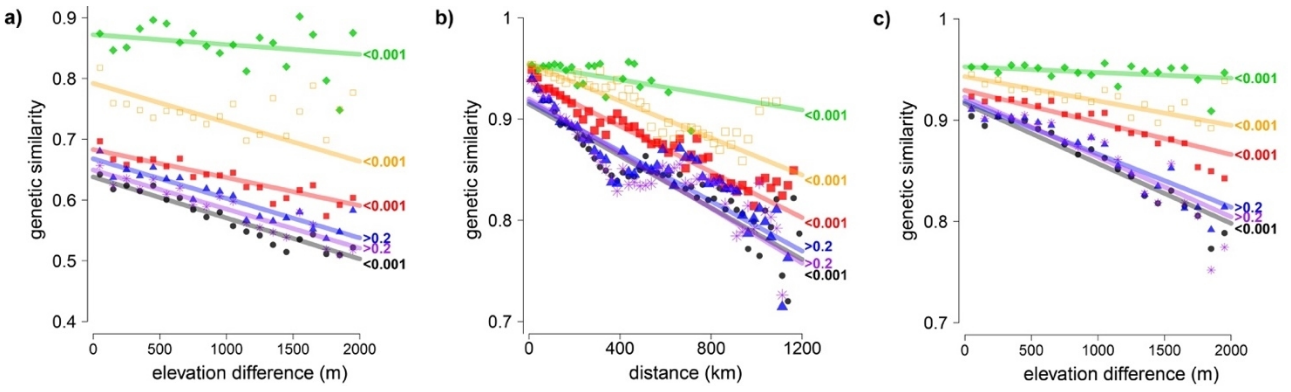


**Figure S6 Genetic similarity correlates with spatial distance and shared cultural factors among Ethiopians**. **(a)** Fitted model for genetic similarity under the “Ethiopia-internal” analysis between pairs of individuals versus elevation difference, with points depicting the average genetic similarity within 100km bins, for all individuals (black; dots) or restricting to individuals who share group label (green; diamonds), speak the same first language (orange; open squares), speak the same second language (blue; triangles), have the same religious affiliation (purple; asterisks), or whose ethnicities are from the same language group (red; closed squares). Labels at right give permutation-based p-values when testing the null hypothesis of no increase in genetic similarity among individuals sharing the given trait (see Methods). **(b)** Analogous fitted model for genetic similarity under the “Ethiopia-external” analysis versus geographic distance, with points depicting averages within 25km bins. **(c)** Analogous fitted model for genetic similarity versus elevation difference under the “Ethiopia-external” analysis, with points depicting averages within 100km bins.

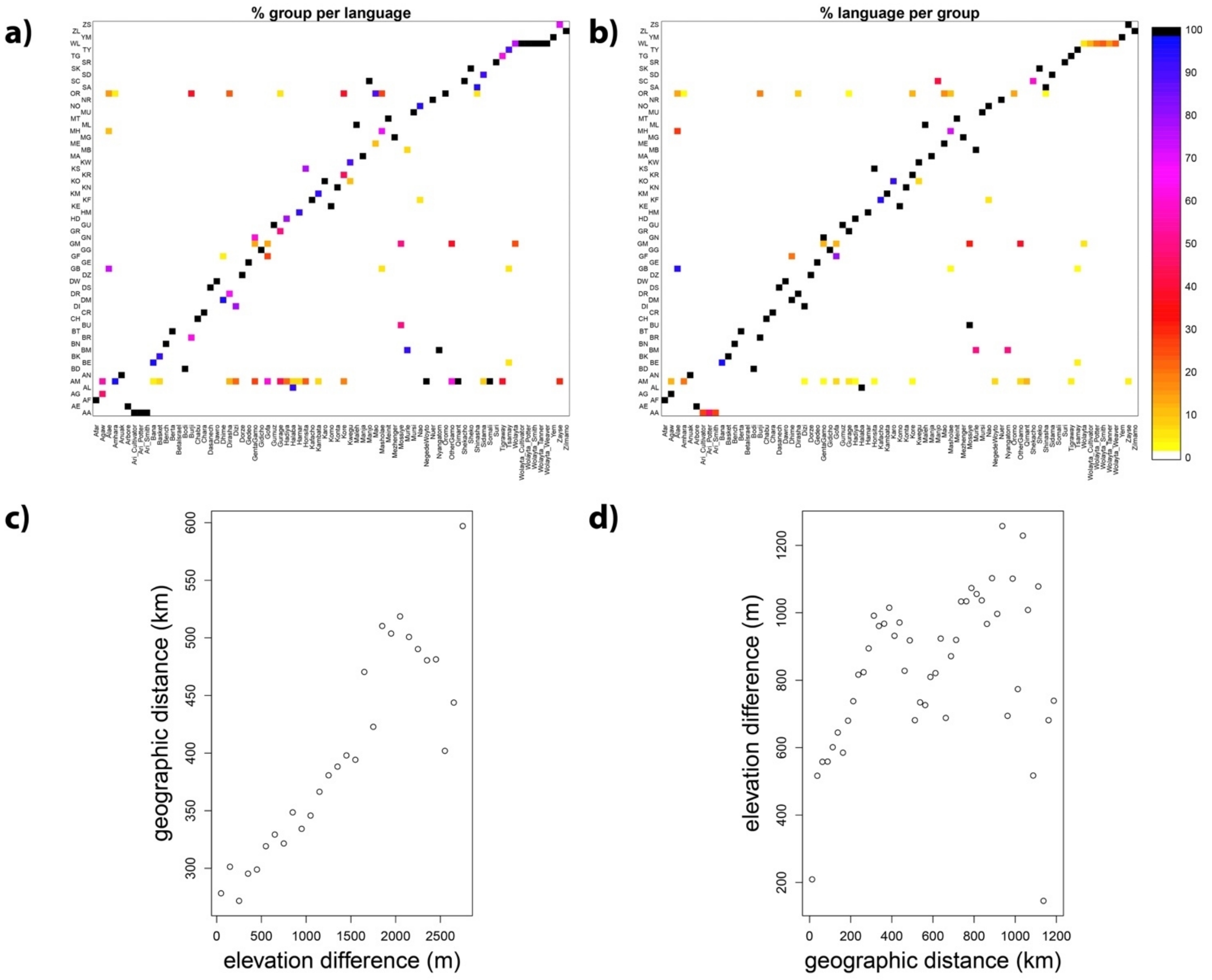


**Figure S7 Correlation between first language and ethnicity, and between geographic and elevation distance. (a)** Percentage of individuals from each group label (column) that speak the given first language (row). **(b)** Percentage of individuals from each first language (row) that fall into each group label (column). I.e. in the left plot, columns sum to 1, while in the right plot, rows sum to 1. The key for each linguistic label is given in Table S1. **(c)** Average geographic distance among individuals within 100-meter bins of elevation difference among individuals. **(d)** Average elevation difference among individuals within 25-kilometer bins of geographic distance.

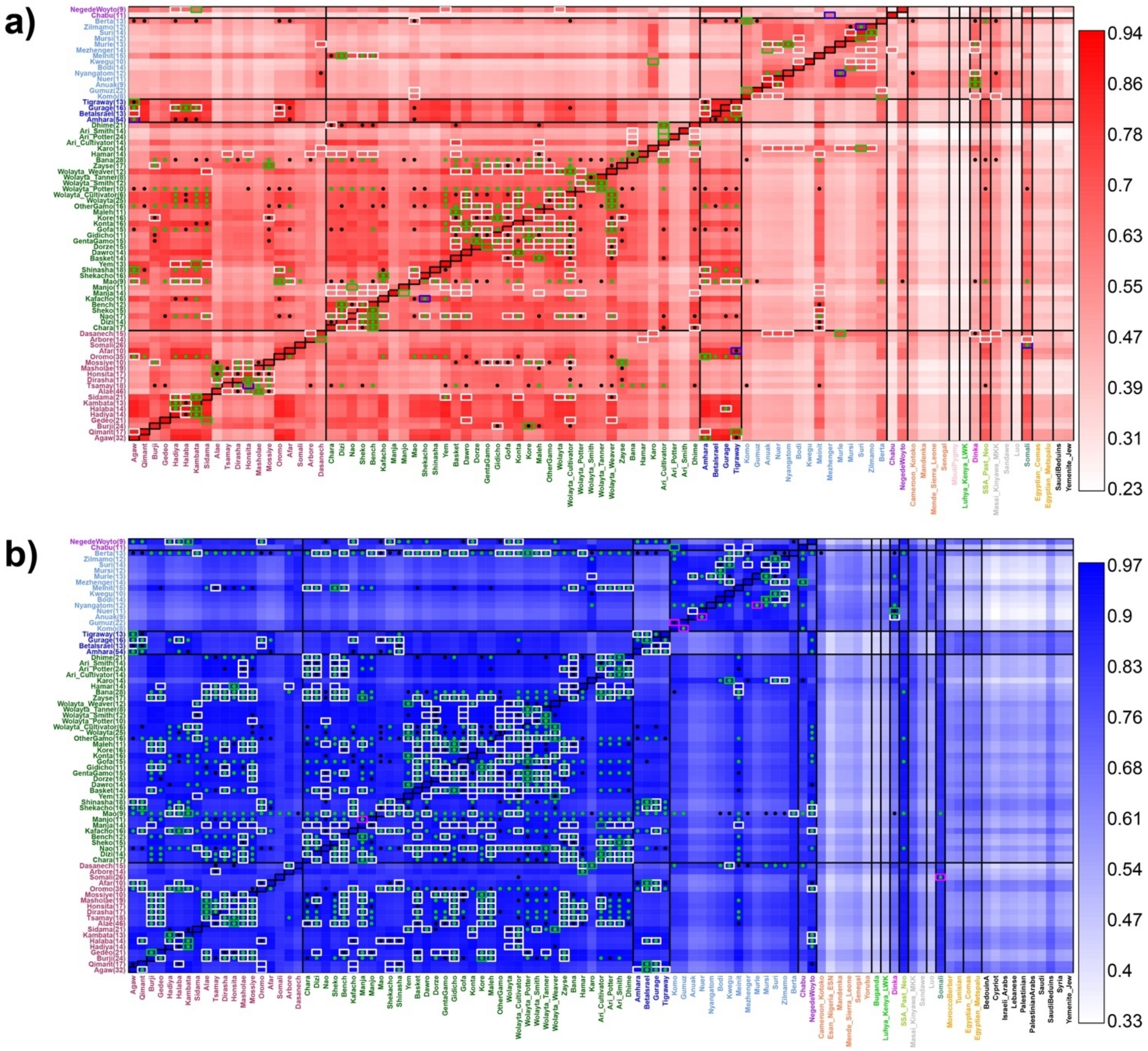


**Figure S8. Average genetic similarity (1-TVD) between each Ethiopian group and other populations under the (a) "Ethiopian-internal" and (b) "Ethiopian-external" analyses.** Each coloured square gives the average genetic similarity (1-TVD; legend at right) between all pairwise comparisons of individuals from the corresponding row, column (values given in Extended Tables 3-4). Columns include all Ethiopian groups, plus all non-Ethiopian groups with >= 7 sampled individuals that had relatively high genetic similarity to at least one Ethiopian group. Within each row *X*, dots denote groups (columns) for which the average genetic similarity between two individuals from group *X* is not significantly higher than that between an individual from *X* and an individual from the column group at Type I error rate = 0.001 (black) or 0.05 (green) (see Methods). Also within each row *X*, colored rectangles enclose: (black) the column for *X*; (green/blue/pink) the group (column) with highest average genetic similarity to *X*; (white) groups whose genetic similarity to *X* is not significantly lower than that between the group with highest genetic similarity to *X* at Type I error rate = 0.001. Blue and pink rectangles in Fig S8a and Fig S8b, respectively, signify that there are no other groups (columns) enclosed in white rectangles for the given row *X*, while green rectangles signify that there are. Ethnic group labels on axes are coloured by language classification for Ethiopian groups (legend in Fig 1a) and by major geographic region for non-Ethiopian groups.

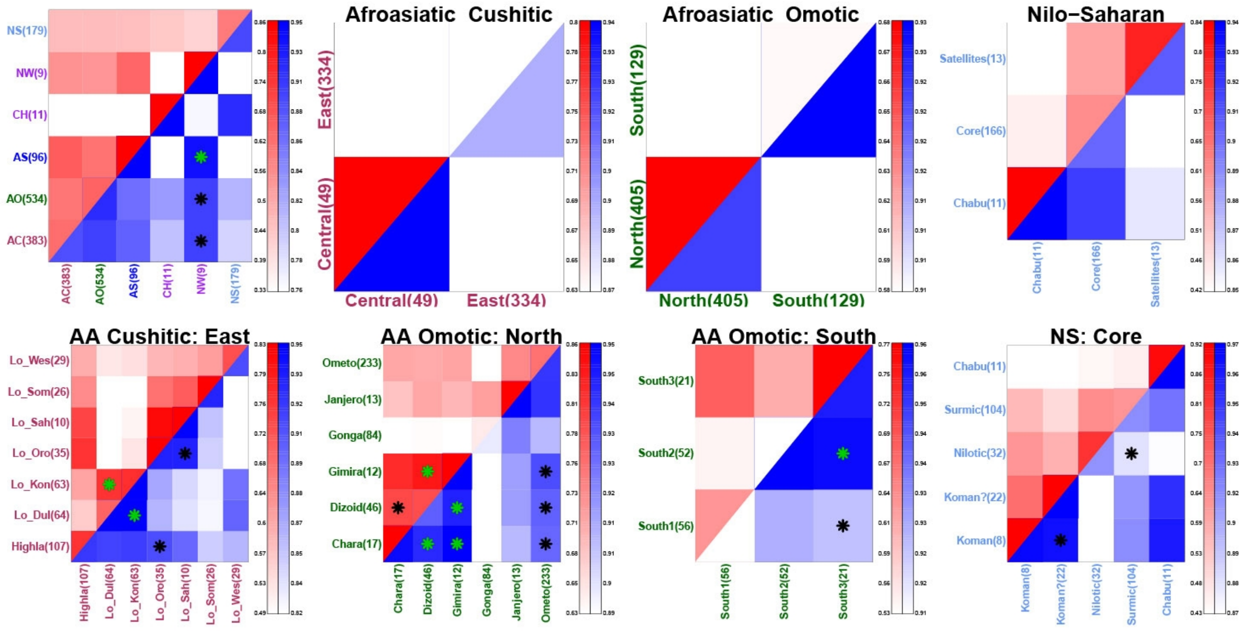


**Figure S9. Average genetic similarity (1-TVD) between Ethiopians whose ethnicities are classified into different language groups.** Row 1 gives results between within-family branches (column 1; second tier linguistic classifications provided by www.ethnologue.com) and between sub-branches (columns 2-4; third tier at www.ethnologue.com). Row 2 gives results for further sub-classifications within sub-branches. Each plot gives results under the "Ethiopia-internal" (upper left triangle, red-scale) and "Ethiopia-external" (lower right triangle, blue-scale) analyses. Each square (row=A, column=B) within a plot gives the average genetic similarity (1-TVD; legend at right of each plot) between all pairwise comparisons of individuals where one individual is from A and the other is from B. Asterisks within each square indicate a lack of significant genetic differentiation between the two classifications at a Type I error level of 0.001 (black) or 0.05 (green), based on 100K permutations individuals' language classifications (see Methods). Numbers in parentheses on the axes give the number of individuals in each language classification, with AC: Afroasiatic Cushitic; AO: Afroasiatic Omotic; AS: Afroasiatic Semitic; NW: Negede-Woyto; NS: Nilo-Saharan; CH: Chabu; Lo_(Dul/Kon/Oro/Sah/Som/Wes): Lowland Dullay/Konso-Gidole/Oromo/Saho-Afar/Somali/Western. "Koman?" refers to the Gumuz. Though the Chabu have no official language family classification, we compared them to Nilo-Saharan sub-family classifications.

**[separate FigS10.pdf file]**

**Figure S10. FineSTRUCTURE inferred clusters of genetically homogeneous Ethiopian groups.** fineSTRUCTURE's best-fitting tree relating its inferred clusters. Each leaf of the tree lists the number of individuals from each labeled group that were assigned to that cluster. Contiguous clusters of the same color were merged into one of the 78 final clusters we used in analysis; we alternate colors here to assist visualisation. Labels for these 78 clusters are provided at right. Full details in Extended Table 2.

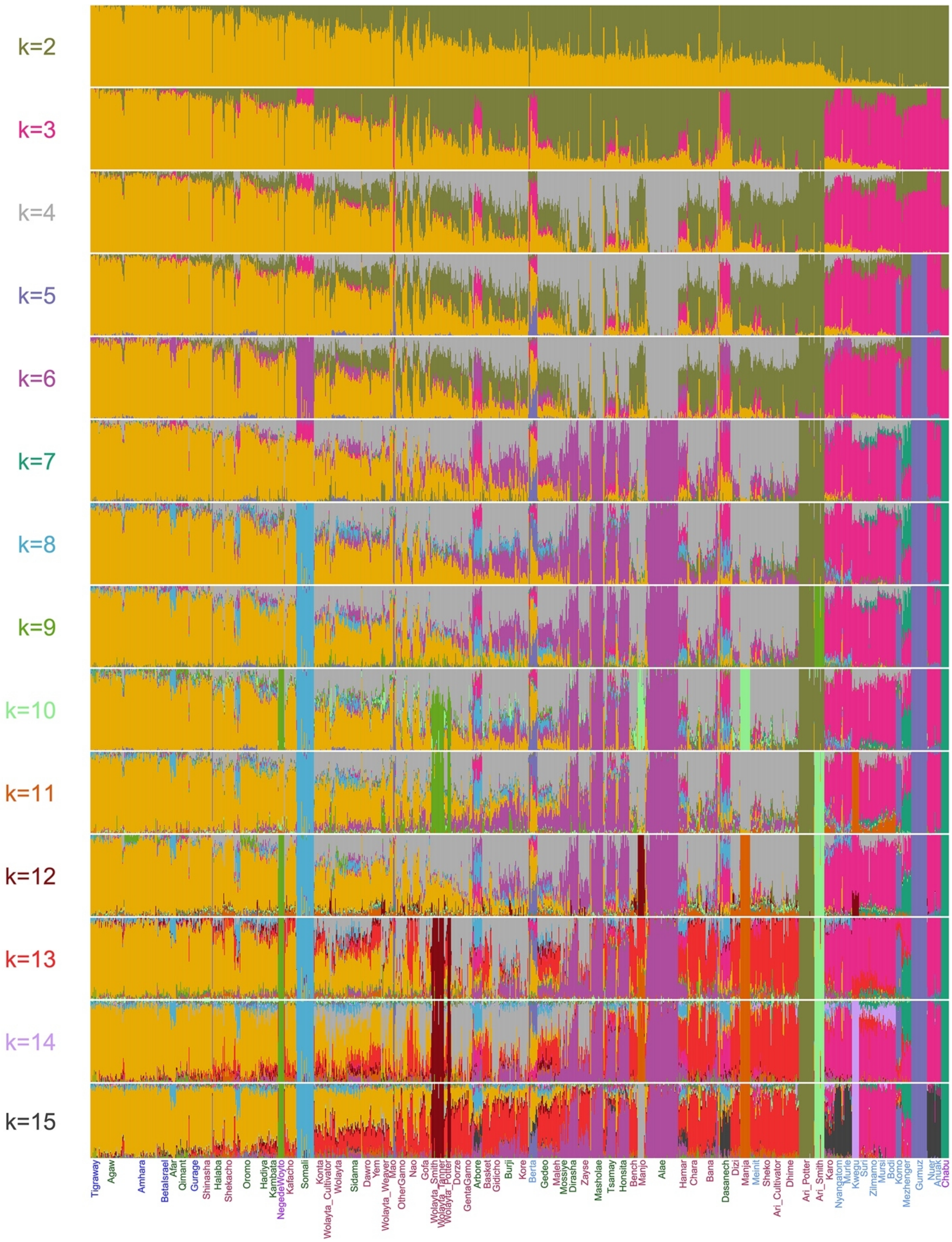
k=2
k=3
k=4
k=5
k=6
k=7
k=8
k=9
k=10
k=11
k=12
k=13
k=14
k=15
Tigraway
Agaw
Amhara
BetaIsrael
Afar
Qimant
Gurage
Shinasha
Halaba
Shekacho
Oromo
Hadiya
Kambata
NegedeWoyto
Kafacho
Somali
Konta
Wolayta_Cultivator
Wolayta
Sidama
Dawro
Yem
Wolayta_Weaver
Mao
OtherGamo
Nao
Gofa
Wolayta_Smith
Wolayta_Tanner
Wolayta_Potter
Dorze
GentaGamo
Arbore
Basket
Gidicho
Burji
Kore
Berta
Gedeo
Majeh
Mossiye
Dirasha
Zayse
Masholae
Tsamay
Honsita
Bench
Manjo
Alae
Hamar
Chara
Bana
Dasanech
Dizi
Manja
Meinit
Sheko
Ari_Cultivator
Dhime
Ari_Potter
Ari_Smith
Karo
Nyangatom
Murle
Kwegu
Suri
Zilmamo
Mursi
Bodi
Komo
Mezhenger
Gumuz
Nuer
Anuak
Chabu

**Figure S11. Inferred clustering of Ethiopians using ADMIXTURE for K=2-15 clusters.** Each bar is an individual, with colors representing clusters. Group labels along the x-axis are colored according to language group (see Fig 1a for key). Labels colors at left denote the new color added in that row.

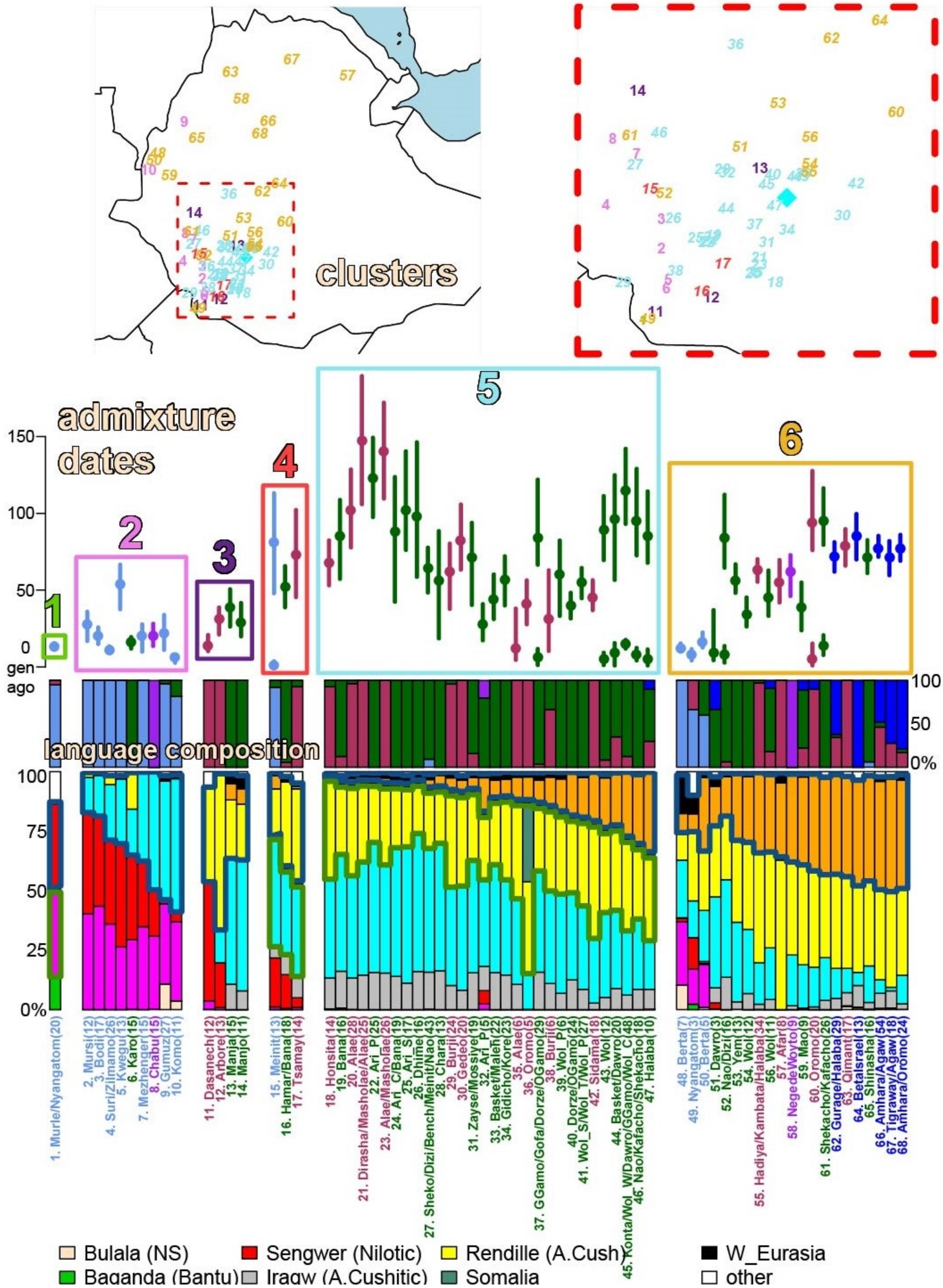
clusters
admixture dates
language composition
1
2
3
4
5
6
150
100
50
0
gen ago
100
50
0%
100
75
50
25
0%
1. Murle/Nyangatom(20)
2. Mursi(12)
3. Bodi(17)
4. Suri/Zilimamo(26)
5. Kwegu(13)
6. Karo(15)
7. Mezhenger(15)
8. Chabu(15)
9. Gumuz(27)
10. Komo(11)
11. Dasanech(12)
12. Arbore(13)
13. Manja(15)
14. Manjo(11)
15. Meinit(13)
16. Hamar/Bana(18)
17. Tsamay(14)
18. Honsita(14)
19. Bana(16)
20. Alae(28)
21. Dirasha/Masholae/Alae(25)
22. Ari_P(25)
23. Alae/Masholae(26)
24. Ari_C/Bana(19)
25. Ari_S(17)
26. Dhime(16)
27. Sheko/Dizi/Bench/Meinit/Nao(43)
28. Chara(13)
29. Burji(24)
30. Gedeo(20)
31. Zayse/Mossiye(19)
32. Ari_P(5)
33. Basket/Malefi(22)
34. Gidicho/Kore(23)
35. Alae(6)
36. Oromo(5)
37. GGamo/Gofa/Dorze/OGamo(29)
38. Burji(6)
39. Wol_P(6)
40. Dorze/OGamo(24)
41. Wol_S/Wol_T/Wol_P(27)
42. Sidama(18)
43. Wol(12)
44. Basket/Dawro(20)
45. Konta/Wol_W/Dawro/GGamo/Wol_C(48)
46. Nao/Kafacho/Shekacho(18)
47. Halaba(10)
48. Berta(7)
49. Nyangatom(3)
50. Berta(5)
51. Dawro(3)
52. Nao/Dizi(16)
53. Yem(13)
54. Wol(12)
55. Hadiya/Kambata/Halaba(34)
56. Wol(11)
57. Afar(8)
58. NegedeWoyto(9)
59. Mao(9)
60. Oromo(20)
61. Shekacho/Kafacho(26)
62. Gurage/Halaba(29)
63. Qimant(17)
64. BetaIsrael(13)
65. Shinasha(16)
66. Amhara/Agaw(54)
67. Tigraway/Agaw(18)
68. Amhara/Oromo(24)
Bulala (NS)
Sengwer (Nilotic)
Rendille (A.Cush)
W_Eurasia
Baganda (Bantu)
Iragw (A.Cushitic)
Somalia
other

**Figure S12. Alternative depiction (to Fig 3 of main text) of inferred ancestral composition and recent admixture events in each Ethiopian cluster. (top, “clusters”:)** FINESTRUCTURE-inferred genetically homogeneous clusters of Ethiopians, with location placed on the map by averaging the latitude/longitude of each cluster’s individuals, colored by which of six types of admixture event (colored labels 1-6 below) that cluster falls into and number corresponding to those at bottom and in Fig 3. The right plot zooms in on the Southern Nation's, Nationalities and Peoples' Region. **(Middle, “admixture dates”:)** Inferred admixture dates in generations from present (dot=mean, line=95% CI), colored by the most-represented language group among ethnicities in that cluster (legend in Fig 1a) and enclosed by the six types of admixture described in Fig 3 legend. **(Middle, “language composition”:)** Barplots give the proportion of individuals from ethnic groups assigned to each language category within each cluster. **(Bottom, “ancestry composition”:)** SOURCEFIND-inferred ancestry proportions for each Ethiopian cluster, with blue and green borders highlighting different admixing sources (see Fig 3 legend). Cluster labels describing ethnic groups contained therein are below each bar, with values in parenthesis giving the cluster sample size.

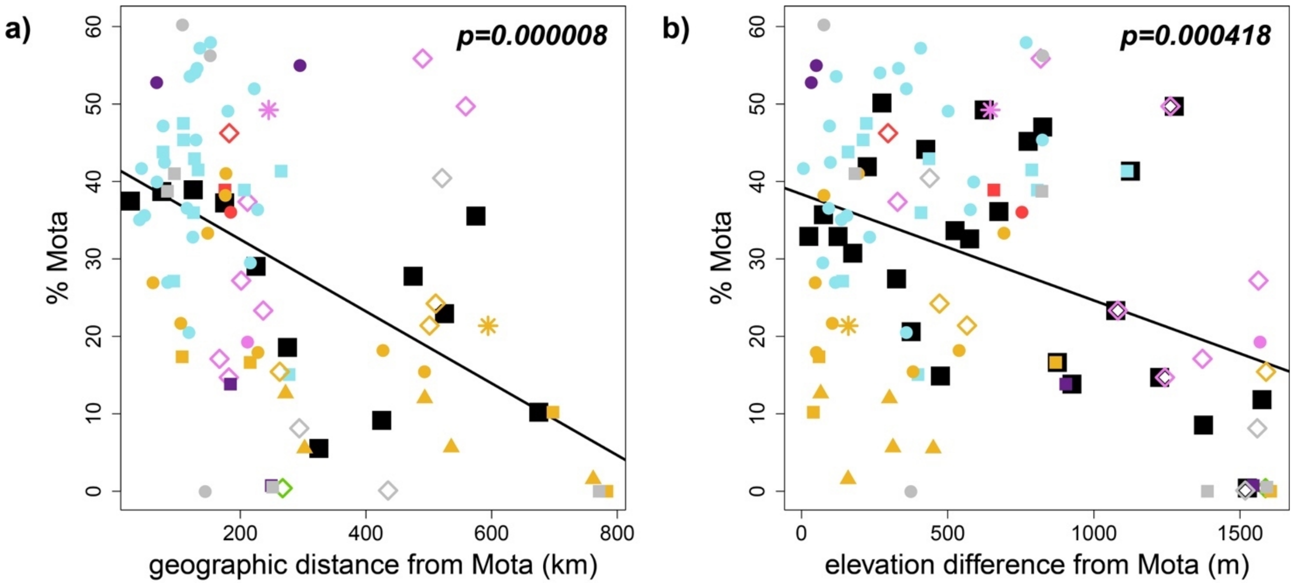


**Figure S13. Matching to Mota decreases with spatial distance.** SOURCEFIND-inferred proportions of DNA matching to Mota versus **(a)** geographic distance or **(b)** elevation difference from Mota for each Ethiopian cluster, with black squares the average proportions within 50 km or m bins and p-value from linear regression at top right. Symbols and colors correspond to language group and admixture type, respectively (see legend in Fig 3); grey dots represent clusters that did not infer admixture events under GLOBETROTTER. Coordinates for Mota are approximated to within 5km of Mota Cave (Gallego-Llorente et al., 2015).

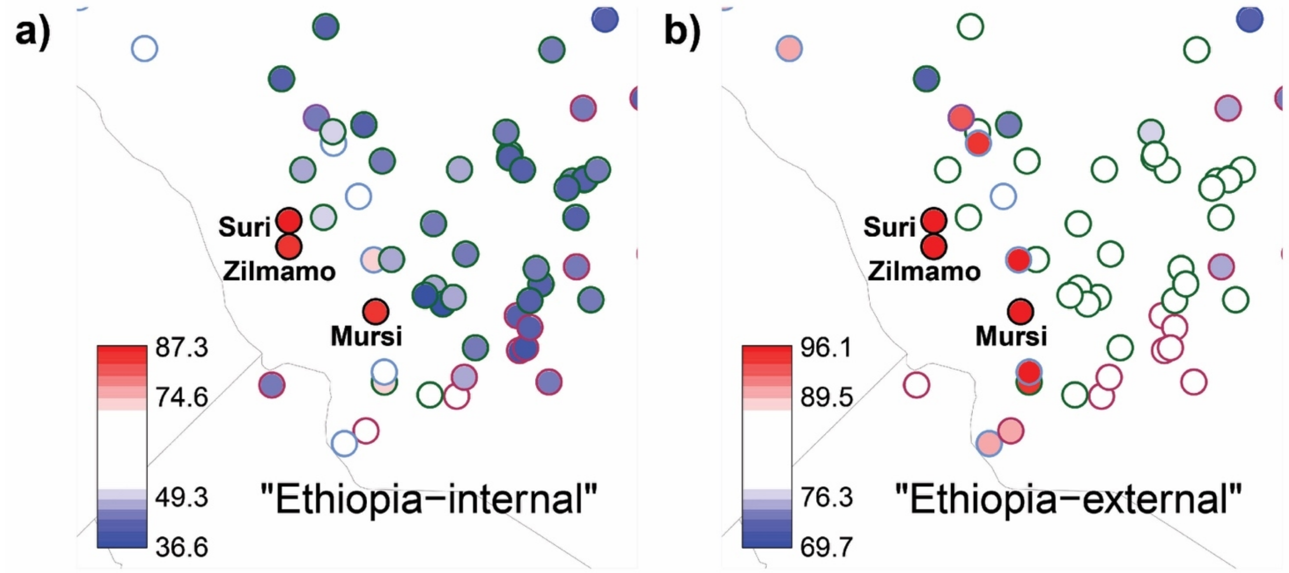


**Figure S14 Average genetic similarity between the NS-speaking Mursi/Suri/Zilmamo** (enclosed with black circles), the only ethnicities in our dataset observed to wear decorative lip plates, and individuals from each labeled group (other circles are colored by the group's language classification as in Fig 1a) under the **(a)** "Ethiopia-internal" and **(b)** "Ethiopia-external" analyses. Circles are placed at the average of each group's individuals' locations, with Suri and Zilmamo slightly shifted (as they have the same such average). Note these three ethnic groups show a relatively high genetic similarity to each other under the "Ethiopia-internal" analysis (Table S12ab), while their similarity is not notably higher than that between these three and other Nilo-Saharan speakers under the "Ethiopia-external" analysis (Table S12cd). These observations are consistent with these three groups' relatively high genetic similarity to one another being primarily attributable to recently separating from one another and/or recently intermixing with each other.

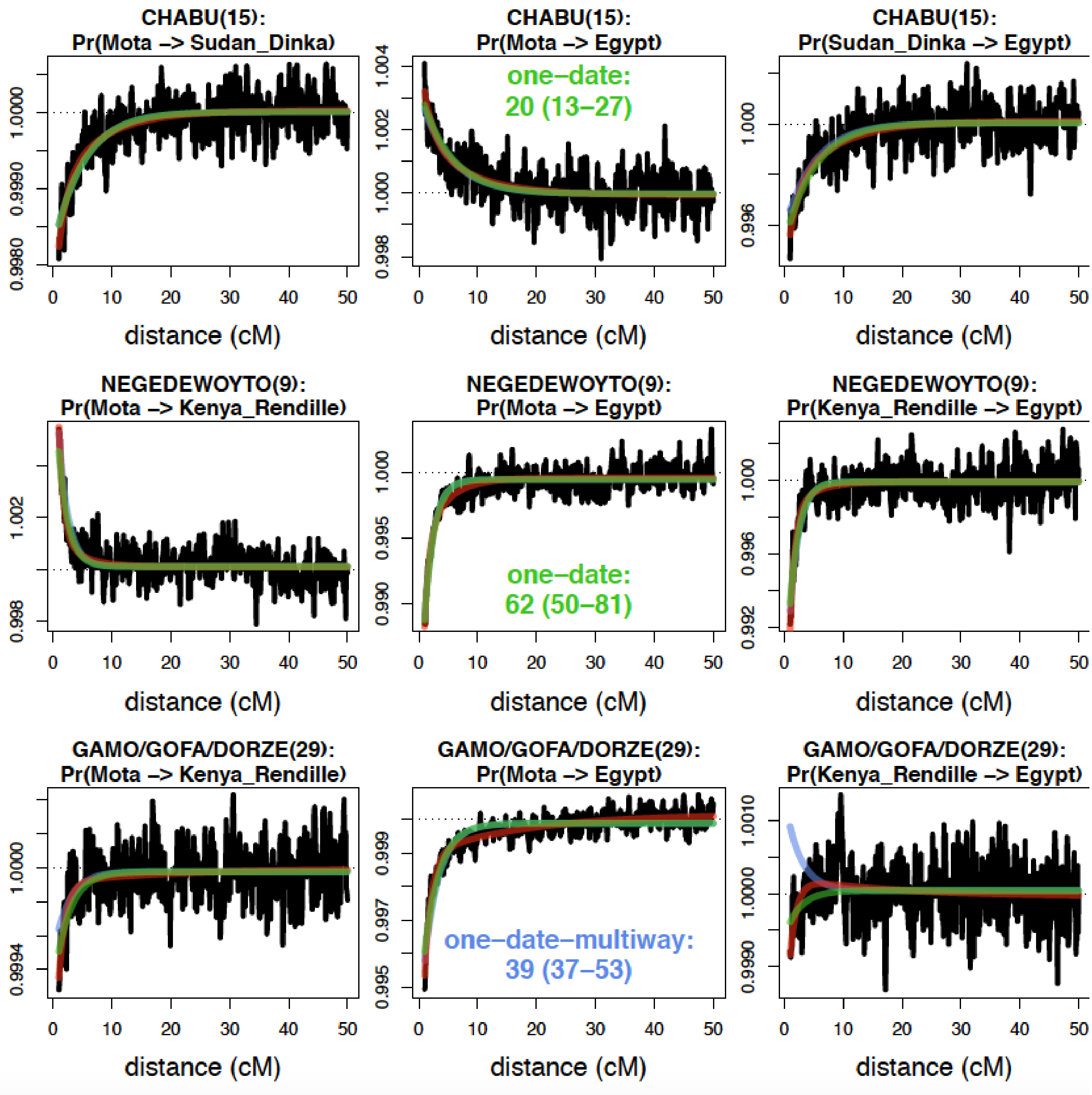


**Figure S15 GLOBETROTTER probability curves testing for admixture in three Ethiopian clusters (rows).** The main ethnic group(s) and total number of individuals within each cluster are given in the title. The black lines in each plot depict how the (scaled) probability of matching two DNA segments within that cluster's individuals to the reference populations listed just below the cluster name changes according to the centimorgan (cM) distance between the two DNA segments. Green lines depict the model fit when assuming a single pulse (date) of admixture, cyan lines when assuming a single pulse of admixture between two sources, and red lines when assuming two pulses of admixture with distinct dates. In the middle plot, we provide the inferred date (in generations ago) and 95% CI.  Curve patterns indicate the number of distinct admixing sources, in that any curves that increase with distance indicate the two reference populations are representing different admixing sources. In contrast, curves that decrease with distance indicate the two reference populations are representing the same admixing source. GLOBETROTTER infers a single admixture event (i.e. “one-date”) between sources related to Egypt/Mota and Sudan_Dinka in the “Chabu” cluster (top row). GLOBETROTTER infers a single admixture event between sources related to Mota/Kenya_Rendille and Egypt in the “Negede_Woyto” cluster (middle row). GLOBETROTTER infers admixture at one date between more than two sources (i.e. “one-date-multiway”) related to Mota, Kenya_Rendille and Egypt in the “Gamo/Gofa/Dorze” cluster (bottom row).

### References


Allentoft, M.E. et al. Population genomics of Bronze Age Eurasia. *Nature* **522**:167-172 (2015).

Alvarez, F. *The Prester John of the Indies: A True Relation of the Lands of the Prester John, being the narrative of the Portuguese Embassy to Ethiopia in 1520* (Beckingham, C. F. and Huntingford, G. W. B. (eds.) Cambridge, 1961).

Ashley, C. Z. Towards a socialised archaeology of ceramics in Great Lakes Africa. *Afr. Archaeol. Rev*. **27**, 135–163 (2010).

Bender, M. L. *Nilo-Saharan Overview In: The Non-Semitic Languages of Ethiopia* (Bender, M. L. (ed) African Studies Center, Michigan State University, Michigan, 1976)

Berry, L. B. *The Solomonic Monarchy at Gonder, 1630-1755: An Institutional Analysis of Kingship in the Christian Kingdom of Ethiopia* (PhD thesis, Boston: Boston University, 1976).

Black, P. *Linguistic Evidence on the Origins of the Konsoid Peoples* (Marcus and Hinnant (eds), Michigan State University, 1975)

Brandt, S. A. New perspectives on the origins of food production in Ethiopia. In: *From Hunters to Farmers. The Causes and Consequences of Food Production in Africa.* (J. D. Clark and S. A. Brandt (eds) Berkeley: University of California Press, 1984).

Broushaki, F. et al. Early Neolithic genomes from the eastern Fertile Crescent. *Science* **353**, 499-503 (2016).

Budge, E. A. W. A Guide to the Babylonian and Assyrian Antiquities. (London: Printed Trustees, 1922).

Castanhoso, D. *Historia das Cousas que o muy Esforçado Capitao Dom Cristovao da Gama fez nos Reinos de Preste Joao com Questrocentos Portugueses que Consigo Levou*, translated in: Whiteway R. S., *The Portuguese Expedition to Abyssinia in 1541-1543 as narrated by Castanhoso* (London 1902, repr. Wiesbaden 1967).

Chacón-Duque, J. C. et al. Latin Americans show wide-spread Converso ancestry and imprint of local Native ancestry on physical appearance. *Nat. Commun.* **9**, 5388 (2018).

Clist, B. A critical reappraisal of the chronological framework of the early Urewe Iron Age industry. *Muntu* **6**, 35–62 (1987)

Crawford, O. G. S. *Ethiopian Itineraries, circa 1400–1524* (Cambridge University Press, 1958).

Currey, J. *Foundations of an African Civilisation: Aksum and the northern Horn* (Oxford, 2014).

Ehret, C. *Cushitic Prehistory, In: The Non-Semitic Languages of Ethiopia* (Benrder, L.M. (ed) African Studies Center, Michigan State University, Michigan, 1976).

Ethiopian Central Statistical Agency. *2007 Ethiopian census, first draft.* (2009).

Gallego-Llorente, M. et al. Ancient Ethiopian genome reveals extensive Eurasian admixture throughout the African continent. *Science* **350**, 820-822 (2015).

Gurdasani, D. et al. The African genome variation project shapes medical genetics in Africa. *Nature* **517**, 327–332 (2015).

Haak, W. et al. Massive migration from the steppe was a source for Indo-European languages in Europe. *Nature* **522**: 207-211 (2015).

Harlan, J. R. Ethiopia: A Center of Diversity. *Economic Botany* **23**, 309-314 (1969).

Hellenthal, G. et al. A genetic atlas of human admixture history. *Science* **343**, 747–751 (2014).

Hofmanova, Z. et al. Early farmers from across Europe directly descended from Neolithic Aegeans. *PNAS* **113**:6686-6891 (2016).

Johanson, D. C. *Lucy (Australopithecus afarensis). In: Evolution: The First Four Billion Years.* (Michael Ruse; Joseph Travis. (eds) Cambridge, Massachusetts: The Belknap Press of Harvard University Press, 2009).

Kalb, J. E. *Adventures in the Bone Trade: The Race to Discover Human Ancestors in Ethiopia's Afar Depression*(New York: Copernicus Books. 2001)

Keller. A. et al. New insights into the Tyrolean Iceman's origin and phenotype as inferred by whole-genome sequencing. *Nature Communications* **3**:698 (2012).

Knibb, M. A. Hebrew and Syriac elements in the Ethiopic version of Ezekiel? *Journal of Semitic Studies* **33**, 11-35 (1988).

Lazaridis, I. et al. Ancient human genomes suggest three ancestral populations for present-day europeans. *Nature* **513**, 409–413 (2014).

Lipson, M. et al. Ancient West African foragers in the context of African population history. *Nature* **577**: 665-670 (2020).

López, S. et al. The Genetic Legacy of Zoroastrianism in Iran and India: Insights into Population Structure, Gene Flow, and Selection. *Am. J. Hum. Genet.* **101**, 353-368 (2017).

Lovejoy, C. O. Re-examining human origins in light of Ardipithecus ramidus. *Science***326**, 74e1:8 (2009).

Malecot, G. Les Voyageurs francajs et les relations entre la France et L'Abyssinle de 1835 à 1870. Outre-Mers. *Revue d'histoire* **212**, 279-353 (1971).

Marcus, H. G. *A History of Ethiopia, Updated Edition* (Berkley, Los Angeles, London: The Regents of the University of California, 2002).

Mathieson, I. et al. Genome-wide patterns of selection in 230 ancient Eurasians. *Nature* **528**:499-503 (2015).

Merid, W. A. *Southern Ethiopia and the Christian Kingdom, 1508-1708, with Special Reference to the Galla Migrations and their Consequences* (Ph.D. Thesis, School of Oriental and African Studies (S.O.A.S.), University of London, 1971).

Muigai, A. W. T. & Hanotte, O. The Origin of African Sheep: Archaeological and Genetic Perspectives. *African Archaeological Review* **30**, 39-50 (2013)

Nersessian, V. & Pankhurst, R. The visit to Ethiopia of Yohannes T'Ovmacean, an Armenian jeweller, in 1764-66. *Journal of Ethiopian Studies* **15**, 79-104 (1982).

Olalde et al. Derived immune and ancestral pigmentation alleles in a 7,000-year-old Mesolithic European. *Nature* **507**, 225-228 (2014).

Pagani, L. et al. Ethiopian Genetic Diversity Reveals Linguistic Stratification and Complex Influences on the Ethiopian Gene Pool. *Am. J. Hum. Genet.* **91**, 83–96 (2012).

Patterson, N. et al. Ancient Admixture in Human History. *Genetics* **192**, 1065-1093 (2012).

Phillips, J. Punt and Aksum: Egypt and the Horn of Africa. *J. African Hist.* **38**, 423-457 (1997).

Phillipson, D. W. *Ancient Ethiopia: Aksum, Its Predecessors and Successors* (London: The British Museum Press, 1998).

Phillipson, D. W. *Foundations of an African Civilisation: Aksum & The Northern Horn 1000 BC – AD 1300* (Suffolk & Rochester, James Currey, 2012).

Phillipson, D. W. *The antiquity of cultivation and herding in Ethiopia. In The Archaeology of Africa: Food, metals and towns* (Thurstan Shaw, Paul Sinclair, Bassey Andah, and Alex Okpoko (eds) London, Routledge, 1993).

Polotsky, H. J. 1964. Aramaic, Syriac, and Ge'ez. *Journal of Semitic Studies* **9**, 1-10 (1964).

Prendergast, M.E. et al. Ancient DNA reveals a multistep spread of the first herders into sub-Saharan Africa. *Science* **365**, eaaw6275 (2019).

Price, A.L et al. Sensitive Detection of Chromosomal Segments of Distinct Ancestry in Admixed Populations. *PLoS Genetics* **5**, e1000519 (2006).

Quintana-Murci, L. et al. Genetic evidence of an early exit of Homo sapiens sapiens from Africa through eastern Africa. *Nat. Genet.* **23**, 437-441 (1999).

Schlebusch, C.M. et al. Southern African ancient genomes estimate modern human divergence to 350,000 to 260,000 years ago. *Science* **358**, 652-655 (2017).

Sergew, H. S. *Ancient and Medieval Ethiopian History to 1270* (Addis Ababa: Artistic Printing Press. 1972).

Shahid, I. The Kebra Nagast in the Light of Recent Research, *Le Museón*, **89** (1976).

Skoglund, P. et al. Reconstructing Prehistoric African Population Structure. *Cell* **171**, 59-71 (2017).

Suwa, G, et al. The Ardipithecus ramidus skull and its implications for hominid origins. *Science*. **326**, 68e1:7 (2009)

Sven, R. *The Survival of Ethiopian Independence* (Heinemann, London, 1978).

Taddesse, T. *Church and State in Ethiopia, 1270-1527* (London: The Clarendon Press, 1972).

The Council of Nationalities (CoN), Southern Nations and Peoples Region (SNNPR), (in Amharic). *A Profile of the Nations, Nationalities and Peoples of the Southern Region, Second Edition* (December 2004EC). Translated from Amharic and sub-edited by Ayele Tarekegn and Neil Bradman, Published by the Council of Nationalities with financial support from the German Co-operation (GIZ), Addis Ababa: Berhanena Selam Printing Enterprise, April 2017 CE.

Council of Nationalities (CoN), Southern Nations, Nationalities and Peoples' Region (SNNPR), (in Amharic). *Yedebubbiheroch, biheresebochna Hizboch Profile* (title translated as *A Profile of the Nations, Nationalities and Peoples of the Southern Region*), Third Edition, Published by the CoN, SNNPR, Addis Ababa: BerhanenaSelam Printing Enterprise, Miazia (April) 2008EC.

Ullendorf, E. 1980. Hebrew, Aramaic and Greek: the versions underlying Ethiopic translations of Bible and intertestamental Literature, In: *The Bible World. Essays in Honor of Cyrus H. Gordon* (G. Rendsburg, R. Adler, M. Arfa and N. H. Winter (eds.), New York, 1980).

Ullendorf, E. *Ethiopia and the Bible—The Schweich Lectures of the British Academy* (London, 1968).

Ullendorf, E. Hebrew elements in the Ethiopic Old Testament. *Jerusalem Studies in Arabic and Islam* **9,** 42–50 (1987).

Ullendorf, E. *The Ethiopians. An Introduction to Country and People* (London and New York, 1960).

Vansina, J. New Linguistic Evidence and 'the Bantu Expansion'. *J. Afr. Hist.* **36**, 173–195 (1995).
