## Supplementary figures and images for "The genetic landscape of Ethiopia: diversity, intermixing and the association with culture"

### FigS10

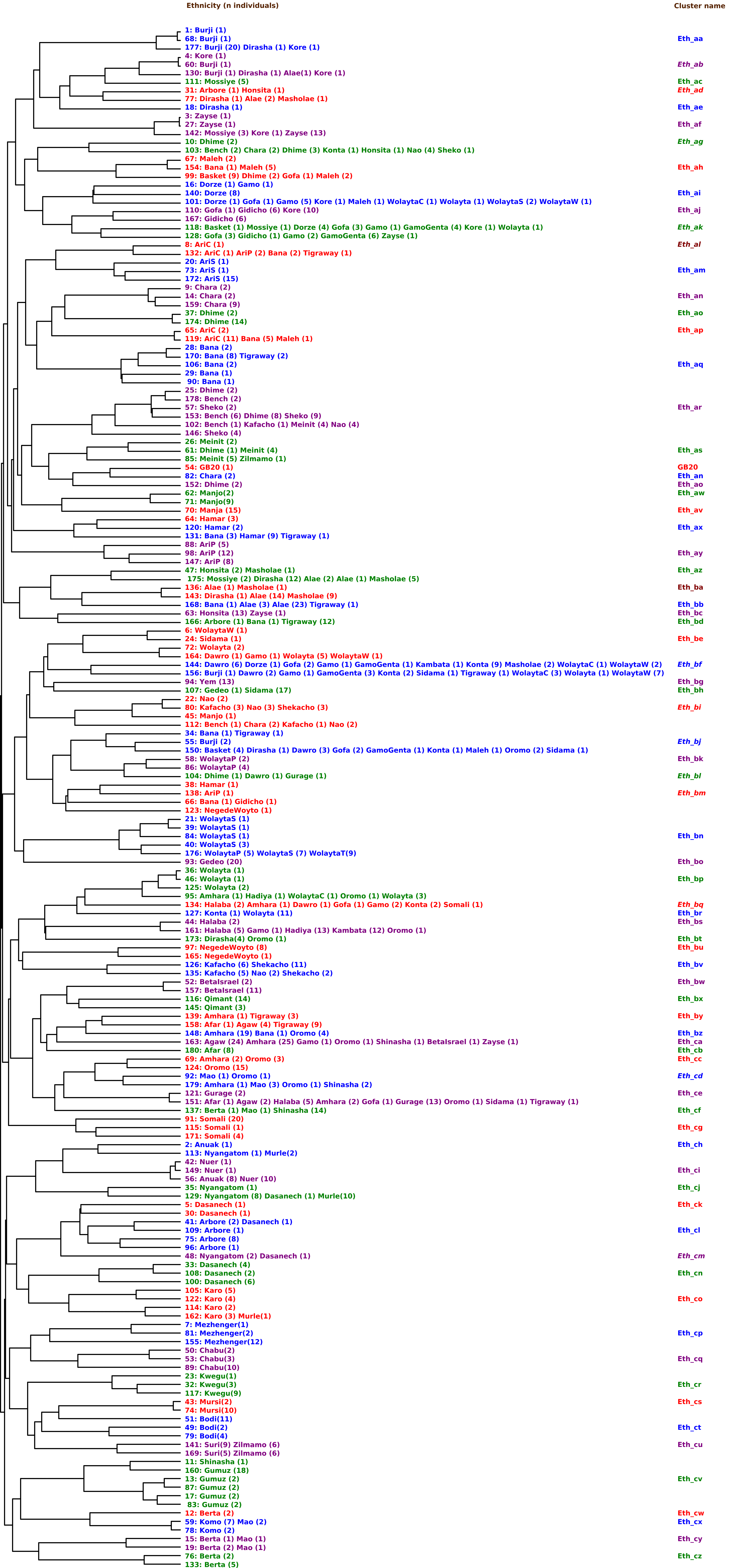
